# Schizophrenia risk gene ZNF804A controls ribosome localization and synaptogenesis in developing human neurons

**DOI:** 10.1101/2024.02.02.578424

**Authors:** Laura Sichlinger, Maximilian Wulf, Lucia Dutan Polit, Fatema Nasser, Rodrigo R.R. Duarte, Timothy R. Powell, Katrin Marcus, Anthony C. Vernon, Deepak P. Srivastava

**Affiliations:** Department of Basic and Clinical Neuroscience, The Maurice Wohl Clinical Neuroscience Institute, Institute of Psychiatry, Psychology, & Neuroscience, King’s College London, London, UK; MRC Centre for Neurodevelopmental Disorders, Institute of Psychiatry, Psychology and Neuroscience, King’s College London, London; United Kingdom; Medizinisches Proteom-Center, Medical Faculty, Ruhr-University Bochum, 44801 Bochum, Germany; Medical Proteome Analysis, Center for Protein Diagnostics (PRODI), Ruhr-University Bochum, 44801 Bochum, Germany; Social, Genetic & Developmental Psychiatry Centre, Institute of Psychiatry, Psychology & Neuroscience, King’s College London, London, United Kingdom

**Author notes:** joint senior authors (Anthony C. Vernon, Deepak P. Srivastava).

## Abstract

*ZNF804A* was amongst the first genes robustly associated with schizophrenia based on findings from large-scale genomic studies. Previous research has implicated *ZNF804A* in the regulation of gene expression and synaptic function, but the role of this gene in neurodevelopment and in schizophrenia pathogenesis remains unclear. To study its function during neurodevelopment, we generated isogenic human induced pluripotent stem cells (hiPSCs) with reduced *ZNF804A* expression, differentiated them into developing cortical glutamatergic neurons and studied their transcriptomic, synaptic and protein signatures. Mutant neurons showed modest evidence changes in gene expression. However, high-content confocal imaging revealed increased excitatory synapse density in mutant neurons. Cell-compartment specific proteomic analysis further revealed that mutant neurons had higher levels of ribosomal and translational proteins within neurites, and high-content imaging confirmed increased local protein synthesis efficiency. Overall, these results demonstrate that in human developing cortical glutamatergic neurons, *ZNF804A* regulates excitatory synapse formation potential via increased local protein translation.

## Introduction

Heritability estimates for schizophrenia (SZ) are among the highest in psychiatric conditions (Sullivan et al., 2012) and large-scale genetic studies have yielded remarkable advances in uncovering the complex polygenic basis of the disorder. It is now understood that both rare genetic variation with high penetrance (Singh et al., 2022; Sriretnakumar et al., 2019) and common single-nucleotide polymorphisms (SNPs) with low penetrance (Trubetskoy et al., 2022) increase susceptibility for SZ. It is however, less well understood how SZ risk loci translate into a neurobiology of the disorder. Evidence from single-cell RNA sequencing (RNAseq) studies suggests that multiple SZ risk genes have enriched expression in glutamatergic neurons present in the developing human cortex (Cameron et al., 2023; Skene et al., 2018). To this end, an increasing number of studies have used human induced pluripotent stem cells (hiPSCs) as a tool to study potential neurodevelopmental mechanisms that may increase susceptibility for SZ (Räsänen et al., 2022). In support of this, hiPSCs generated from individuals with SZ and subsequently differentiated towards neuronal fates show reproducible phenotypes in abnormal neural progenitor cell proliferation, and synaptic disruptions in developing neurons (Brennand et al., 2015; Grunwald et al., 2019; Ishii et al., 2019; Robicsek et al., 2013). While studies using patient-derived hiPSCs have thus enhanced our understanding of the neurobiology of SZ (Räsänen et al., 2022), they have also highlighted the need for functional genomic studies to dissect how specific genes contribute to overall phenotypes. Indeed, a key challenge remains in understanding how specific genetic susceptibility influences neurobiological processes linked with SZ (Murray et al., 2022).

Variations within zinc finger protein 804A (*ZNF804A*) have been repeatedly associated with SZ on a genome-wide scale in large cohorts, establishing it as one of the most robust common risk genes (O’Donovan et al., 2008; Schizophrenia Working Group of the Psychiatric Genomics, 2014; Trubetskoy et al., 2022). Neuropimaging and cognitive studies in both healthy individuals and patients with schizophrenia have associated variants in ZNF804A with SZ-relevant endophenotypes (Chang et al., 2017; Esslinger et al., 2009; Lencz et al., 2010; Rasetti et al., 2011). However, what is less clear is what role this gene plays at the cellular and molecular level. Functional studies using SH-SY5Y and human neural progenitor cell models have suggested that ZNF804A has a role in controlling gene transcription (Chapman et al., 2018; Chen et al., 2015; Hill et al., 2012). In mature rodent neurons, ZNF804A has been shown to localize to synapses (Deans et al., 2017), where it has been implicated in the maintenance of dendritic spines, the site where the majority of excitatory synapses occur (Deans et al., 2017; Dong et al., 2020; Zhou et al., 2020). In addition, knockdown of ZNF804A attenuated structural remodeling of dendritic spines in response to activity-dependent stimulation (Deans et al., 2017). Consistent with these findings, a mutant *ZFP804A* (rodent homologue) mouse model has been described to display reduced dendritic spine density and impaired contextual fear and spatial memory (Huang et al., 2020). ZNF804A has also been suggested to directly interact with cell-adhesion, translation-related and RNA-binding proteins (Aabdien et al., 2024; Dong et al., 2020; Klockmeier et al., 2021; Zhou et al., 2018), and to regulate translational efficiency in rodent CAD cells (Zhou et al., 2018). While these studies have begun to uncover the molecular and cellular role of ZNF804A, the majority of them have used mature rodent neurons. This is of note as *ZNF804A* shows highest expression during fetal development, when dysfunction of this gene is thought to contribute to the pathogenesis of SZ (Hill & Bray, 2012; Tao et al., 2014). It is therefore necessary to study the role of ZNF804A during neurodevelopment to gain a better understanding of how variations in this gene may contribute to SZ.

In this study, we explored the role of *ZNF804A* in developing human neurons by employing isogenic mutant hiPSC cell lines with reduced ZNF804A expression, generated by Clustered Regularly Interspaced Short Palindromic Repeats (CRISPR)/Cas9 genome engineering. Initial analysis of single-cell transcriptomic data of developing human cortex identified fetal cortical neurons to express *ZNF804A* at the highest level. Following this, mutant isogenic hiPSC lines were differentiated into developing glutamatergic cortical neurons for assessment. Bulk RNAseq only revealed only minor differences in gene expression in mutant *ZNF804A* neurons, suggesting that transcriptional regulation might not be a primary function of ZNF804A in this cell type. However, genes with altered expression were enriched for cell adhesion and synaptic functioning, aligning partially with prior studies (Chapman et al., 2018; Hill et al., 2012). High-content confocal imaging revealed an increase in pre- and post-synaptic protein levels and a higher putative synapse density along MAP2-positive (MAP2^+^) neurites of mutant glutamatergic neurons, indicating a role for ZNF804A in regulating synaptogenesis in developing neurons. Proteomic analysis further revealed changes in the distribution of translation machinery and translation regulation proteins between neurites and cell bodies in ZNF804A mutant neurons. In line with these observations, we observed elevated local protein synthesis efficiency and increased expression of ribosomal protein S6 (RPS6) along MAP2^+^ neurites in ZNF804A mutant neurons. These findings provides insight into ZNF804A’s role in developing glutamatergic neurons, and indicate that dysfunction of this gene could result in altered synaptogenesis and regulation of local protein synthesis during neurodevelopment.

## Results

### Developing glutamatergic cortical neurons are a suitable model to study ZNF804A function

Previous studies have shown that *ZNF804A* is highly expressed during the second trimester of human cortical neurodevelopment and that full length and a truncated form of ZNF804A, *ZNF804A^E3E4^* is associated with risk for SZ (Hill & Bray, 2012; Tao et al., 2014). However, we lack information on the cell type and isoform expression profile of this gene in the developing cortex. We therefore first extracted gene specificity scores for *ZNF804A* expression from published single-nuclei RNAseq data obtained from the frontal cortices of second trimester human fetuses (Cameron et al., 2023). This revealed that *ZNF804A* transcripts were broadly distributed across all identified cell populations, with an enrichment in neuronal cells, particularly within developing upper layer excitatory cortical neurons (specificity score 31%; **Fig. 1A**). Next, we investigated the isoform expression of ZNF804A across six different microstates of cortical neural progenitor cell (NPC) development by differentiating three clones from three male neurotypical donor lines towards a cortical linage (**fig. S1**) (Adhya et al., 2021). We specifically assessed the expression of the SZ risk variant *ZNF804A^E3E4^* and full-length transcripts at six different timepoints. (**table S1**) Three-way ANOVA showed that developmental timepoint (F(6) = 111.3, p < 0.0001, η^2^ = 0.786), isoform (F(1) = 15.2, p < 0.001, η^2^ = 0.018), and individual genetic make-up (i.e. different donor lines) (F(2) = 27.7, p < 0.00001, η^2^ = 0.065) had a statistically significant main effect on transcript abundance (**table S2 and Fig. 1B**). Tukey *post-hoc* analysis revealed that *ZNF804A^E3E4^* expression was significantly lower than the full-length variant throughout NPC development (p.adj = 0.0002). Moreover, Tukey *post-hoc* analysis showed significant differences when comparing almost all time-points with higher expression of both isoforms at later stages of development (**table S2; Fig. 1C**). Comparisons of later developmental stages however, did not yield statistically significant results (D14 vs. D16 (p.adj = 0.15); D16 vs. D18 (p.adj = 0.12), and D18 vs. D20 (p.adj = 0.99), **table S2; Fig. 1C**). These data suggest that *ZNF804A* isoform expression gradually increases throughout cortical NPC development, reaching its highest levels in developing glutamatergic neurons.

**Fig. 1.**
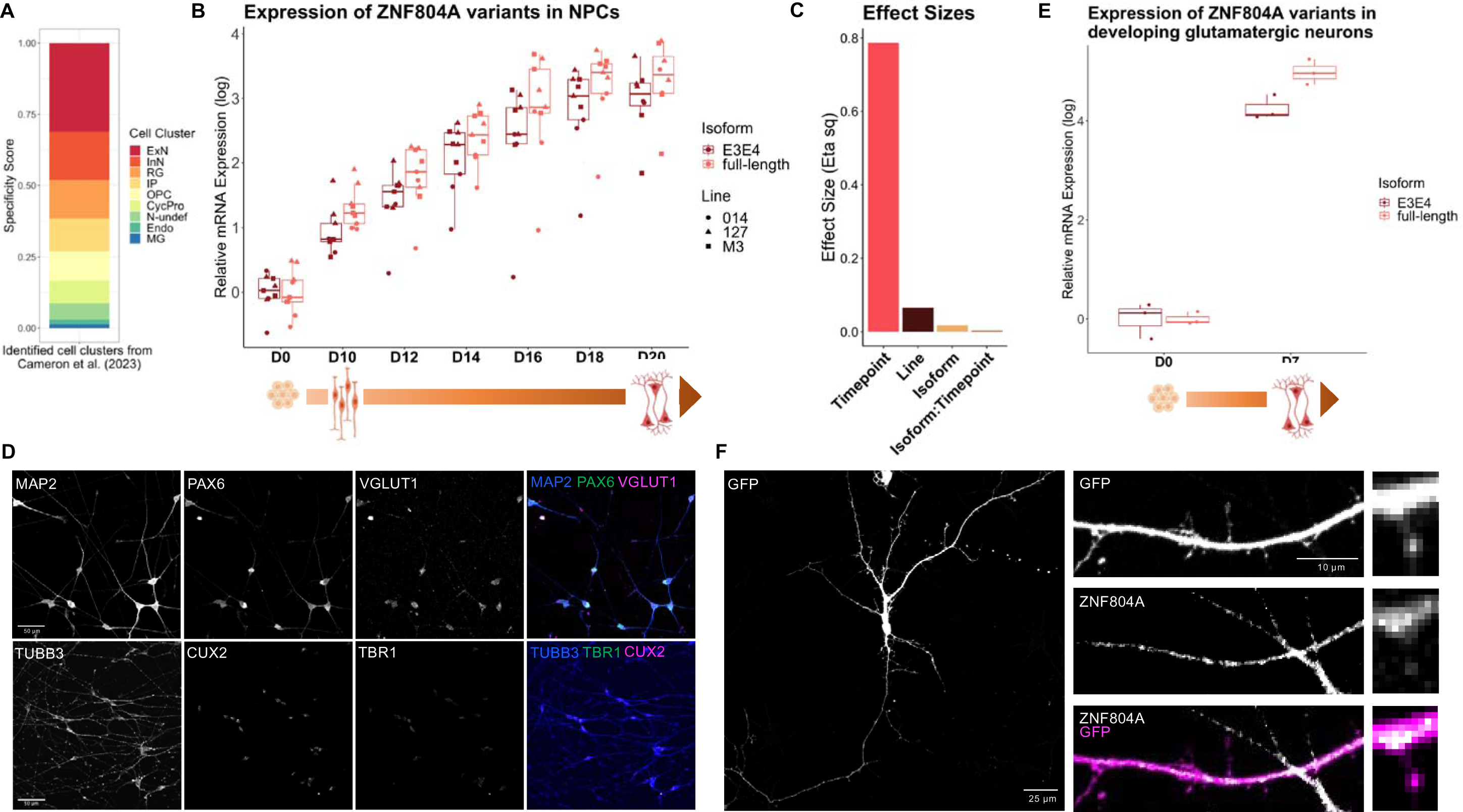
Expression profile of ZNF804A isoform transcripts across glutamatergic neuron development. (A) Stacked barchart of gene specificity scores for ZNF804A in different cell population clusters of the foetal cortex. Annotations: ExN = developing excitatory neuron; InN = developing inhibitory neuron; N-undef = neuron of undefined class; IP = intermediate progenitor; RG = radial glia; CycPro = cycling progenitor cells; OPC = oligodendrocyte precursor cell; Endo = endothelial cell; MG = microglia. (B) Boxplots showing ZNF804A^full-length^ and ZNF804A^E3E4^ expression at D0-D20 timepoints reflecting microstates of NPC development. Overlaying dotplots show values for each cell line. Three-way ANOVA revealed significant results for timepoint (F(6) = 111.3, p < 0.0001), isoform (F(1) = 15.2, p < 0.001) and line (F(2) = 27.7, p < 0.00001). (C) Effect sizes (η^2^) of three-way ANOVA. (D) Representative confocal image of D7 glutamatergic forebrain neurons showing immunostaining for pan-neuronal marker microtubule-associated protein 2 (MAP2; blue), excitatory presynaptic marker VGLUT1 (magenta) and Pax6 (green) (upper images); pan-neuronal early-stage marker class III beta-tubulin (TUBB3; blue), cortical layer VI-specific marker TBR1 (green) and upper layer-specific marker Cux2 (magenta) (lower images). (E) Boxplots showing ZNF804A^full-length^ and ZNF804A^E3E4^ expression in D7 forebrain neurons. Overlaying dotplot shows values for each biological replicate. Student’s t-test revealed significant reductions for ZNF804A^E3E4^ at D7 (t(4) = −3.58, p < 0.05) (F) Representative confocal image of green fluorescent protein (GFP)-expressing D7 neuron demonstrating ZNF804A immunostaining along dendrites and in the shaft of putative dendritic spines.

We next studied ZNF804A isoform expression in developing glutamatergic cortical neurons by employing a forward programming approach (Pavlinek et al., 2022; Pawlowski et al., 2017). To avoid heterogeneity in the population of glutamatergic cell type generated by this approach (Treutlein et al., 2016), *NGN2* overexpression was combined with a directed differentiation approach utilizing SMAD and Wnt inhibition (Nehme et al., 2018). This approach has been reported to yield developing upper layer glutamatergic cortical neurons, a cell population shown to exhibit an enrichment of schizophrenia risk variants (Cameron et al., 2023) and pronounced *ZNF804A* gene expression (**Fig. 1A**). At day 7 (D7) of differentiation following this differentiation protocol, glutamatergic neurons were confirmed to express upper layer cell identity markers including cut like homeobox 2 (*CUX2*) transcripts (**Fig. S2**) and protein (**Fig. 1D**) using RT-qPCR and confocal microscopy, respectively. Furthermore, neurons highly expressed Vesicular Glutamate Transporter 1 (VGLUT1), confirming their glutamatergic fate (**Fig. 1D**). When we examined *ZNF804A* isoform expression, a substantial increase in transcript abundance for both isoforms was observed at D7. However, expression levels of the *ZNF804A^E3E4^* transcript were significantly lower than full-length *ZNF804A* ([mean ± SD] *ZNF804A^E3E4^* [4.25 ± 0.25] vs. *ZNF804A^full-length^* [4.98 ± 0.26], t(4) = −3.58, p < 0.05), mirroring the expression profile observed across the development of NPCs (**Fig. 1E**). Finally, we confirmed ZNF804A protein expression in D7 glutamatergic neurons by confocal microscopy, with protein expression observed along neurites and at the base of putative dendritic spines (**Fig. 1F).** Together, these data demonstrate that developing glutamatergic cortical neurons have the highest expression levels of *ZNF804A* transcripts and thus is a suitable cell-type and time-point for studying ZNF804A neurobiological function.

### Generation of isogenic mutant ZNF804A hiPSCs for functional genomic studies

To study the function of ZNF804A, we used a dual-single guide RNA (sgRNA) CRISPR/Cas9 approach to disrupt the function of the gene. Specifically, we targeted exon 3 for complete ablation due to its presence in all known *ZNF804A* transcripts. SgRNAs were designed to induce double-strand breaks within intron 2 (sgRNA1) and intron 3 (sgRNA2), resulting in a large genomic excision of 1783 base pairs, including exon 3 (**Fig. 2A**) – this was predicted to induce frame shifts and premature stop codons in exon 4 upon its deletion (**fig. S3**). Six single-cell isogenic clones – two for each genotype – were chosen for subsequent analysis: 2x clones containing homozygous excisions (hom; ZNF804A^−/−^ clones (#20 and #44)), 2x clones containing heterozygous excisions (het; ZNF804A^+/−^ clones (#10.23 and #42.13)), and 2x clones with no edits to exon 3 (wildtype (wt); ZNF804A^+/+^ (clones #4 and #61)) (**Fig. 2B**). Sanger sequencing confirmed deletion of 1783bp for ZNF804A^−/−^ and ZNF804A^−/+^ mutant clones, along with identical cut sites (chr2: +184933496 & +184935280). ZNF804A^+/+^ clones showed unaltered sequences up- and downstream of each putative Cas9 cut site (**fig. S4**). *ZNF804A* transcript and protein expression was subsequently assessed in D7 developing glutamatergic neurons. RT-qPCR confirmed significant differences in mRNA abundance of full-length (Kruskal-Wallis: *χ*^2^(2) = 14.79, *p* < 0.001) and *ZNF804A^E3E4^* (Kruskal-Wallis: *X*^2^(2) = 15.16, *p* < 0.001) across all groups (**Fig. 2C & fig. S5**). Although reductions in mRNA abundance were observed in both heterozygous and homozygous clones for both isoforms, statistical significance was primarily attributable to differences between ZNF804A^+/+^ and ZNF804A^−/−^ clones (full-length: p.adj < 0.01, *ZNF804A^E3E4^*: p.adj < 0.001), as indicated by Dunn’s *post-hoc* tests with Holm corrections.

**Fig. 2.**
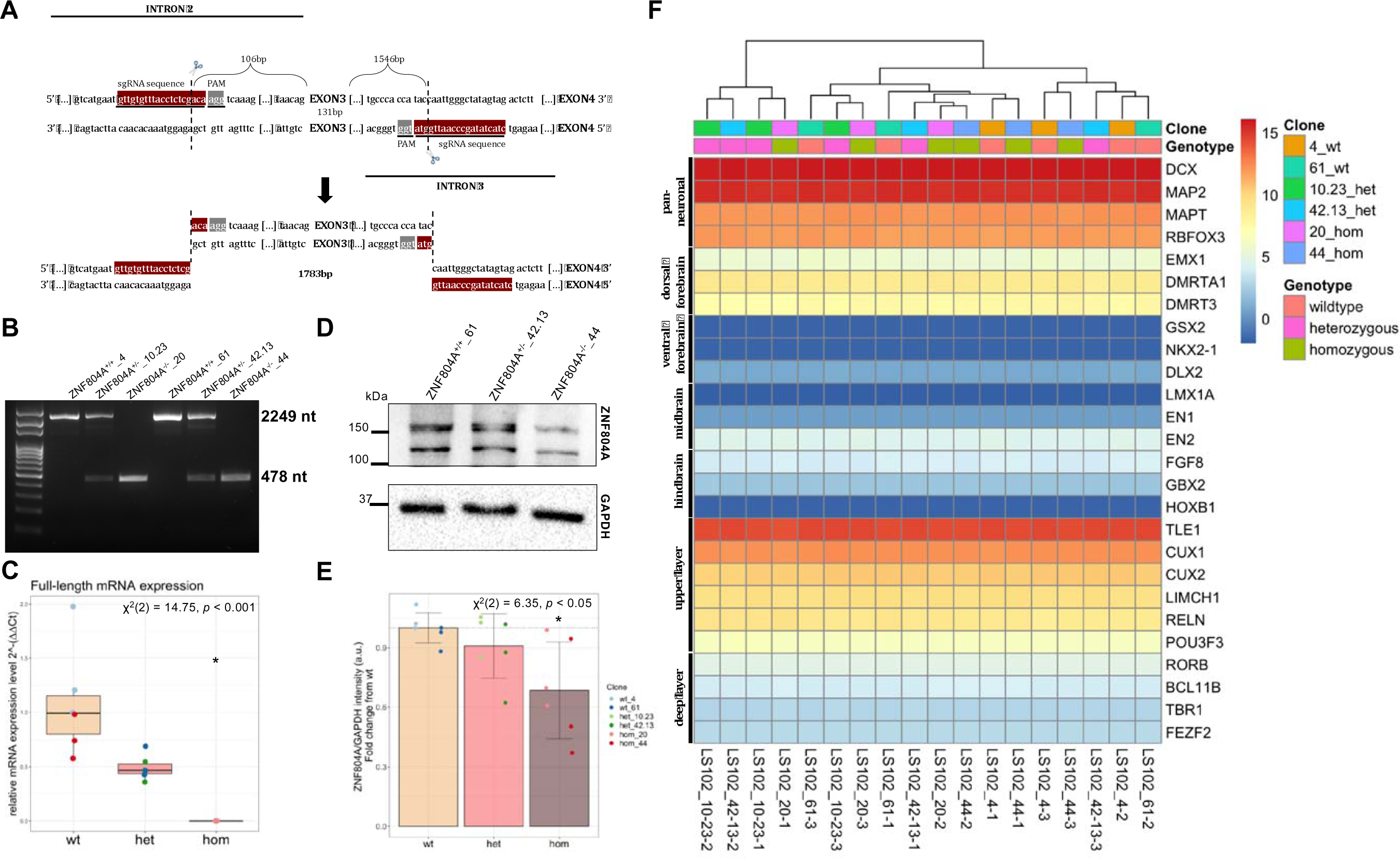
Generation of mutant ZNF804A hiPSC model. (A) Schematic of excision site (1783 base pairs (bp)) of dual single guide RNA (sgRNA) approach; sgRNAs induced double strand breaks within either intron 2 (sgRNA1) or intron 3 (sgRNA2). (B) Representative PCR blot of ZNF804A^+/+^, ZNF804A^+/−^, and ZNF804A^−/−^ clones. Primers spanned the excision site amplifying either 2249 nucleotide (nt; ZNF804A^+/+^, ZNF804A^+/−^) or 478 nt (ZNF804A^−/−^, ZNF804A^+/−^) sequences. Clone ids: wildtype: ZNF804A^+/+^_4, ZNF804A^+/+^_61; heterozygous: ZNF804A^+/−^_10.23, ZNF804A^+/−^_42.13; homozygous: ZNF804A^−/−^_44, ZNF804A^−/−^_20. (C) Boxplots of ZNF804A^full-length^ expression in D7 neurons derived from ZNF804A^+/+^, ZNF804A^+/−^, and ZNF804A^−/−^ lines (n = 18; 3 differentiations per clone). Expression in individual clones is depicted in overlapping dotplots. * p.adj. < 0.05. (D) Representative western blot illustrating decrease in ZNF804A protein levels in ZNF804A^−/−^, as detected by C-terminal binding ZNF804A antibody C2C3. GAPDH was used as a loading control. (E) Bar chart displaying the mean integrated density quantification of two ZNF804A C2C3 bands between 100-150kDa. Overlapping dot plot illustrate ZNF804A C2C3 expression for each clone. Statistical analysis using a Kruskal-Wallis test (*X*^2^(2) = 6.35, p = 0.042) followed by Dunn’s post-hoc tests with Holm corrections revealed a significant reduction of protein in ZNF804A^−/−^ mutants when compared to the ZNF804A^+/+^. * p.adj. < 0.05. (F) Heatmap listing hallmark genes associated with different parts of the cortex and their expression levels in our D7 glutamatergic neurons derived from control and mutant hiPSC lines. Transcriptomic signatures in cells closely resemble developing glutamatergic dorsal forebrain neurons as they mature into upper-layer cortical cells. Log-transformed expression of DESeq2 normalized counts of markers associated with specific cell types is plotted using colour key scaled from 15 to 0 with 15 (red) representing high gene expression and 0 (dark blue) representing low gene expression. There is no discernible clustering based on clone or genotype. List of genes is provided in **table S3**.

To evaluate protein expression, we conducted immunoblotting using a previously validated antibody targeting the C-terminus of ZNF804A (ZNF804A C2C3) (Deans et al., 2017), and quantifying the integrated density of two bands in the 100-150kDa range (**Fig. 2D**). While protein expression appeared unaffected in ZNF804A^+/−^ mutants, a significant reduction was observed in ZNF804A^−/−^ mutants when compared to ZNF804A^+/+^ controls (Kruskal-Wallis test: *X*^2^(2) = 6.35, p = 0.042; **Fig. 2D**). This was confirmed by Dunn’s *post-hoc* test with Holm corrections, which indicated statistical significance in the comparison between ZNF804A^−/−^ and ZNF804A^+/+^ mutants (wt vs. hom p.adj. < 0.05), while differences between ZNF804A^+/−^ and ZNF804A^+/+^ mutants (wt vs. het p.adj. > 0.05) and between heterozygous and homozygous mutants (het vs. hom p.adj. > 0.05) were not significant. On average, we observed an approximate 35% reduction in signal intensity in ZNF804A^−/−^_20 and a 45% reduction in ZNF804A^−/−^_44 clones in comparison to the ZNF804A^+/+^ clones (**Fig. 2D-E**). We further assessed all clones for potential off-target changes following gene-editing using RT-qPCR, ICC and Sanger sequencing (**Fig. S6-S72**; Supplementary Text). Based on this analysis and due to the significant reduction in transcript and protein expression, we deemed ZNF804A mutant lines appropriate for further investigation into ZNF804A functionality in developing glutamatergic neurons.

### Transcriptomic analysis of mutant neurons reveals modest effects on gene expression

Previous functional studies have implicated ZNF804A with having a role in regulating gene transcription in NPCs and neuroblastoma cell models (Chapman et al., 2018; Chen et al., 2015; Hill et al., 2012). To test whether disruption of ZNF804A function results in significant transcriptome-wide gene expression differences in developing glutamatergic neurons, we conducted an RNAseq analysis of mutant neurons. Examination of aligned reads confirmed exon 3 excision in ZNF804A^+/−^ and ZNF804A^−/−^ mutant neurons. However surprisingly, exon 4 appeared to be transcribed in mutation clones, which contradicted the predicted outcomes from *in silico* modeling (**fig. S8**). Given the absence of any previous reports on ZNF804A isoform lacking exon 3, these data potentially suggests that ZNF804A mutant lines may give rise to the synthesis of a truncated protein. However, it is of note that no additional bands were observed in our western blot analysis using an antibody targeting the C-terminal of the protein (Fig 2D and FigSxx). Therefore, it is likely that mutant ZNF804A lines give rise to a dysfunctional protein or unstable mRNA.

Next, we assessed the impact of ZNF804A mutation on neuronal differentiation, through analysis of relevant gene expression patterns in mutant neurons. At D7, all clones had high expression of genes associated with a dorsal forebrain fate (*EMX1*, *DMRTA1* and *DMRT3*), while showing low expression of genes highly expressed in developing ventral forebrain (*NKX2*-1, *DLX2*), midbrian (*LMX1A*), and hindbrain (*HOXB1*) neurons (**Fig. 2F**). Consistent with the generation of developing upper layer cortical neurons, these cells also highly expressed *TLE1*, *CUX1* and *CUX2*, while only showing minimal expression of *FEFZ2* and *TBR1*. These results confirmed the generation of a population of developing upper layer glutamatergic cells (Nehme et al., 2018; Treutlein et al., 2016). Moreover, mutation of *ZNF804A* did not appear to disrupt cell fate acquisition as these expression patterns were consistent across all clones and genotypes (**Fig. 2F**).

We then examined transcriptomic variations caused by mutated ZNF804A by comparing ZNF804A^−/−^ and ZNF804A^+/−^ mutation genotypes with ZNF804A^+/+^ cells. Initial principal component analysis (PCA) of the gene expression data indicated some limited sample clustering (**fig. S9**), which may contradict its previously hypothesised role as a broad regulator of gene transcription (e.g., via its transcription factor-like properties of its C2H2 zinc-finger domain). Notably, samples with ZNF804A^−/−^ and ZNF804A^+/+^ genotypes clustered according to genotype, while no noticeable clustering was observed between clones within each genotype (**fig. S9**). To comprehensively evaluate the effect of the ZNF804A mutation on gene expression, we used three different gene expression comparison signatures, each analyzed using DESeq2. All differentially expressed genes (DEGs) of each signature are provided in **table S4**. Signature A (“Wildtype vs. Homozygous”) comparisons yielded a total of 45 DEGs with 23 genes down- and 22 genes up-regulated (**Fig. 3A**) and top 20 protein-coding DEGs (10 up- and 10 down-regulated) clustered according to genotype (**fig. S10A**). Signature B (“Wildtype vs. Heterozygous”) had fewer DEGs than Signature A, consistent with the lack of clustering of ZNF804A^+/−^ samples observed in the PCA analysis (**fig. S9A**), which is expected due to the milder, mono-allelic genomic effects of *ZNF804A* mutation. Nevertheless, 4 genes were significantly down-, and 6 genes up-regulated in heterozygous compared to wildtype conditions (FDR < 0.05) (**Fig. 3A**). The heatmap of all 10 DEGs generally showed clustering by genotype, except for replicate 2 of wildtype clone ZNF804A^+/+^_4, which appeared more similar to ZNF804A^+/−^ clones (**fig. S10B**). Finally, comparisons in Signature C (“Heterozygous vs. Homozygous”) yielded the highest number of DEGs (47 genes, FDR < 0.05), with only 7 genes down- and 35 genes up-regulated (**fig. S11A**). Heatmap clustering of the top 20 protein-coding DEGs (10 up-regulated and 10 down-regulated) confirmed the genotype-based impact on gene expression, as ZNF804A^−/−^ and ZNF804A^+/−^ clones clustered according to their genotype (**fig. S11B**).

**Fig. 3.**
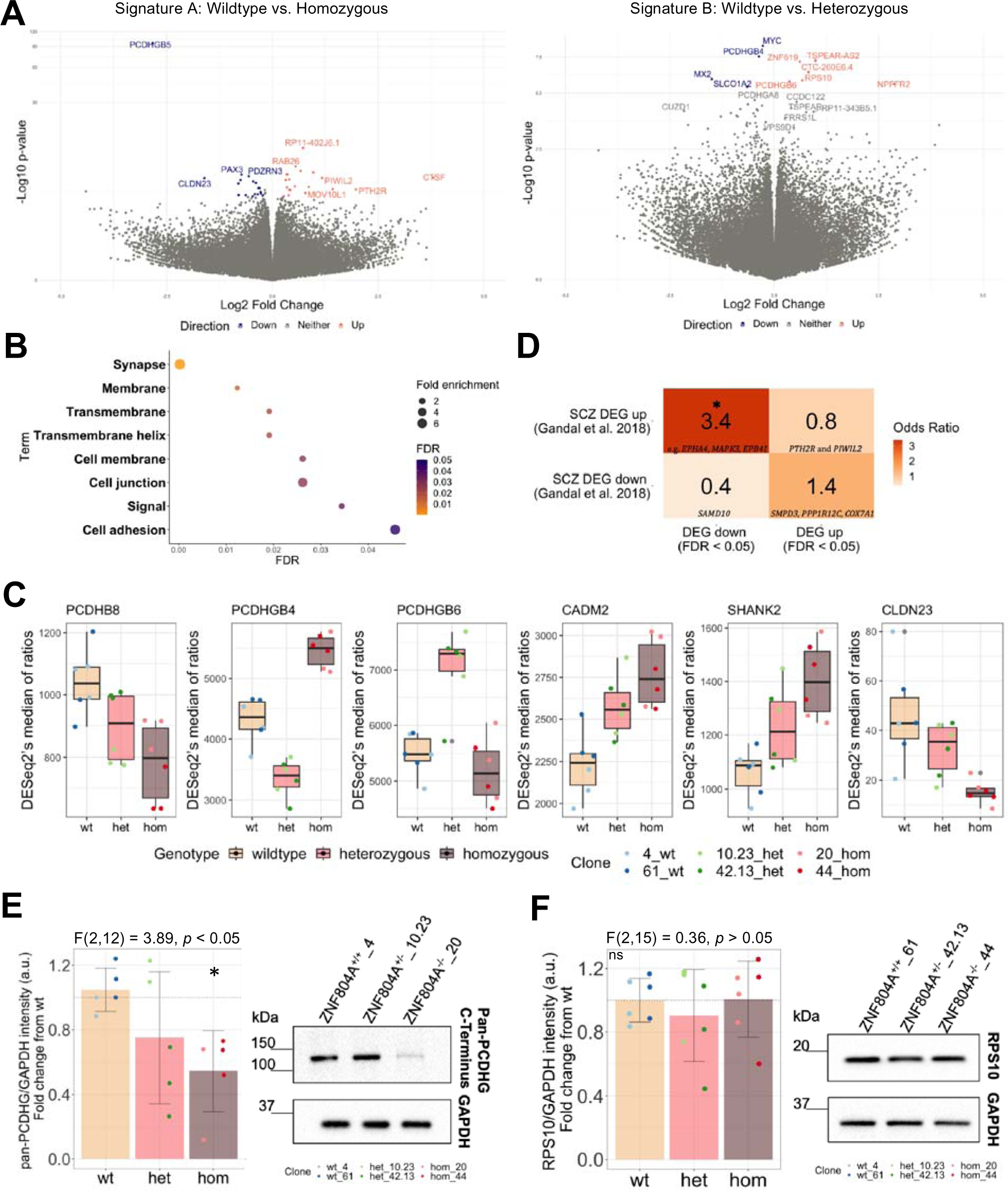
Transcriptomic profiling of mutant ZNF804A neurons show dysregulation of genes associated with cell adhesion and the glutamatergic synapse. (A) Volcano plots depicting differentially expressed genes (DEGs) in Signature A (left) and Signature B (right). The x-axis representing log2FC, y-axis -log10 p-value. Significantly up-regulated DEGs (FDR < 0.05) are indicated in red, significantly down-regulated DEGs in blue. (B) Gene ontology (GO) analysis of DEGs in Signature A (FDR < 0.07). X-axis displays FDR, y-axis lists terms with significant enrichment (FDR < 0.05). Icon size represents fold enrichment. (C) Boxplots depict normalized reads (DESeq2’s median of ratios) for counts of DEGs associated with cell adhesion and glutamatergic synapse. Individual clone counts are represented in overlapping dotplots, color-coded based on the clone. (D) Heatmap illustrating Fisher’s exact test results. The color key represents odds ratio results (dark orange = high odds ratio; light orange = low odds ratio). An asterisk denotes significance (FDR < 0.05) for the gene set overlap analysis between the significantly down-regulated gene set of Signature A and the significantly up-regulated gene set from post-mortem schizophrenia brain tissue (Gandal et al., 2018). Examples of overlapping genes are shown in the squares of their respective comparison sets. * FDR < 0.05. (E) Representative western blot and barchart showing reduced pan-gamma protocadherin (PCDHG) protein in ZNF804A^−/−^. The housekeeping protein glyceraldehyde-3-phosphate dehydrogenase (GAPDH) serves as loading control. Barchart depicting the quantification of integrated densities for pan-PCDHG band (∼120kDa). Overlapping dotplots display pan-PCDHG clone intensities. One-way ANOVA (F(2,12) = 3.89, p = 0.0498) with subsequent Tukey HSD post-hoc tests revealed significant reductions of pan-PCDHG in ZNF804A^−/−^. (F) Representative western blot and barchart showing no change in ribosomal protein S10 (RPS10; ∼19kDa) expression (F(2,15) = 0.36, p = 0.701). Error bars: mean ± standard deviation (SD), * p.adj. < 0.05.

To better understand how these DEGs may converge on shared biological or molecular processes, we investigated associations between gene products and gene ontology (GO) terms using the Database for Annotation, Visualization, and Integrated Discovery (DAVID) (Sherman et al., 2022). We subjected DEGs with corrected p-values < 0.07 to analysis against a predefined biologically relevant context, which included all genes expressed in the samples. Across functional annotations (including biological process, cellular component, and molecular function) and GO, terms including “Synapse” (Fold enrichment (FE) = 7.80, p < 0.001, False Discovery Rate (FDR) = 0.0002), “Cell Junction” (FE = 5.3, p < 0.01, FDR = 0.03), “Cell adhesion” (FE = 6.60, p < 0.01, FDR = 0.045) and “Signal” (FE = 1.8, p < 0.01, FDR = 0.03) were significantly associated with the DEG set (**Fig. 3B & table S5**). These data are consistent with those from previous transcriptomic analyses of ZNF804A function in human cellular models (Chapman et al., 2018; Hill et al., 2012). While each signature identified unique DEGs, there were substantial overlaps between the gene sets (**fig. S12**). Notably, genes related to cell adhesion proteins, such as protocadherins (e.g. *PCDHB8* (wt vs. hom: log2 fold change (log2FC) = −0.4, p < 0.001, FDR = 0.03; wt vs. het: log2FC = −0.2, p < 0.05, FDR > 0.05), *PCDHGB4* (wt vs. hom: log2FC = 0.3, p < 0.001, FDR < 0.001; wt vs. het: log2FC = −0.3, p < 0.001, FDR < 0.001), *PCDHGB6* (wt vs. hom: log2FC = −0.1, p > 0.05, FDR > 0.05; wt vs. het: log2FC = 0.4, p < 0.001, FDR = 0.005)), and glutamatergic synapse genes (e.g. *SHANK2* (wt vs. hom: log2FC = 0.4, p < 0.001, FDR = 0.002; wt vs. het: log2FC = 0.2, p < 0.01, FDR > 0.05)), exhibited dysregulation across all signatures, supporting a role for ZNF804A in synaptic regulation and formation (**Fig. 3C**).

Next, we sought to corroborate the association between ZNF804A and SZ through overlap analysis of DEGs from Signature A (“Wildtype vs. Homozygous”) with DEGs measured in *post-mortem* brain tissue of SZ patients (Gandal et al., 2018). Fisher’s exact test on two sets of DEGs (up and down-regulated) from Signature A vs. two sets of DEGs from *post-mortem* schizophrenia brain tissue samples against a physiologically relevant background (4 comparisons in total) revealed a significant overlap between down-regulated genes from Signature A and up-regulated genes found in schizophrenia *post-mortem* samples (n genes in Signature A down-regulated = 24, n genes in cases = 2274, overlap size = 7 genes, p = 0.011, OR = 3.4, FDR < 0.05; **Fig. 3D**). GO analysis of the overlapping genes did not yield significant terms at a 5% FDR threshold, but six of the seven DEGs matched a “Membrane” gene list that included *EPHA4, MAPK3,* and *EPB41.* Overall, both the data presented here and the existing literature (Chapman et al., 2018; Hill et al., 2012) support the hypothesis that ZNF804A mutation yields transcriptomic disruptions related to cell adhesion. However, the extent of this disruption at the protein level remains unclear. Thus, we assessed gamma protocadherin (PCDHG) protein expression in developing glutamatergic cortical neurons derived from wildtype and mutant cells. Pan-PCDHG protein levels were significantly reduced in ZNF804A^−/−^ cell lysates compared to ZNF804A^+/+^ controls, but ZNF804A^+/−^ mutant neurons did not have a statistically significant effect on protein abundance (one-way ANOVA: F(2,12) = 3.89, p = 0.0498; Tukey HSD: wt-hom: p.adj. < 0.05, all other comparisons p.adj. > 0.05; mean FC from wt: hom = 0.5, het = 0.3, **Fig. 3E**). Based on previous research linking disruptions in protein synthesis to ZNF804A function (Zhou et al., 2018) and results of the RNAseq analysis yielding differential expression for ribosomal protein S10 (RPS10) in ZNF804A^+/−^ conditions (log2FC = 0.7, p < 0.001, FDR = 0.005), we investigated the impact of ZNF804A mutation on RPS10 protein expression as above but found no significant effect (F(2,15) = 0.36, p = 0.701; **Fig. 3F**). Taken together, transcriptomic analysis of mutant ZNF804A neurons only revealed modest effects on gene expression contrary to previous reports suggesting a primary role for ZNF804A in regulating gene expression (Chapman et al., 2018; Chen et al., 2015). However, the limited number of genes showing differential expression includes those associated with synapses and cell adhesion, which are relevant for SZ neurobiology as previously reported in human cell models (Chapman et al., 2018; Chen et al., 2015; Hill et al., 2012).

### Altered expression of excitatory synaptic proteins along dendrites of mutant ZNF804A neurons

Results of transcriptomic analysis and previous literature (Deans et al., 2017; Hill et al., 2012) suggested disruptions of the glutamatergic synapse in ZNF804A mutation models. To investigate the potential impact of ZNF804A mutation on developing glutamatergic synapses, we examined the density of glutamatergic pre- and post-synaptic proteins and their co-localization across various cellular compartments. We conducted this analysis using high-throughput confocal imaging and automated image analysis, detecting puncta specifically in somata, neurites, and the entire cellular regions of MAP2^+^ cells. Protein puncta were analyzed relative to neurite length, somata count, and overall cell quantity and normalized to corresponding values obtained from wildtype clones within the same experimental setup. We collected data from 54,654 MAP2^+^ neurons (ZNF804A^+/+^_4: 4681, ZNF804A^+/+^_61: 11,445, ZNF804A^+/−^_10.23: 5,323, ZNF804A^+/−^_42.13: 7,323, ZNF804A^−/−^_20: 6,833, and ZNF804A^−/−^_44: 19,049) for post-synaptic GluN1 staining and 45,371 cells (ZNF804A^+/+^_4: 4,241, ZNF804A^+/+^_61: 10,302, ZNF804A^+/−^_10.23: 5,257, ZNF804A^+/−^_42.13: 5,580, ZNF804A^−/−^_20: 5,550, and ZNF804A^−/−^_44: 14,441) for pre-synaptic VGLUT1. While genotype had no significant effect on GluN1 puncta count in somata (Kruskal-Wallis test: *X*^2^(2) = 0.05, p = 0.9, n = 18) or entire cell regions (one-way ANOVA: F(2,15) = 0.3, p = 0.8), we observed a difference in puncta count when analyzing MAP2^+^ neurites (Kruskal-Wallis test: *X*^2^(2) = 8.49, p = 0.01, n=18). Specifically, Dunn’s test with holm corrections showed that the statistical significance was driven by *increased* puncta count in MAP2+ neurites of ZNF804A^+/−^ mutant neurons as compared to wildtype controls (p.adj. < 0.05, mean FC from wt = 4.7 (range: 1.1 – 20.7)). Although the *post-hoc* comparisons between ZNF804A^+/+^ and ZNF804A^−/−^ mutation neurons just failed to reach the threshold for statistical significance (p.adj = 0.055), it’s important to note that there was a clear numerical increase in GluN1 puncta count in MAP2+ neurites of neurons carrying ZNF804A^−/−^ mutations in all but one replicate, with a mean FC of 4.8 (range: 1.2 – 18.2; **Fig. 4A-C).** The same analysis for pre-synaptic VGLUT1 showed statistically significant effects for genotype on puncta counts within somata (one-way ANOVA: F(2,12) = 4.5, p = 0.03, n = 15), entire cell regions (Kruskal-Wallis test: *X*^2^(2) = 8.5, p = 0.007, n = 15), and puncta counts along MAP2^+^ neurites (Kruskal-Wallis test: *X*^2^(2) = 6.4, p = 0.04, n = 15). While *post-hoc* tests showed significant reductions of VGLUT1 puncta in somata (mean FC for hom from wt = 0.8 (range: 0.6 – 0.9), Tukey HSD: p.adj. < 0.05) and entire cell regions with ZNF804A^−/−^ and ZNF804A^+/−^ mutations (Dunn’s test with holm corrections: mean FC of hom from wt = 0.8 (range: 0.6 – 1), p.adj. < 0.05; mean FC of het from wt = 0.8 (range: 0.6 – 1), p.adj. < 0.05), neurites of ZNF804A^−/−^ (Dunn’s test with holm corrections: mean FC from wt = 3.3 (range: 1 – 8.3), p.adj. < 0.05) and ZNF804A^+/−^ mutation neurons (mean FC from wt = 3.1 (range: 1.1 – 6.9), p.adj. < 0.05) showed significant increase in VGLUT1 puncta counts (**Fig. 4A-B**). Analysis of further pre- and post-synaptic proteins (Bassoon, SV2A and PSD95) showed similar effects for both ZNF804A^−/−^ and ZNF804A^+/−^ mutations (**fig. S13A - F**). In fact, effect size analysis of main statistical tests revealed that ZNF804A genotype had the largest effects on VGLUT1 and PSD95 puncta in entire cellular regions (η^2^ = 0.65) and substantial effects on pre- and post-synaptic protein puncta in neurites (η^2^ = 0.37 – 0.42). Genotype had small effects on puncta counts in somatic regions except for a large effect on VGLUT1 puncta in somatic regions (η^2^ = 0.42; **Fig. 4D**). This indicates a pronounced impact of ZNF804A mutation on glutamatergic synaptic proteins in different cellular compartments of immature glutamatergic neurons.

**Fig. 4.**
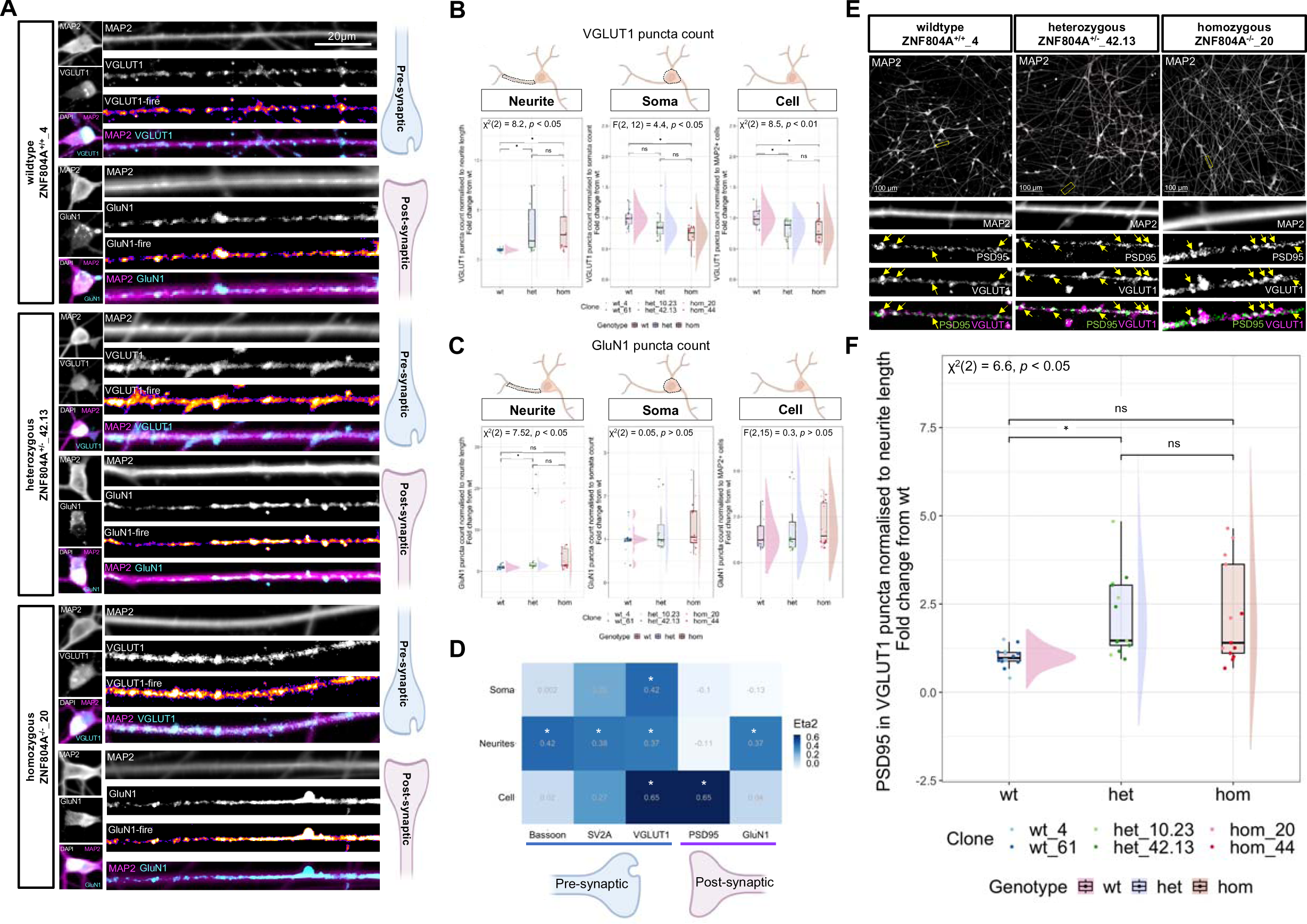
ZNF804A mutant neurites show increased localization of glutamatergic synaptic proteins. (A) Representative confocal images of D7 ZNF804A^+/+^, ZNF804A^+/−^, ZNF804A^−/−^neurons captured with high-content microscope. Cells were immunostained for pan-neuronal marker microtubule associated protein 2 (MAP2; magenta) to identify cellular regions. Puncta for NMDA Receptor Subunit 1 and vesicular glutamate transporter 1 (GluN1 and VGLUT1; fire LUT and cyan) are shown in neurites and somata. (B) Raincloud plot showing quantification of VGLUT1 puncta in different cellular compartments of MAP2^+^ neurons. Overlaying dotplots show puncta counts for each technical replicate. VGLUT1 puncta were significantly decreased in whole cellular regions (Kruskal-Wallis: X^2^(2) = 8.5, p = 0.007) of ZNF804A^+/−^ and ZNF804A^−/−^ neurons, and in somata (one-way ANOVA: F(2,12) = 4.5, p = 0.03) of ZNF804A^−/−^ neurons, whereas significant increases were observed along MAP2^+^ neurites of ZNF804A^+/−^ and ZNF804A^−/−^ neurons (Kruskal-Wallis: *X*^2^(2) = 6.4, p = 0.04, n = 15). (C) Raincloud plot showing quantification of GluN1 puncta as above. Puncta were significantly increased in MAP2^+^ neurites of ZNF804A^+/−^ neurons (Kruskal-Wallis: *X*^2^(2) = 7.52, p = 0.01, n = 18). (D) Heatmap illustrating effect sizes (Eta2/η2) based on ANOVA and Kruskal-Wallis tests of ZNF804A mutation on pre- and post-synaptic proteins in different cellular compartments. The impact of ZNF804A mutation on each synaptic protein (x-axis) was evaluated in each tested cellular compartment (y-axis). The magnitude of the effect size is represented using a blue color range. (E) Representative confocal image of high-content system of co-localised VGLUT1 and PSD95 puncta in D7 ZNF804A^+/+^, ZNF804A^+/−^, ZNF804A^−/−^ neurons. Cells were immunostained for MAP2 (greyscale), VGLUT1 (magenta), and PSD95 (green). Positive co-localisation is indicated with yellow arrowheads. (F) Raincloud plot showing quantification of co-localisation of VGLUT1 and PSD95 puncta as above. Co-localised puncta were significantly higher in ZNF804A+/− neurites (Kruskal-Wallis: *X*^2^(2) = 6.6, p = 0.04, n = 15) * p.adj. < 0.05

Next, we assessed co-localization of PSD95 and VGLUT1 puncta along neurites of 45,371 MAP2^+^ neurons (ZNF804A^+/+^_4: 4,241, ZNF804A^+/+^_61: 10,302, ZNF804A^+/−^_10.23: 5,257, ZNF804A^+/−^_42.13: 5,580, ZNF804A^−/−^_20: 5,550, and ZNF804A^−/−^_44: 14,441) to quantify pre- and postsynaptic marker intersection as a readout of putative immature synapses (**Fig. 4E**). Indeed, statistical analysis on averaged technical replicates showed significant group differences in co-localization quantities with genotype as an independent variable (Kruskal-Wallis test: *X*^2^(2) = 6.6, p = 0.04, n = 15). Dunn’s *post-hoc* analysis with holm corrections confirmed significant increases in co-localized VGLUT1 and PSD95 puncta along neurites of ZNF804A^+/−^ mutation neurons (mean FC from wt = 2 (range: 1.2 – 3.5), p.adj. < 0.05). Although we observed an increase in co-localization in all but one replicate, post-hoc comparisons of neurites carrying ZNF804A^−/−^ mutations and ZNF804A^+/+^ did not yield statistically significant results (mean FC from wt = 2 (range: 0.8 – 3.5), p.adj. > 0.05) (**Fig. 4F**). Overall, this data indicates that components of glutamatergic synapse are over-recruited to neurites in ZNF804A mutant neurons, potentially leading to excessive formation of developing synapses.

### Assessing neurite- and soma-specific proteomes in developing glutamatergic neurons

Our high-throughput confocal imaging analysis suggested an increased localization of synaptic proteins to MAP2^+^ neurites of ZNF804A mutant developing glutamatergic neurons, whereas the expression of synaptic proteins in the somata and entire cell region were either decreased or remained unchanged (**Fig. 4D**). Furthermore, bulk RNAseq did not reveal any genotype-related transcriptome-wide variations in these synaptic genes (**table S4**). This suggests that the increase in synaptic proteins within neurites likely occurs through a mechanism independent from changes to gene expression in a cell-compartment specific manner. Based on prior evidence that ZNF804A binds to ribosomal proteins and regulates protein translational efficiency (Zhou et al., 2018), we hypothesized that neurite-localized increases in synaptic proteins may be due to changes in local protein synthesis. Hence, we tested whether ZNF804A mutation leads to an alteration in the local proteome of distinct sub-cellular compartments. To this end, we separated the neurites and soma of developing glutamatergic neurons and used proteomic analyses to examine the local proteome of each sub-cellular compartment (Zappulo et al., 2017). Cell lines from all three genotypes were differentiated on a membrane with a laminin-coated underside allowing neurites to extend through pores, enabling the separation of cell bodies from neurite processes (**Fig. 5A**). We validated the effective separation of neurites from somata using immunostaining and western blotting (**Fig. 5B and fig. S14**). Analysis of Z-stacks imaged using a confocal microscope revealed that neurites mainly extended and grew on the bottom of the membrane, while 4′,6-diamidino-2-phenylindole (DAPI)-stained somata were only present on the top (**Fig. 5B**). We then proceeded to lyse neurites and somata grown on the same membrane separate from each other for proteomic analysis. We employed liquid chromatography-coupled tandem mass spectrometry (LC–MS/MS) in data-independent acquisition (DIA)-mode on a total of 35 samples (neurites: ZNF804A^+/+^ n = 6, ZNF804A^+/−^ n = 5, ZNF804A^−/−^ n = 6; somata: ZNF804A^+/+^ n = 6, ZNF804A^+/−^ n = 6, ZNF804A^−/−^ n = 6) to analyze the localized proteomes of both isolated neurites and somata. Using a label-free quantification (LFQ) approach, we successfully detected a total of 5,393 proteins (range per sample: 581 – 5,307, **Fig. 5C & table S6**). Initial unsupervised heatmap analysis showed hierarchical clustering of samples according to cellular compartment (**Fig. 5D**). To refine our analysis, we removed proteins expressed in fewer than 70% of the samples and three samples showing significantly lower identified proteins (ZNF804A^+/−^_10.23_RP3_soma, ZNF804A^+/−^_61_RP3_soma, ZNF804A^+/−^_42.23_RP3_soma; **Fig. 5D**).

**Fig. 5.**
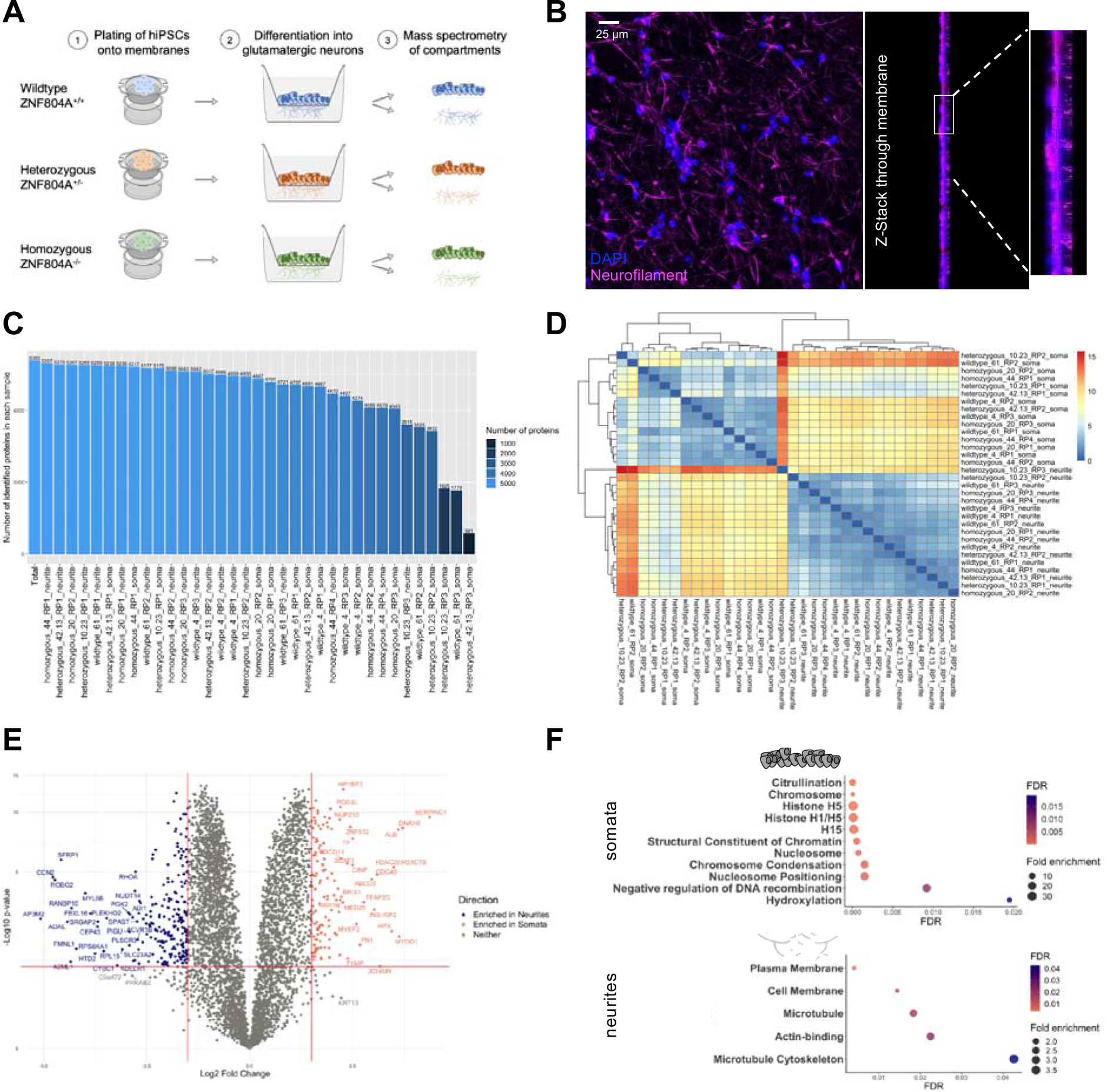
Successful separation of neurite and soma proteomes. (A) Schematic representation of cellular compartment separation assay. (B) Representative confocal image of and Z-stack through membrane confirming successful separation of neurites and somata. 4′,6-Diamidin-2-phenylindol (DAPI; blue)-stained somata remain on the surface of the membrane, whereas neurofilament (magenta)-rich neurites grow mainly on the bottom. (C) Barchart of number of identified proteins (y-axis) by mass spectrometry in each tested sample (x-axis). (D) Unsupervised clustering heatmap of all samples tested in proteomic approach. Dendrograms show clustering according to cellular compartment. (E) Volcano plot showing enrichment of proteins in neurites and somata. The x-axis represents log2FC, y-axis -log10 p-value. Neurite-enriched proteins (log2FC < −1.5; FDR < 0.05) are indicated in blue, soma-enriched proteins (log2FC > 1.5; FDR < 0.05) are indicated in red. (F) GO analysis of soma-enriched (top) and neurite-enriched (bottom) proteins. X-axis displays FDR, y-axis lists terms with significant enrichment (FDR < 0.05). Icon size represents fold enrichment.

To confirm the effectiveness of our approach to separate neurites from somata, we compared neurite- and soma-specific proteomes (regardless of genotype) by applying the *limma* package for differential protein expression analysis (Ritchie et al., 2015). This allowed us to assess proteome-wide differences between sub-cellular samples. In total, 351 proteins were differentially expressed (p.adj. < 0.05, log2FC > 1.5 or < −1.5), comprising 224 proteins showing greater enrichment in neurites and 127 proteins displaying a higher abundance in somata (**Fig. 5E & table S7**). As expected, GO analysis of differentially expressed proteins confirmed that neurite-enriched proteins are significantly enriched in gene lists termed “Cell Membrane” (FE = 1.5, p < 0.01, FDR = 0.01), “Actin-binding” (FE = 2.8, p < 0.001, FDR = 0.02) and “Microtubule Cytoskeleton” (FE = 3.5, p < 0.001, FDR = 0.04), whereas somata-enriched proteins overlapped with genes listed under “Nucleosome” (FE = 7.7, p < 0.001, FDR < 0.001), “Structural Constituent of Chromatin” (FE = 12.2, p < 0.001, FDR < 0.001) and “Chromosome Condensation” (FE = 21.1, p < 0.001, FDR = 0.001) (**Fig. 5F & table S8**). Thus, this data validates the successful separation of neurites and somata.

### Sub-cellular compartment-specific proteome analysis suggests over-recruitment of translational machinery to ZNF804A mutant neurites

Having validated the cellular compartment separation model, we next tested potential differences in the somatic and neurite proteomes of cells carrying ZNF804A mutations relative to isogenic control lines. Using the *limma* package, we carried out differential protein expression analyses employing the following comparisons:

(i) Neurite sections: wildtype vs. homozygous
(ii) Neurite sections: wildtype vs. heterozygous
(iii) Soma sections: wildtype vs. homozygous
(iv) Soma sections: wildtype vs. heterozygous

No proteins in any of these comparisons passed the FDR threshold of 5% suggesting no compartment specific proteome-wide differences attributable to ZNF804A mutation. To account for potentially more subtle or variable differences, we subjected all proteins that showed nominal differential expression (p < 0.05), up- or down-regulated, to exploratory GO analysis. Interestingly, up-regulated proteins in ZNF804A mutant neurites showed enrichment in terms associated with protein translation and homeostasis including “Ubiquitin-like protein transferase activity” (wt vs. hom: FE = 27.2, p < 0.001, FDR = 0.008), “Translation regulation” (wt vs. het: FE = 8.7, p < 0.001, FDR = 0.009) and “Ribosomal protein” (wt vs. hom: FE = 5.7, p < 0.01, FDR = 0.04) suggestive of excessive protein synthesis and homeostasis mechanisms in mutant neurites (**Fig. S15 & table S9**). Additionally, down-regulated proteins in ZNF804A^+/−^ mutant somata significantly clustered with gene lists termed “Ribonucleoprotein” (FE = 3.4, p < 0.001, FDR = 0.005) and “Ribosomal protein” (FE = 3.8, p < 0.001, FDR = 0.02; **table S9**), further suggesting an over-recruitment of translational machinery from somata to distal regions of mutated neurons. Next, we extracted log2-transformed LFQ values of up- and down-regulated proteins from “Ribosomal protein” and “Translation regulation” lists and tested potential genotype-dependent effects on abundance of these specific proteins within somata and neurites using one-way ANOVA. As expected, statistical analysis revealed cell compartment-specific differences in abundance of large ribosomal subunit 60S (neurite RPLP1: F(2,14) = 6.0, p = 0.013, η^2^ = 0.46), small ribosomal subunit 40S (soma RPS21: F(2,12) = 5.0, p < 0.001, η^2^ = 0.81), and RNA-binding proteins (neurite RBMXL1: F(2,14) = 5.2, p = 0.02, η^2^ = 0.43; **Fig. 6A & table S10**). Post-hoc tests confirmed significant differences in protein abundance within somata and neurites according to ZNF804A mutation genotype (**table S10**). Interestingly, proteins associated with translation regulation, including RACK1 (F(2,14) = 7.7, p = 0.006, η^2^ = 0.52), NCBP2 (F(2,14) = 5.4, p = 0.02, η^2^ = 0.44), and TARBP2 (F(2,14) = 4.5, p = 0.03, η^2^ = 0.39; **Fig. 6C & table S10**), only showed significant differences in abundance in ZNF804A mutant neurites, not in somata, confirmed by Tukey HSD post-hoc corrections (**table S10).**

**Fig. 6.**
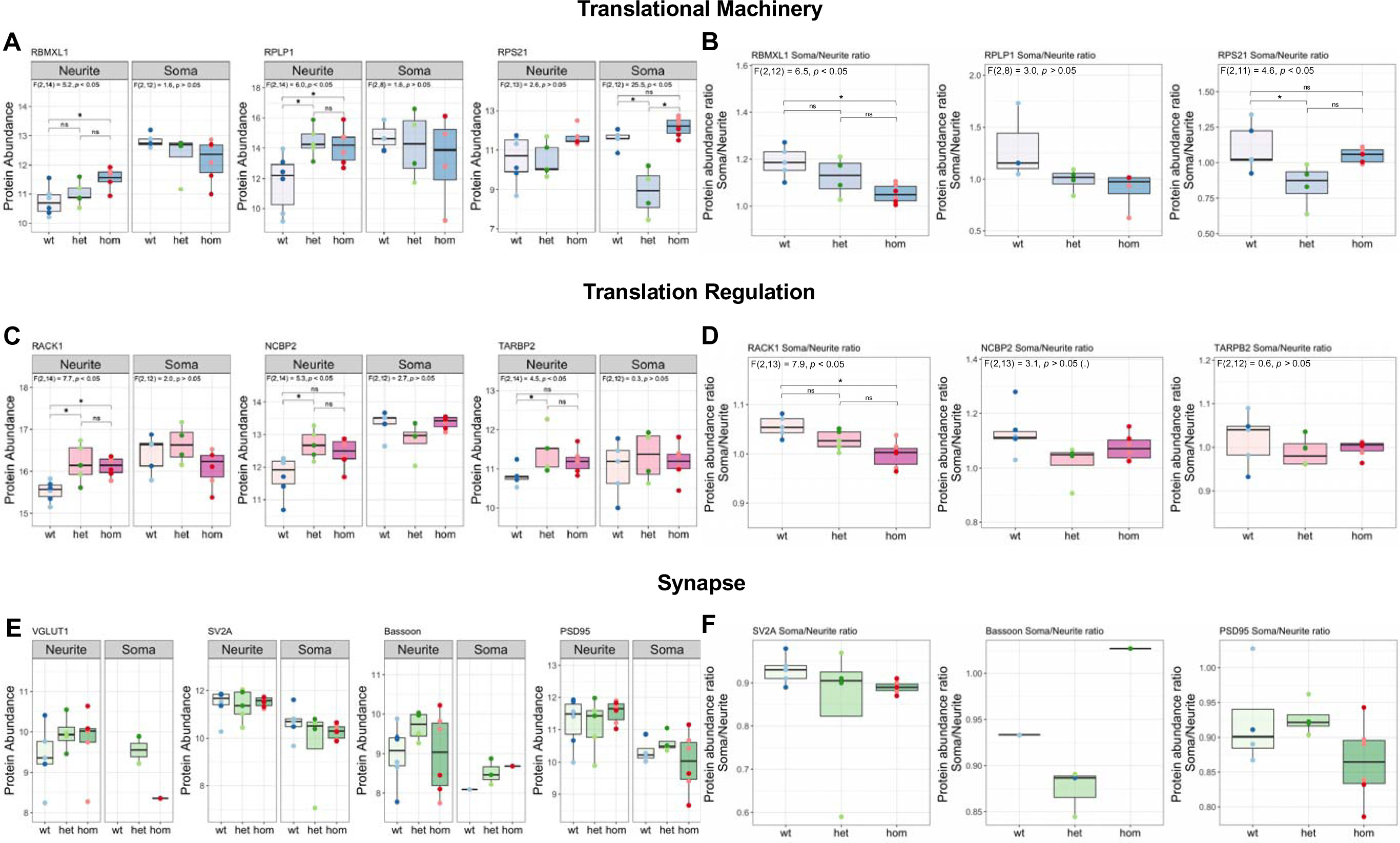
Over-recruitment of translational machinery to ZNF804A mutant neurites. (**A**) Boxplots showing LFQ protein abundances (y-axis) for proteins associated with translational machinery in somata and neurites of ZNF804A^+/+^, ZNF804A^+/−^, and ZNF804A^−/−^ neurons. RNA Binding Motif Protein, X-Linked Like 1 (RBMXL1), Ribosomal Protein Lateral Stalk Subunit P1 (RPLP1) and Ribosomal Protein S21 (RPS21). (**B**) Boxplots showing soma/neurite ratios of abundances of proteins above (y-axis). Overlaying dotplots show protein expression in each clone. One-way ANOVA results depicted in each graph. Boxplots of LFQ protein abundances (**C**) and soma/neurite ratios (**D**) of abundances of proteins associated with translational regulation. Receptor For Activated C Kinase 1 (RACK1), Nuclear Cap Binding Protein Subunit 2 (NCBP2), and TARBP2 Subunit of RISC Loading Complex (TARBP2). Plotted and analysed as above. Boxplots of LFQ protein abundances (**E**) and soma/neurite ratios (**F**) of abundances of proteins associated with the synapse. Vesicular glutamate transporter 1 (VGLUT1), Synaptic Vesicle Protein 2A (SV2A), presynaptic cytomatrix protein Bassoon, and post-synaptic density protein 95 (PSD95). Plotted and analysed as above. * p.adj. < 0.05; (.) p < 0.07

We then sought to evaluate whether synaptic proteins tested in the high-content imaging paradigm also showed significant differences in compartment-specific protein abundance. From those tested above, only VGLUT1, SV2A, Bassoon, and PSD95 were detected in our mass spectral analysis. Both VGLUT1 (F(2,11) = 0.7, p > 0.05, η^2^ = 0.12) and Bassoon (F(2,13) = 1.2, p > 0.05, η^2^ = 0.15) appeared to show increased abundance in neurites neurite compartments of ZNF804A^−/−^ lines, but one-way ANOVA statistical analysis did not reveal any significant differences between conditions or compartment (**table S10 & Fig. 6C**). It is of note, however, that our high-content imaging data specifically analyzed expression of synaptic proteins along/juxtaposed to MAP2^+^ dendrites, whereas the proteomic approach assesses protein expression in all dendrites and axons which may account for differences in sensitivity between these assays.

Lastly, we evaluated whether proteins may be over-recruited to distal cellular regions in ZNF804A^−/−^ and ^+/−^ mutant conditions by calculating an abundance ratio for each protein. Here we divided the log2-transformed LFQ values in the soma by those in the neurites within the same replicate. One-way ANOVA showed significant differences in ratios of proteins associated with the translational machinery (RBMXL1: F(2,12) = 6.5, p = 0.01, η^2^ = 0.52), RPS21: F(2,11) = 4.6, p = 0.03, η^2^ = 0.46) and translational regulation (RACK1: F(2,13) = 7.9, p = 0.006, η^2^ = 0.55). Intriguingly, Tukey HSD post-hoc analyses revealed effects in the same direction for all tested translational proteins (p.adj. < 0.05); soma/neurite ratios were significantly lower in ZNF804A^−/−^ and ^+/−^ mutant conditions compared to ZNF804A^+/+^ (**Fig. 6B & D**), suggesting an over-recruitment of these proteins to neurites from somatic regions. The same analysis was conducted for synaptic proteins sufficiently detected in both compartments (Bassoon and VGLUT1 protein was only identified in two and three soma samples, respectively). As above, one-way ANOVA testing yielded no significant differences in soma/neurite abundance ratios of SV2A (F(2, 12) = 1.1, p > 0.05, η^2^ = 0.15) or PSD-95 (F(2, 11) = 1.7, p > 0.05, η^2^ = 0.23). However, we observed trends towards lower ratios in ZNF804A^−/−^ mutant lines (**Fig. 6F**).

### ZNF804A mutant neurites show increased protein synthesis efficiency

As our proteomic data suggested an over-recruitment of translational machinery and regulatory effectors of protein synthesis to neurites in ZNF804A^+/−^ and ZNF804A^−/−^-mutant neurons, we next assessed whether this had a functional impact on protein translation. To this end, we employed the SUnSET assay in combination with ICC and high-throughput imaging to assess localized protein synthesis efficiency in MAP2^+^ neurites. Cell lines of each genotype underwent three distinct differentiations and prior to fixation, cells were treated with puromycin (10µg/ml) for 5 minutes. Puromycin-incorporation into the newly forming polypeptide chain was visualised using immunolabelling and high-content imaging confocal microscopy (**Fig. 7A**). An automated analysis script measured the mean intensity of puromycin staining in neurites of MAP2^+^ cells. A total of 35,461 MAP2^+^ cells (ZNF804A^+/+^_4: 8,539, ZNF804A^+/+^_61: 5,313, ZNF804A^+/−^_10.23: 7,268, ZNF804A^+/−^_42.13: 5,487, ZNF804A^−/−^_20: 4,970, ZNF804A^−/−^_44: 3,353) were identified in this experiment. As above, we performed statistical analysis on the averaged mean signal intensities of each clone per replicate to avoid pseudo-replication and we normalized puromycin signal intensities within neurites of each clone to the ZNF804A^+/+^ clone imaged in the same plate to calculate a FC difference. One-way ANOVA revealed significant differences in puromycin mean intensities within neurites due to genotype (F(2,15) = 4.8, p = 0.02). Interestingly, this statistically significant difference was attributed to an increase in puromycin staining intensities in neurites of ZNF804A^−/−^ mutation lines (Tukey HSD: mean FC from wt = 1.5 (range: 0.9 – 1.8), p.adj. < 0.05). Puromycin staining within neurites from ZNF804A^+/−^ mutation lines was not significantly different from ZNF804A^+/+^ counterparts (Tukey HSD: mean FC from wt = 1.1 (range: 0.6 – 1.8), p.adj. > 0.05, **Fig. 7B & C**). This implies that ZNF804A^−/−^ mutation results in a significant increase of newly synthesized proteins in neurites of developing human glutamatergic neurons, supporting the more exploratory proteomic findings. To determine if this increase is accompanied by elevated levels of key components of the translational machinery in distal regions beyond the soma, we visualized RPS6 staining in MAP2^+^ neurites using the high-content imaging approach described above. We imaged RPS6 puncta in three differentiations involving two clones per genotype (n=17, 6 wt, 5 het, and 6 hom – we excluded one replicate of ZNF804A^+/−^_42.13 due to staining failure). An automated analysis script identified RPS6 puncta in neurites of 55,150 MAP2^+^ cells (ZNF804A^+/+^_4: 6,712, ZNF804A^+/+^_61: 13,241, ZNF804A^+/−^_10.23: 8,984, ZNF804A^+/−^_42.13: 4,838, ZNF804A^−/−^_20: 7,772, ZNF804A^−/−^_44: 13,603). We normalized RPS6 puncta counts as above and observed a significant group difference when comparing counts across genotype (Kruskal-Wallis test: *X*^2^(2) = 7.7, p = 0.021; **Fig. 7D & E**). Post-hoc Dunn’s test with Holm adjustments revealed significant increases in RPS6 puncta in ZNF804A^−/−^ (mean FC from wt = 3.1 (range: 0.7 – 10.1), p.adj. < 0.05) and ZNF804A^+/−^ (mean FC from wt = 2.2 (range: 1.2 – 3.2), p.adj. < 0.05) conditions compared to ZNF804A^+/+^ (**Fig. 7E**), supporting proteomic findings indicating an over-recruitment of translational machinery to neurites. Lastly, to identify a potential mechanism driving these changes in local protein synthesis, we tested abundance of RPS6 phosphorylation at Ser235/236 using Western blotting. We adopted this approach because RPS6 phosphorylation is known to predominantly affect the translation of a specific subset of mitochondrial-related mRNAs, not having a global impact on translation (Puighermanal et al., 2017; Ruvinsky & Meyuhas, 2006), and it is phosphorylated at Ser235/236 near active synapses in response to synaptic activity (Pirbhoy et al., 2016), suggesting its potential significance in regulating the local proteome rather than the global one. While group comparisons of full protein abundances were not statistically significant (Kruskal-Wallis test: *X*^2^(2) = 5.6, p = 0.06, n=15), phosphorylated RPS6 exhibited a significant increase in both ZNF804A^−/−^ and ZNF804A^+/−^ mutation lines (one-way ANOVA: F(2,12) = 5.06, p = 0.02, n=15; Tukey HSD: wt-hom & wt-het: p.adj. < 0.05, het-hom p.adj. > 0.05; mean FC from wt: hom = 1.6 (range: 1.3 – 1.9), het = 1.62 (range: 0.9 – 2.2), **Fig. 7F & G**). Overall, these results are in support of the proteomic data suggesting ZNF804A mutation results in an over-recruitment of translational machinery to neurites and offer a novel perspective for the function of the schizophrenia susceptibility gene in local translational control.

**Fig. 7.**
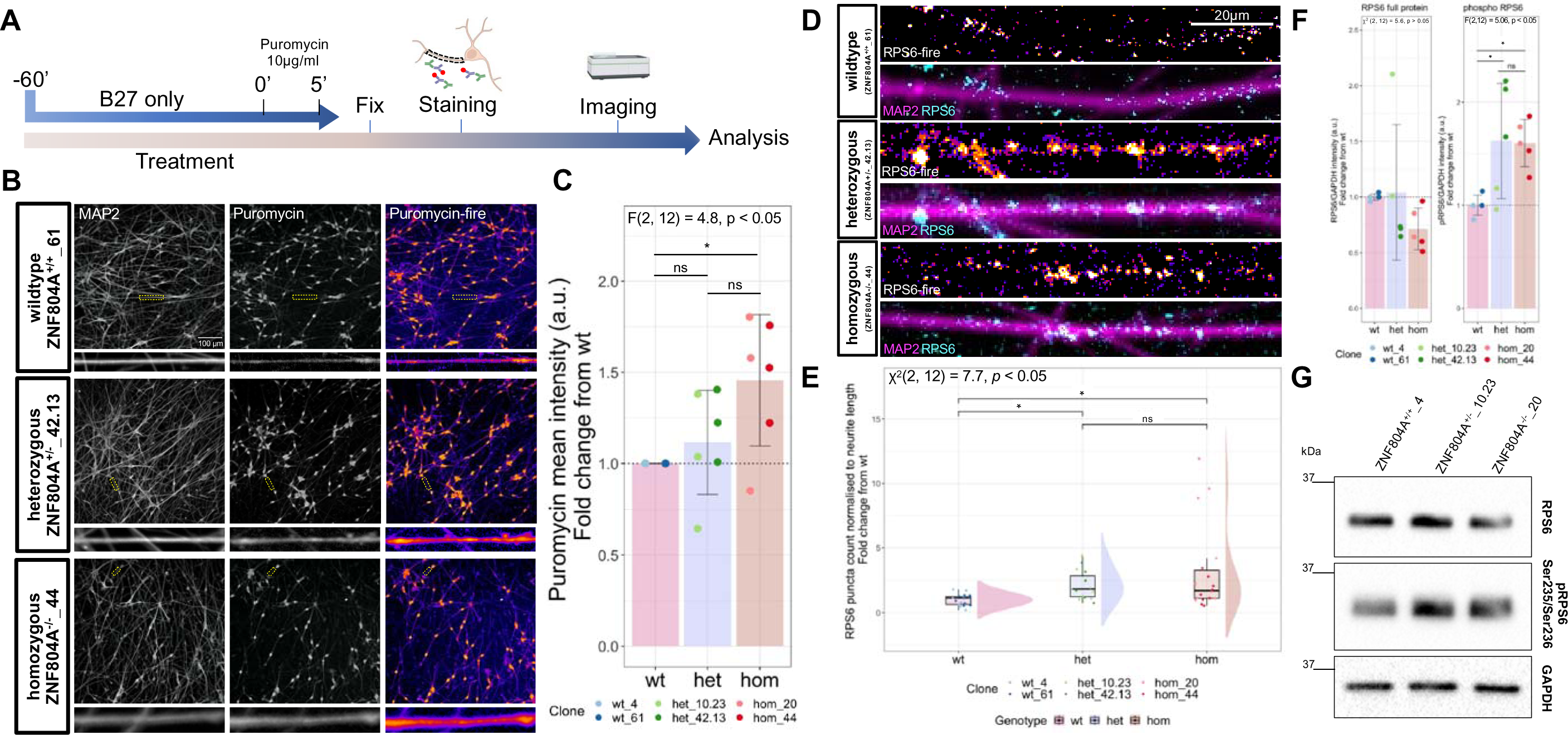
Local protein synthesis efficiency is increased in ZNF804A mutant neurites. (A) Experimental timeline of SUnSET approach in combination with high-content imaging. (B) Representative confocal image of puromycin incorporation (fire LUT) in MAP2^+^ neurites of ZNF804A^+/+^, ZNF804A^+/−^, ZNF804A^−/−^ D7 neurons captured with high-content microscope. (C) Barchart showing quantification of mean intensities of puromycin staining in MAP2^+^ neurites. Overlaying dotplots show staining intensities for each clone. ZNF804A^−/−^ show significantly increased puromycylated polypeptides in MAP2^+^ neurites (one-way ANOVA: F(2,15) = 4.8, p = 0.02, n =18). (D) Representative confocal image of ribosomal protein S6 (RPS6; cyan) puncta in MAP2^+^ neurites (magenta) of ZNF804A^+/+^, ZNF804A^+/−^, ZNF804A^−/−^ D7 neurons. (E) Raincloud plot showing quantification of RPS6 puncta in MAP2^+^ neurites. Overlaying dotplots show puncta counts for each technical replicate. RPS6 puncta were significantly increased in MAP2^+^ neurites of ZNF804A^+/−^ and ZNF804A^−/−^ neurons (Kruskal-Wallis test: *X*^2^(2) = 7.7, p = 0.021, n = 18). (F) Barcharts showing quantification of integrated densities (ID) of RPS6 total and phosphorylated (Ser235/Ser236) protein bands. Overlaying dotplots show IDs for each clone. Phosphorylated RPS6 was increased in ZNF804A^+/−^, ZNF804A^−/−^ D7 neurons (one-way ANOVA: F(2,12) = 5.06, p = 0.02, n=15) (G) Representative western blot of full and phosphorylated RPS6 (∼32 kDa) and housekeeper GAPDH. Error bars: mean ± standard deviation (SD), * p.adj. < 0.05

## Discussion

There remains a considerable gap in our understanding of observable neurobiological processes linked to the genetic susceptibility of complex psychiatric conditions such as schizophrenia (SZ). In this study we characterized the function of a gene robustly associated with SZ, *ZNF804A*, in a cell type and at a neurodevelopmental time point relevant for peak brain ZNF804A gene expression and SZ aetiology. We first established developing human glutamatergic neurons, starting to take on an upper layer cortical neuronal fate, as a suitable model to study ZNF804A gene function. We then applied a dual-sgRNA based CRISPR/Cas9 design to excise exon 3 of the ZNF804A locus. We successfully validated two clones per excision genotype with reduced ANF804A protein expression (ZNF804A^+/+^, ZNF804A^+/−^, ZNF804A^−/−^) and confirmed transcriptomic disruptions due to ZNF804A mutation in genes associated with cell adhesion and glutamatergic synapse using bulk RNAseq. Interestingly, dysregulated genes in ZNF804A^−/−^ were enriched for a gene set that has been found to be dysregulated in *post-mortem* brains of schizophrenia patients, which strengthens the role of ZNF804A in the disorder and supports the validity of our model. We then proceeded to test hypotheses generated by our RNAseq results and previous literature and assessed glutamatergic synaptic proteins and early putative synapses in mutation lines in a high-throughput confocal imaging experimental setup. Surprisingly, both pre- and post-synaptic proteins as well as putative synapses were increased in ZNF804A mutant MAP2^+^ neurites (**Fig. 5A – F**). However, mass spectrometry analysis of neurite and soma proteomes suggested no proteome-wide differences in protein expression according to genotype, but rather significant differences in neurite/somata protein abundance ratios of translational machinery and translation regulation proteins (**Fig. 6A – C**). This suggests an over-recruitment of translational machinery to ZNF804A mutant MAP2^+^ neurites, which was accompanied by increases in local protein synthesis efficiency (**Fig. 7B-C**). To our knowledge, the mechanisms by which ZNF804A functions at this stage of early neuronal development and in this cell type have not been analysed before and results of this study will aid in our understanding of how disruptions in this gene contribute to SZ aetiology during neurodevelopment.

A major consideration of our study design was to develop a cellular model that is relevant to both susceptibility gene function and the neurobiology of schizophrenia. As it has been proven difficult to translate the plethora of identified common disease-associated genetic variants into observable neurobiological processes (Trubetskoy et al., 2022), we aimed to take information about disorder-relevant cell types and developmental time-points into account, laying the foundation for a novel functional genetics approach. Previous research has identified the second trimester of foetal development as a developmental period when schizophrenia risk genes reach a peak in expression (Sey et al., 2020) and single-cell transcriptomics demonstrated that common risk genes were specifically expressed in immature glutamatergic neurons of the foetal frontal cortex and hippocampus (Cameron et al., 2023; Polioudakis et al., 2019). In line with this, existing literature (Hill & Bray, 2012; Tao et al., 2014) and our findings showed a peak of *ZNF804A* expression, both full-length and the SZ risk variant *ZNF804A^E3E4^,* in developing cell types, specifically in glutamatergic neurons with a forebrain, cortical upper layer identity. Notably, our statistical analysis yielded the largest effect size for developmental time-point on transcript expression, underlining the importance of taking temporal context specificity into account when characterizing gene function. It is also important to note that the SZ-associated SNP variant in ZNF804A is associated with a reduction in the expression of the gene (Gandal et al., 2018; Gusev et al., 2018; Hill & Bray, 2012). The CRISPR/Cas9-mediated excision of exon 3 allowed for the generation of isogenic cell lines with reduced expression of both full-length and SZ-associated *ZNF804A^E3E4^* isoforms (Tao et al., 2014), and thus in a similar manner to how the risk variant may impact the expression of both isoforms. Although holistic transcriptomic analysis revealed moderate effects, we confirmed disruptions in ZNF804A mutation lines linked to the glutamatergic synapse and cell adhesion, which is a replication of a robust finding in previous transcriptomic studies of ZNF804A function (Chapman et al., 2018; Hill et al., 2012). Furthermore, down-regulated DEGs from Signature A in our model were significantly enriched in an up-regulated gene set from *post-mortem* brain tissue from individuals with schizophrenia (FDR < 0.05) (Gandal et al., 2018). Intriguingly, the observed effect seemed to manifest in opposite directions, possibly due to development-related differences in gene expression (development vs. *post-mortem*) or a compensatory mechanism in adult brain. Despite these differences in *post-mortem* brain samples and our developmental model, a significant correlation between gene expression in idiopathic schizophrenia and ZNF804A mutation suggests shared molecular processes, reinforcing the role of ZNF804A in the disorder. Whilst it will be important to assess the functions of ZNF804A in cell lines with different gene risk backgrounds (Dobrindt et al., 2021) or when multiple risk genes are perturbed simultaneously (Deans et al., 2023; Schrode et al., 2019) these studies can be conducted with knowledge of the mechanism by which this risk gene impacts cellular and molecular function in developing neurons based on our work herein.

A key finding from our work is that ZNF804A potentially regulates early glutamatergic synaptogenesis in developing neurons. Interestingly, previous work finds associations between reduced ZNF804A expression and decreased dendritic spine density (Deans et al., 2017; Dong et al., 2020) and disruptions in activity-dependent spine plasticity (Deans et al., 2017). Consistent with this, our transcriptomic analysis of mutant ZNF804A neurons also showed significant disruptions in genes associated with the glutamatergic synapse. Surprisingly, however, high-content confocal imaging of synaptic proteins showed an increased redistribution of pre- and post-synaptic proteins to MAP2^+^ neurites with ZNF804A mutations and an increase of VGLUT1 and PSD95 overlap, suggesting a larger quantity of putative synapses. This, again, may be due to different developmental time-points analysed here and in previous studies; while decreases in dendritic spines were observed in more mature neurons with fully developed dendritic arbours, increases in early synapses in our cellular model may be an indication for the excessive formation of synapses due to ZNF804A mutation before neural circuitry is fully established. Indeed, a prominent theory is that excessive, or aberrant synaptic pruning contributes to synaptic dysfunction in SZ (Forrest et al., 2018; Forsyth & Lewis, 2017). Interestingly, studies using mutant mouse models with blocked neurotransmitter release have demonstrated that excitatory synapses can still form and function without glutamatergic neurotransmission (Sando et al., 2017; Sigler et al., 2017; Verhage et al., 2000), suggesting that a proportion of developmental synapse formation is activity-independent and potentially “hard wired” compared to postnatal and adult synaptogenesis. Thus, due to the difference in response to *ZNF804A* deficiency during the later stages of activity-dependent synaptogenesis compared to the initial phases of activity-independent synapse formation, ZNF804A might take on different roles at the synapse throughout neurodevelopment. Indeed, the increases in synapses observed here early in development could potentially have profound implications, affecting crucial processes in later life, such as synapse specification or synaptic pruning (Penzes et al., 2011; Südhof, 2018), ultimately contributing to disordered signal transmission as observed in schizophrenia.

One question that remains to be answered now is what caused this increase in putative early synapse formation in ZNF804A mutant neurons. Previously, ZNF804A has been shown to interact with ribosomal proteins (Zhou et al., 2018) and to play a role in structural remodelling of dendritic spines in response to activity-dependent stimulation (Deans et al., 2017). These data suggest a role for the susceptibility gene in local translational control. Dendritic translation is also robustly implicated in long-term synaptic plasticity in mature neurons (Huber et al., 2000; Kang & Schuman, 1996) and presynaptic axonal translation occurs during synapse formation (Wong et al., 2017). Additionally, while RNA granules have been shown to dynamically move along main axons and dendrites, there is some evidence that stationary RNA granules not only localize to pre-synaptic terminals (Wong et al., 2023), but also translocate towards synaptic spine heads upon neuronal stimulation (Mitsumori et al., 2017). This indicates facilitation of local translation-dependent synapse formation is based on the function of stationary ribosomes near synapses. An interesting observation in our data is that ribosomes as well as proteins associated with translation regulation seem to be excessively recruited to MAP2^+^ dendrites of ZNF804A mutation neurons, which was accompanied by increased dendritic translation efficiency and RPS6^Ser235/236^ phosphorylation, a phosphorylation site specifically associated with local translation initiation (Pirbhoy et al., 2016). ZNF804A dysfunction may therefore contribute to excessive redistribution of stationary ribosomes within immature MAP2^+^ dendrites, leading to increased synapse formation early on in development.

While ZNF804A expression is most pronounced in developing glutamatergic cells, transcripts of the susceptibility gene have also been found in other brain cell types including immature inhibitory neurons and radial glia (**Fig. 1A**). This suggests that its function is not exclusive to glutamatergic neurons. Therefore, the isogenic *ZNF804A* mutant model could be integrated with differentiation protocols aimed at generating cells of non-glutamatergic lineages, such as microglial (Couch et al., 2023) or GABAergic (Maroof et al., 2013) cells, enabling the characterization of gene function in mono- or co-culture paradigms. Lastly, translatability of findings is inevitably dependent on the characteristics of the study population. It becomes more and more evident that sex-differences are often overlooked but important in not only the genetics of psychiatric disorders but also their underlying neurobiology (Christiansen et al., 2022; Wingo et al., 2023). In the case of ZNF804A, there is evidence of a sex-specific SNP associated with schizophrenia (Zhang et al., 2011) as well as a mouse model indicating sex-dependent variations in schizophrenia-like phenotypes (Huang et al., 2020). Hence, future investigations should take biological sex into consideration when modelling gene function.

Overall, to our knowledge, this study is the first of its kind to characterize the function of schizophrenia susceptibility gene ZNF804A in a cellular model that takes developmental timepoint and cell type specificity into account. Furthermore, by demonstrating an excessive localisation of ribosomes to dendrites of developing glutamatergic neurons coupled with an increase in early putative synapses, we suggest a link of the previously identified processes of ZNF804A function, synapse maintenance and translational control, by its ability to mediate local protein synthesis efficiency. Ultimately, research building on our findings will contribute to unravelling cellular neurodevelopmental processes directly linked to the aetiology of schizophrenia, and potentially aid in the identification of targets for future therapeutic interventions.

## Materials and Methods

### Experimental Design

The overarching aim of this study was to characterize the function of a robust schizophrenia susceptibility gene, *ZNF804A*, in a cellular model system that is both (i) relevant to gene function and (ii) schizophrenia etiology. Therefore, to identify the most suited developmental timepoint and cell type we first conducted a comprehensive expression profile of ZNF804A throughout glutamatergic cell development. Using two distinct differentiation protocols to generate hiPSC-derived cortical NPCs and post-mitotic forebrain neurons in combination with RT-qPCR, ICC and confocal microscopy, we identified immature post-mitotic forebrain neurons taking on an upper layer identity as a suitable model to study ZNF804A gene function. Subsequently, we carried out CRISPR/Cas9 genome editing to excise exon 3 of ZNF804A, inducing mutations to the ZNF804A locus relevant to schizophrenia. To account for potential genetic, epigenetic and phenotypic heterogeneity within a cell culture prior to single-cell isolation, we validated two clones per ZNF804A mutation genotype (ZNF804A^+/+^, ZNF804A^+/−^, ZNF804A^−/−^). Thereafter, we conducted bulk RNAseq to determine transcriptional perturbations due to ZNF804A mutation. Based on these findings, we tested early immature synapses and pre- and post-synaptic protein expression in a compartment-specific automated analysis paradigm on confocal images of ZNF804A^+/+^, ZNF804A^+/−^, and ZNF804A^−/−^ glutamatergic neurons acquired with a high-content microscope. Further hypotheses were generated to test the involvement of ZNF804A in local translational control. These hypotheses were tested using mass spectrometry on neurite- and soma-specific proteomes as well as SUnSET to assess translational efficiency within dendrites of ZNF804A mutation cells. If not otherwise stated, no data was excluded. In **Fig. 1B**, NPCs derived from a given hiPSC line were considered to be biological replicates when they were generated from a different clone of a donor line. The rest of the study was carried out on CRISPR/Cas9-edited isogenic lines, with biological replicates considered to be independent when neurons were derived from genome edited hiPSC clones with a different passage number. Statistical analysis was carried out on averaged technical replicates to avoid pseudo-replication. All figure legends and/or figures themselves specify sample sizes and statistical tests used in the individual experiments. Detailed parameters of each experiment and analysis procedures can be found in the methodology sections below.

### Cell lines and hiPSC maintenance

HiPSC lines 014_CTM, M3_CTM and 127_CTM (**table S11**) were generated and characterized previously (Adhya et al., 2021; Couch et al., 2023) with participant recruitment and procedures adhering to the ‘Patient iPSCs for Neurodevelopmental Disorders (PiNDs) study’ with ethical approval from the NHS Research Ethics Committee at the South London and Maudsley (SLaM) NHS R&D Office. Genetic editing was performed in a male neurotypical hiPSC line containing NGN2 under the control of an inducible, Tet-ON-controlled transgene overexpression system (NGN2-OPTi-OX). This cell lines was a kind gift from Dr. Mark Kotter (University of Cambridge) and has previously been described (Pawlowski et al., 2017). HiPSCs were maintained in Geltrex™-coated (Gibco; A1413302) plates in StemFlex media (Gibco; A3349401) in hypoxic conditions, media was changed every 48h, cells were passaged at 70-90% confluency using Versene (Gibco; 15040066), and neuralized between passages 10-24.

### Cortical neuroprogenitor differentiation

To characterize ZNF8004A expression throughout NPC development, cortical NPCs were differentiated following a previously validated modified dual SMAD inhibition (SMADi) protocol (Shum et al., 2020). In brief, neuralization was induced in hiPSC cultures when they reached at least 90% confluency by adding neuralization medium containing small molecule inhibitors LDN193189 (1µM; Sigma; SML0559) and SB431542 (10µM; Cambridge Bioscience; ZRD-SB-50). Neuralization medium was changed daily until D7, when cells were passaged (1:1) using StemPro® Accutase® (Gibco; A1110501) and 10µm Rho-associated, coiled-coil containing protein kinase (ROCK) inhibitor (ROCKi; Enzo Life Sciences, LKT-Y1000-M005) to a new plate, which contained NPC maintenance medium without SMADi (**table S12**). Further neuropassages were conducted on D12, D16, and D18.

### Combining OPTi-OX system with inhibitors to generate patterned forebrain neurons

The NGN2-OPTi-OX hiPSC line utilizes inducible *NGN2* transgene overexpression for rapid neuronal differentiation; however, there is a concern regarding the potential heterogeneity of these neurons (Treutlein et al., 2016). To address this, a differentiation protocol was tailored for differentiation of more patterned forebrain neurons by combining inducible *NGN2* forward-programming with Wnt inhibition and SMADi (Nehme et al., 2018) (**figure S16**).

Plates were coated with Poly-D-Lysine (PDL; Millipore; A-003-E) in 1x borate buffer (Thermo Scientific; 28341) for 4h at 37°C, followed by laminin (Sigma; L2020) overnight at 37°C. See **table S23** for plating densities and PDL/laminin concentrations according to experimental design. The terminal plating process included rinsing hiPSCs with HBSS, incubating them with Accutase (4min), counting cells with an automated cell counter (TC10 Bio-rad), and plating them at appropriate densities (**table S13**) using D1 neuralization medium supplemented with 10μM ROCKi, 2µM doxycycline hyclate (dox; Sigma, D9891), SMADi (as above), and Wnt (2µm; XAV939; Sigma-Aldrich, X3004) inhibitors to facilitate patterned differentiation into glutamatergic forebrain neurons. SMAD and Wnt inhibition were discontinued after D3 and the neurons were matured with daily medium changes involving 10µM N-[2S-(3,5-difluorophenyl)acetyl]-L-alanyl-2-phenyl-1,1-dimethylethyl ester-glycine (DAPT; Santa Cruz Biotechnology, SC-201315A), neurotrophic factors (brain-derived neurotrophic factor (BDNF; 10ng/ml; PeproTech, 450-02) and Glial Cell Derived Neurotrophic Factor (GDNF; 10ng/ml; PeproTech, 450-10), and B27-based maturation medium until D7. Daily media changes were conducted. Developing glutamatergic neurons were transfected with peGFP-N1 (BD Biosciences Clontech) using a calcium phosphate transfection approach. At D4, 1 µg of plasmid DNA was mixed with 1M CaCl2 and HBS (50mM HEPES buffer, 280mM NaCl, and 1.5mM Na2HPO4; pH 7.0); solution was allowed to precipitate for 30 minutes, before being added dropwise to D3 glutamatergic neurons. Cells were incubated with calcium phosphate/DNA mixture for 3 hours, before being replaced with B27-media with Ri; cells were fixed at D7. Detailed timeline of differentiation protocol and media compositions are listed in **table S14 and figure S16**.

### Genome engineering

CRISPR, CRISPR-associated protein 9 (Cas9) and a dual sgRNA approach was chosen to excise exon 3 of *ZNF804A*. CRISPR RNA sequences (crRNA) were designed for sgRNAs to target either intron 2 or intron 3 of ZNF804A using the Benchling web tool (https://www.benchling.com/). Before their *in vitro* application, the specificity and efficiency of sgRNAs were evaluated using online resources CRISPOR (http://crispor.tefor.net) and IDT (https://eu.idtdna.com/pages) (**table S15**).

CRISPR/Cas9 complexes were delivered into the nuclei of OPTi-OX hiPSCs via nucleofection. Prior to this, cells were cultured on a Geltrex-coated 10cm petri dish and allowed to proliferate in StemFlex medium until reaching 70% confluency. On the day of nucleofection, the CRISPR/Cas9 complex was formed following manufacturer’s instructions. Briefly, crRNA:tracrRNA was annealed, added to Cas9 protein (2µl), and incubated for 20min to form the ribonucleoprotein (RNP) complex. 30min before nucleofection, StemFlex was supplemented with RevitaCell™ (Gibco; A2644501; StemFlex:RC hereafter) to promote cell survival. For nucleofection, hiPSCs were incubated in Accutase (4min). After trituration and cell straining (40µm cell strainer, Falcon; 08-771-1), 1×10^6^ cells were added to 10ml StemFlex:RC, pelleted by 2min centrifugation at 1250 RPM, and the nucleofection reaction was prepared using the Cell Line Nucleofector™ Kit V (Lonza; VCA-1003) as per manufacturer’s instructions. The cell pellet was resuspended in nucleofection buffer, combined with the CRISPR/Cas9 construct and electroporation enhancer, and nucleofected using the Amaxa® Nucleofector® II and Nucleofector^®^ programme B-016. Following nucleofection, cells were transferred to Synthemax (Corning; 734-2634)-coated 6-well plates containing StemFlex:RC and incubated for 24h. A full-media change with StemFlex supplemented with CloneR^TM^ (Stemcell Technologies; 05888; 1:10 dilution) was performed once, and 24h later, CloneR was omitted from the medium to allow for hiPSC recovery. Media changes were carried out every 48h until cells reached 70% confluency.

To obtain a homogenous cell culture with ZNF804A mutation genotypes, single-cell cloning was performed. Cells were transferred to two Synthemax-coated 10cm dishes at low densities (500 and 1000 cells) using Accutase and 40µm cell strainers. After single-cell passaging, hiPSCs were allowed to recover and form colonies in StemFlex supplemented with CloneR^TM^ (1:10 dilution) as per manufacturer’s instructions, to ensure robust cloning efficiency and single-cell survival. Subsequently, single cell-derived colonies were allowed to proliferate for a week with full media changes every 48h.

For clone picking, individual colonies were selected and transferred into a Geltrex-coated Nunc™ MicroWell™ 96-Well plate (Thermo Fisher; 243656). Individual colonies, each originating from a single cell, were scraped using a P20 micropipette and transferred to separate wells containing 100µl of Stemflex:RC in preparation for genotyping. After 24h, medium was fully replaced, excluding RC, and after a two-day recovery period, the colonies were ready for genotyping.

### Cellular compartment separation assay

Separation of neurites and somata was achieved by differentiating and maintaining neurons on Nunc™ Polycarbonate Cell Culture Inserts for 6-well plates with a 3µm pore size (Thermo Fisher Scientific; 140642). Prior to cell plating, the inserts were subjected to a bottom coating process involving the application of PDL in borate buffer (25µg/ml), followed by laminin (20 µg/ml) only to the bottom of the membrane. This served as a guidance cue for the cells, facilitating the extension of their processes through the membrane towards the lower side. HiPSCs were terminally plated at a density of 500,000 cells/insert and differentiated into developing glutamatergic forebrain neurons using the protocol above. Media changes were performed daily by applying 2ml to the bottom and 2ml to the top of the insert until D7, when protein extraction was performed separately on neurites and somata.

Freshly prepared DIGE buffer (7M urea, 2M thiourea, 30mM TrisBase, pH 8.0) was used for protein extraction, with the procedure taking place on ice. Somata and neurites were extracted separately from the same membrane. First, membranes were transferred into clean 6-well plates with ice cold phosphate-buffered saline with magnesium and calcium (PBS-MC Gibco; 14040141). Somata were then dislodged from the upper surface of the membrane through gentle trituration with ice-cold PBS, transferred to a pre-cooled microcentrifuge tube and pelleted at 2000g for 2min at 4°C. Meanwhile, the remaining somata fractions were removed from the top of the membrane with a cotton-tip applicator, any excess PBS aspirated from the top and bottom sides, and the insert was then placed upside down on the bench. The membrane was excised from the insert using a sharp scalpel. It was then held bottom-up with forceps, and a cut was made from the circumference to the center using scissors. The resulting membrane piece was placed on top of a microcentrifuge tube. By applying gentle pressure with forceps, the center of the membrane was pushed down into the tube, creating a funnel shape until it was fully immersed in 150µl DIGE buffer. The tube was briefly vortexed, and after a 15-second spin, the membrane was discarded. Meanwhile, the centrifugation of the somata had completed, and the pelleted somata fractions were resuspended in 150µl DIGE buffer. Both fractions were then prepared for mass spectrometric analysis.

### RNA sequencing

Bulk RNAseq was conducted on 18 samples, which included ZNF804A^+/+^, ZNF804A^+/−^, ZNF804A^−/−^ lines (n = 2 clones per genotype and 3 differentiations per clone) at Genewiz Inc in Azenta Life Sciences, South Plainfield, NJ. The libraries were prepared using the NEBNext Ultra II RNA Library Prep Kit for Illumina (NEB, Ipswich, MA, USA), and the preparation involved a polyA selection method. The libraries were quantified with a Qubit 4.0 Fluorometer (Life Technologies, Carlsbad, CA, USA) and assessed for RNA integrity using an RNA Kit on an Agilent 5300 Fragment Analyzer (Agilent Technologies, Palo Alto, CA, USA). The sequencing libraries were then multiplexed and loaded onto the Illumina NovaSeq 6000 instrument’s flowcell, and sequencing was performed with a 2×150 Pair-End (PE) configuration v1.5, as per the manufacturer’s instructions. Lastly, image analysis and base calling were carried out using the NovaSeq Control Software v1.7 on the NovaSeq instrument. Information about the number of reads and quality scores per sample can be found in **table S16**. The FastQ screen tool (Wingett & Andrews, 2018) was employed to ensure quality control of the FASTQ files, which were then aligned to the human reference genome (GRCh38) using STAR (Dobin et al., 2013). Subsequently, the aligned files were sorted based on chromosomal coordinates using the samtools sort function (Danecek et al., 2021), and duplicate reads were identified using the Picard command MarkDuplicates (http://broadinstitute.github.io/picard/). A counts table, providing information about the number of aligned reads overlapping exons, was generated using the HTseq tool htseq-count (Anders et al., 2015). This table was used for downstream DGEA.

### Differential gene expression and gene set enrichment analyses

DGEA was performed on the transcriptomes from ZNF804A^+/+^, ZNF804A^+/−^, ZNF804A^−/−^ lines using the default Wald test in the DEseq2 package with the following comparisons:

**Signature A**: “Wildtype vs. Homozygous”, ZNF804A^+/+^_4, ZNF804A^+/+^_61, n = 6 and ZNF804A^−/−^_20, ZNF804A^−/−^_44, n = 6. We used ZNF804A^+/+^ as baseline for differential gene expression analysis (DGEA).

**Signature B**: “Wildtype vs. Heterozygous”, ZNF804A^+/+^_4, ZNF804A^+/+^_61, n = 6 and ZNF804A^+/−^_10.23, ZNF804A^+/−^_42.13, n = 6. We used ZNF804A^+/+^ as baseline for DGEA.

**Signature C**: “Heterozygous vs. Homozygous”, ZNF804A^+/−^_10.23, ZNF804A^+/−^_42.13, n = 6 and ZNF804A^−/−^_20, ZNF804A^−/−^_44, n = 6. We used ZNF804A^+/−^ as baseline for DGEA.

DEGs were identified based on an adjusted p-value threshold of < 0.05 using the Benjamini-Hochberg method. Enrichment analyses were performed on DEGs using the DAVID Gene Functional Classification Tool (Sherman et al., 2022). The DEG sets were tested against a physiologically relevant background, which included all genes meeting DESeq2’s internal filtering criteria for expression, i.e. those with adjusted p-values not equal to NA. Significant terms were determined with an adjusted p-value of < 0.05.

To assess the overlap of genes affected by ZNF804A mutation with those in *post-mortem* brain samples from schizophrenia cases (Gandal et al., 2018), Fisher’s exact tests were applied using GeneOverlap (https://bioconductor.org/packages/release/bioc/html/GeneOverlap.html), considering FDR correction with significance at FDR < 5%.

### Western blotting

D7 neurons were collected in ice-cold RIPA buffer, including protease and phosphatase inhibitors, followed by sonication (10 pulses at 40%). Protein concentrations were measured at 562nm using the Pierce Bicinchoninic Acid (BCA) kit (Thermo Scientific; 10678484). Proteins were prepared for SDS-PAGE separation by denaturation with 2x Laemmli sample buffer (Bio-Rad Laboratories; 161-0737) with 355mM β-mercaptoethanol for 5min at 95°C.

Proteins (5-10µg) were loaded onto self-made acrylamide gels (10-15%), separated at 100V for approximately 90min, and transferred onto PVDF membranes (BioRad; 1620177) at 78mA overnight at 4°C. The membranes were blocked in 5% BSA (Sigma; A7906) diluted in TBS-T solution, followed by primary antibody incubation overnight, washes in TBS-T, and secondary antibody incubation for 1h at room temperature. After a final wash, membranes were incubated in ECL Western Blotting Substrate (GE Healthcare; RPN2106) before imaging. See **table S17** for antibody details.

Membrane images were analyzed using Image Studio™ Lite software version 5.2 (LICOR). Integrated density (ID) of each protein signal was measured and background corrected using the software’s automatic detection. Signal quantifications were processed in Microsoft Excel. The ID of each protein was normalized to the housekeeping gene (GAPDH) probed on the same blot. To calculate FC compared to wildtypes, these ratios were adjusted for potential batch effects by dividing them by the wildtype ratio from the same membranes. The normalization process followed the provided formulas, with ΔΔID representing the normalized ID value used for statistical analysis and data visualization:

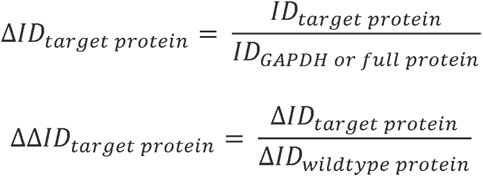

### In-gel digestion

For proteomic analysis, samples were prepared trough a tryptic in-gel digestion as described before (Barkovits et al., 2020) with slight modifications. Briefly, 20µg of protein were loaded on self-made acrylamide gels (12%) for neurites and somata fractions. After proteins were separated for 15 minutes at 50V and 2-3 minutes at 180V, protein bands were excised after staining with Coomassie Blue and destaining with 10% acetic acid. Afterwards, remaining dye was removed using repetitive washing steps with 50mM ammonium bicarbonate and 25mM ammonium bicarbonate/50% acetonitrile. Before tryptic digestion, destained gel pieces were dried in a Concentrator plus (Eppendorf), and afterwards rehydrated with 50mM ammonium bicarbonate. Afterwards, 0.5µg of trypsin (Serva) was added and samples were incubated overnight at 37°C and 300rpm. Digestion was stopped through the addition of trifluoric acid and peptides were eluted through consecutive incubation steps with 50% acetonitrile/0.05% trifluoric acid. Resulting peptide solution was concentrated in a fresh Eppendorf tube and reconstituted in 20µl of 0.1% TFA. 0.5µl of this solution was afterwards used for mass spectrometric measurements.

### Mass spectrometric analysis

The LC–MS/MS measurements were performed on an Ultimate 3000 RSLC nano LC system (Dionex) coupled to an Orbitrap Fusion Lumos Tribrid mass spectrometer (Thermo Fisher Scientific). Peptides were concentrated on a pre-column (Acclaim PepMap nanoViper, Thermo Fisher Scientific; 100μm. 2cm, 5μm particle size) and washed for 7 min with 0.1% TFA at 30μl/min. Subsequently, the pre-column was connected to an analytical C18 column (Acclaim PepMap nanoViper, Thermo Fisher Scientific; 75μm. 50cm, 2μm particle size). Separation of peptides was performed at a flow rate of 400nL/min with a gradient starting with 95% solution A (0.1% formic acid) and 5% solution B (84% acetonitrile, 0.1% formic acid). The concentration of solution B was increased up to 30% after 105min, then within 2 min to 95% and maintained at that level for 3 further minutes. Subsequently, the column was adjusted back to 5% solution B.

For DIA MS runs, the MS1 full scans were performed at a mass range of 350–1400*m*/*z* (mass/charge) with a resolution of 120,000 at 200*m*/*z* (AGC 4 × 10^5^, 20ms maximum injection time). Fragment analysis (MS2) was subdivided into 45 DIA isolation windows of different widths (10.5 – 268.4*m*/*z* wide) using a resolution of 30,000 at 200*m*/*z* (AGC 5 × 10^4^, 54ms maximum injection time).

### Mass spectrometric data analysis

DIA data were analyzed with Spectronaut (Version 16.5, Biognosys) in a direct-DIA approach using the human reference proteome obtained from Uniprot (UniProt, 2023) using the following settings. Calibration was set to non-linear iRT calibration with precision iRT enabled. Identification was performed using 1% q-value cutoff on precursor and protein level whereas the maximum number of decoys was set to a fraction of 0.1 of library size. The mass tolerance for matching precursor and fragment ions was set to dynamic (default), which lets SN determine the optimal value. For quantification interference correction was enabled with at least three fragment ions used per peptide, major and minor group quantities were set to mean peptide and mean precursor quantity, respectively with Top3 group selection each. Quantity was determined on MS2 level using area of XIC peaks with enabled cross run normalization. Proteomic data has been deposited to the ProteomeXchange Consortium via the PRIDE partner repository (Perez-Riverol et al., 2022) with the dataset identifier PXD047788.

Prior to statistical analysis, the list of identified proteins was filtered based on *NA* values; only proteins that were identified in 70% of the samples were subjected to further analysis. Neurite- and soma-specific proteomes were assessed using the limma package’s integrated moderated t-statistic and makeContrasts function with res = “detected proteins in neurite samples” – “detected proteins in soma samples”. Neurite-enriched proteins were identified based on an adjusted p-value threshold of < 0.05 using the Benjamini-Hochberg method (?) and a log2 fold change of < −1.5. Soma-enriched proteins were identified based on the same adjusted p-value threshold and a log2 fold change of > 1.5. Neurite- and soma-enriched list of proteins were then subjected to enrichment analyses using the DAVID Gene Functional Classification Tool as described above.

The same statistical analyses were performed on proteomic comparisons based on genotype with the following designs:

(i)Neurite sections: wildtype vs. homozygous (n = 6 wt & 6 hom clones)
(ii) Neurite sections: wildtype vs. heterozygous (n = 6 wt & 5 het clones)
(iii) Soma sections: wildtype vs. homozygous (n = 5 wt & 6 hom clones)
(iv) Soma sections: wildtype vs. heterozygous (n = 5 wt & 5 het clones)

### Surface Sensing of Translation (SUnSET)

Protein synthesis rates were assessed in MAP2^+^ neurites of developing glutamatergic neurons using SUnSET assay (Schmidt et al., 2009). Puromycin (Sigma; P8833) incorporation was validated at a 10µg/ml concentration and puromycin was allowed to integrate into the newly forming polypeptide chain of proteins for 5min before fixation, and an anti-puromycin antibody (Millipore; MABE343) was used to visualize the rate of protein synthesis through ICC.

### Immunocytochemistry

Cells were washed in PBS-MC and fixed using either double-fixation with 4% paraformaldehyde (PFA) in 4% sucrose in PBS-MC (10min) and 100% methanol on ice (10min) or single fixation with 4% PFA in 4% sucrose only (SUnSET experiment). After fixation, cells were washed with PBS-MC, followed by incubation in a permeabilization and blocking solution containing Triton-x100 and 2% Normal Goat Serum (NGS; Cell signalling: 5425S) for 1 hour at room temperature. Primary antibodies were applied overnight at 4°C, followed by washing and incubation with Alexa Fluor® secondary antibodies for 1 hour at room temperature. Cells were then stained with DAPI and cells/membranes on coverslips were mounted with ProLong™ Gold antifade mountant (Life Technologies: P36930). Antibodies are listed in **table S17**.

### Confocal microscopy

Fluorescent staining of cells and membranes on coverslips was visualized using a Leica SP5 confocal microscope equipped with 40x (numerical aperture (NA) 1.25) and 63x (NA 1.4) oil-immersion objectives and 405/488/594nm lasers. LAS-AF software (v2.7.3) was used to capture Z-series images with a 0.5μm Z-step. Images were processed using ImageJ version 2.1.0/1.53c. This involved maximum intensity projection, background subtraction using ImageJ’s built-in subtraction algorithm based on the “rolling ball” method, and adjustment of contrast and brightness to ensure consistent image quality for all images in the experiment.

### High-content imaging and automated image analysis

High-content imaging was performed on neurons grown on Nunc™ MicroWell™ 96-Well optical-bottom plates with a coverglass base (Thermo Scientific; 164588) using an Opera Phenix High-Content Screening System (PerkinElmer, Waltham, MA) with a 20x water-immersion objective (NA 1.0). Immunofluorescence was captured using confocal imaging with 385nm, 488nm, 561nm and 640nm lasers. An automated imaging script was generated, adjusting exposure times and laser powers for each channel and antibody using negative control wells containing DAPI-stained cells treated with primary or secondary antibodies only. Once generated, scripts remained consistent across all biological replications. Each plate contained neurons from a single clone per genotype. Each condition was imaged in technical triplicate, capturing 15 fields-of-view per well in z-stacks with 3 planes and 1.5µm plane spacing. The acquired images were further processed using the Harmony High Content Imaging and Analysis Software.

An automated image analysis script was developed to analyze synaptic and ribosomal proteins. The script used max-projected images from a z-stack (3 planes) as input and defined regions of interest (ROIs) for separate analysis of protein puncta and mean intensities in somata, neurites, and whole cellular regions. It began by identifying DAPI^+^ cell nuclei and segmenting the MAP2^+^ areas around these nuclei, marking them as soma ROIs. Neurites were defined as ROIs based on the MAP2 channel and an algorithm by the Australian CSIRO research institute (http://www.csiro.au). Whole cell outlines were also marked as ROIs based on the MAP2 channel. The analysis script was tailored to detect puncta (0.4 – 4 µm) and measuring signal mean intensities separately within each ROI. To filter out erroneous puncta detections, a threshold was established using negative control wells to assess the mean intensity of background staining. Only puncta exceeding this threshold value were considered for analysis (**fig. S17**). For co-localisation analysis, the script was instructed to identify positively co-localised puncta by overlaying the PSD95 channel onto the VGLUT1 channel. Puncta were counted as co-localised if a positively identified PSD95 punctum had at least 50% overlap with a positively identified VGLUT1 punctum. Puncta counts for each cellular region and mean intensities of puromycin staining within neurite ROIs were averaged per well and exported to Microsoft Excel for additional analysis.

For SUnSET analyses, FC in puromycin MIs were determined by dividing the MI of the mutant by the MI of the wildtype cells from the same plate. This was done to mitigate the influence of batch effects and technical variability:

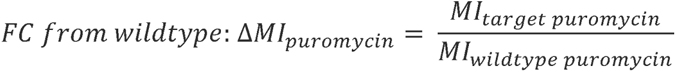

For synaptic and ribosomal proteins, puncta counts were initially normalized to ROI variable parameters, which included normalizing somatic puncta to somata counts, whole-cell puncta to MAP2^+^ cell counts, and neurite puncta to neurite lengths. This normalization process corrected for technical variations in automated analyses, such as varying numbers of detected somata in different wells. Subsequently, FC were computed using the provided formulas (PC = puncta count):

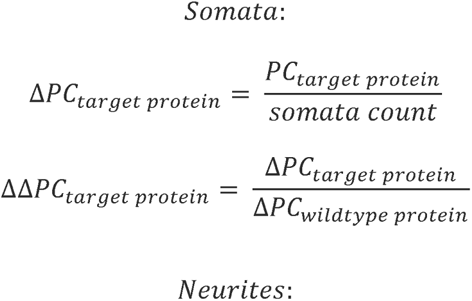

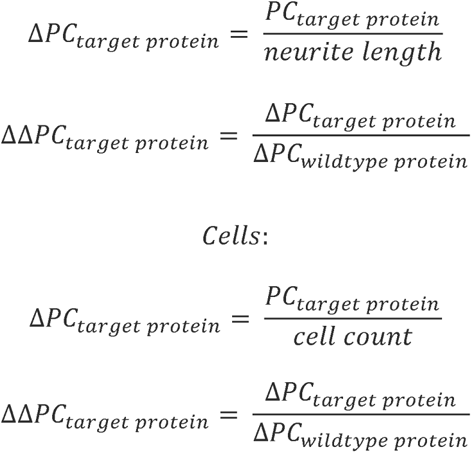

Statistical analysis was performed on averaged technical replicates to yield a single data point per biological replicate. Each biological replicate represented the averaged mean intensities or puncta counts of 2-3 wells. For better representation of data variability technical replicates are plotted in raincloud plots.

### Statistical analysis

All statistics was conducted in RStudio version 1.3.1093. Data normality was assessed through Shapiro-Wilk tests. Parametric tests were applied to normally distributed data, while non-parametric tests were used for data that did not follow a normal distribution. Non-parametric testing was carried out using Kruskal-Wallis tests and Dunn’s post hoc comparisons, while parametric testing was conducted using one-way/two-way ANOVA with TukeyHSD corrections or unpaired Student’s t-tests. Statistical significance was determined based on p-values and adjusted p-values, with <0.05 considered significant. Effect sizes were determined for ANOVA and Kruskal-Wallis tests using R’s integrated functions that computed η^2^. The data was presented graphically through barcharts, boxplots, and raincloud plots using ggplot2. Standard deviation is indicated by error bars on the barcharts.

## Acknowledgments

We express our gratitude to the Wohl Cellular Imaging Centre (WCIC) at the Institute of Psychiatry, Psychology, and Neuroscience (IoPPN), King’s College London, for their valuable assistance in microscopy and the King’s Computational Research, Engineering and Technology Environment (CREATE) for providing a platform to perform the alignment, sorting, and counts table generation of our RNAseq analysis.

## Funding

The study was supported by grants from UK Medical Research Council, Grant No. MR/L021064/1 to DPS; DPS and AVC acknowledge UK Medical Research Council Centre for Neurodevelopmental Disorders (Grant No. MR/N026063/1) and UK Medical Research Council, Cluster Grant No MR/Y012968/1; Royal Society UK (Grant RG130856) to DPS; Independent Researcher Award from the Brain and Behavior Foundation (formally National Alliance for Research on Schizophrenia and Depression (NARSAD) (Grant No. 25957), awarded to DPS; ACV acknowledge financial support for this study from the National Centre for the Replacement, Refinement and Reduction of Animals in Research (NC/S001506/1); LS is supported by the UK Medical Research Council (MR/N013700/1) and King’s College London member of the MRC Doctoral Training Partnership in Biomedical Sciences; RRRD received funds from the Coordenação de Aperfeiçoamento de Pessoal de Nível Superior (CAPES, BEX1279/13-0).

## Author contributions

Conceptualization: ACV, DPS

Methodology: LS, LDP, RRRD, TP, KM, DPS

Investigation: LS, MW, FN, RRRD, DPS

Visualization: LS, DPS

Supervision: ACV, DPS

Writing—original draft: LS, DPS

Writing—review & editing: LS, RRRD, TP, ACV, DPS

## Competing interests

ACV and DPS receive research funding from bit.bio for an unrelated project; DPS receives funding from AstraZeneca for an unrelated project. All other authors declare they have no competing interests.

## Data and materials availability

All data are available in the main text or the supplementary materials or upon direct request from the authors. Proteomic data has been deposited to the ProteomeXchange Consortium via the PRIDE partner repository with the dataset identifier PXD047788. RNASeq data can be found at Gene Expression Omnibus (GEO - https://www.ncbi.nlm.nih.gov/geo/) with accession number: GSE254523.

## References

Aabdien, A., Sichlinger, L., Borgel, Z., Jones, M. R., Waston, I. A., Gatford, N. J. F., Raval, P., Tanangonan, L., Powell, T. R., Duarte, R. R. R., & Srivastava, D. P. (2024). Schizophrenia risk proteins ZNF804A and NT5C2 interact in cortical neurons. Eur J Neurosci. 10.1111/ejn.16254

Adhya, D., Swarup, V., Nagy, R., Dutan, L., Shum, C., Valencia-Alarcón, E. P., Jozwik, K. M., Mendez, M. A., Horder, J., Loth, E., Nowosiad, P., Lee, I., Skuse, D., Flinter, F. A., Murphy, D., McAlonan, G., Geschwind, D. H., Price, J., Carroll, J., … Baron-Cohen, S. (2021). Atypical neurogenesis in induced pluripotent stem cells from autistic individuals. Biological Psychiatry, 89(5), 486–496. 10.1016/j.biopsych.2020.06.014

Anders, S., Pyl, P. T., & Huber, W. (2015). HTSeq — a Python framework to work with high-throughput sequencing data. Bioinformatics, 31(2), 166–169. 10.1093/bioinformatics/btu638

Barkovits, K., Pacharra, S., Pfeiffer, K., Steinbach, S., Eisenacher, M., Marcus, K., & Uszkoreit, J. (2020). Reproducibility, Specificity and Accuracy of Relative Quantification Using Spectral Library-based Data-independent Acquisition. Molecular & Cellular Proteomics, 19(1), 181–197. 10.1074/mcp.RA119.001714

Brennand, K., Savas, J. N., Kim, Y., Tran, N., Simone, A., Hashimoto-Torii, K., Beaumont, K. G., Kim, H. J., Topol, A., Ladran, I., Abdelrahim, M., Matikainen-Ankney, B., Chao, S. h., Mrksich, M., Rakic, P., Fang, G., Zhang, B., Yates, J. R., & Gage, F. H. (2015). Phenotypic differences in hiPSC NPCs derived from patients with schizophrenia. Molecular Psychiatry, 20(3), 361–368. 10.1038/mp.2014.22

Cameron, D., Mi, D., Vinh, N.-N., Webber, C., Li, M., Marín, O., O’Donovan, M. C., & Bray, N. J. (2023). Single-Nuclei RNA Sequencing of 5 Regions of the Human Prenatal Brain Implicates Developing Neuron Populations in Genetic Risk for Schizophrenia. Biological Psychiatry, 93(2), 157–166. 10.1016/j.biopsych.2022.06.033

Chang, H., Xiao, X., & Li, M. (2017). The schizophrenia risk gene ZNF804A: clinical associations, biological mechanisms and neuronal functions. Molecular Psychiatry, 22(7), 944–953. 10.1038/mp.2017.19

Chapman, R. M., Tinsley, C. L., Hill, M. J., Forrest, M. P., Tansey, K. E., Pardiñas, A. F., Rees, E., Doyle, A. M., Wilkinson, L. S., Owen, M. J., O’Donovan, M. C., & Blake, D. J. (2018). Convergent Evidence That ZNF804A Is a Regulator of Pre-messenger RNA Processing and Gene Expression. Schizophrenia Bulletin. 10.1093/schbul/sby183

Chen, J., Lin, M., Hrabovsky, A., Pedrosa, E., Dean, J., Jain, S., Zheng, D., & Lachman, H. M. (2015). ZNF804A Transcriptional Networks in Differentiating Neurons Derived from Induced Pluripotent Stem Cells of Human Origin. Plos One, 10(4), e0124597. 10.1371/journal.pone.0124597

Christiansen, D. M., McCarthy, M. M., & Seeman, M. V. (2022). Where Sex Meets Gender: How Sex and Gender Come Together to Cause Sex Differences in Mental Illness. Frontiers in psychiatry, 13, 856436. 10.3389/fpsyt.2022.856436

Couch, A. C. M., Solomon, S., Duarte, R. R. R., Marrocu, A., Sun, Y., Sichlinger, L., Matuleviciute, R., Polit, L. D., Hanger, B., Brown, A., Kordasti, S., Srivastava, D. P., & Vernon, A. C. (2023). Acute IL-6 exposure triggers canonical IL6Ra signaling in hiPSC microglia, but not neural progenitor cells. Brain, Behavior, and Immunity, 110, 43–59. 10.1016/j.bbi.2023.02.007

Danecek, P., Bonfield, J. K., Liddle, J., Marshall, J., Ohan, V., Pollard, M. O., Whitwham, A., Keane, T., McCarthy, S. A., Davies, R. M., & Li, H. (2021). Twelve years of SAMtools and BCFtools. GigaScience, 10(2). 10.1093/gigascience/giab008

Deans, P. J. M., Raval, P., Sellers, K. J., Gatford, N. J. F., Halai, S., Duarte, R. R. R., Shum, C., Warre-Cornish, K., Kaplun, V. E., Cocks, G., Hill, M., Bray, N. J., Price, J., & Srivastava, D. P. (2017). Psychosis risk candidate ZNF804A localizes to synapses and regulates neurite formation and dendritic spine structure. Biological Psychiatry, 82(1), 49–61. 10.1016/j.biopsych.2016.08.038

Deans, P. M., Seah, C., Johnson, J., Gonzalez, J. G., Townsley, K., Cao, E., Schrode, N., Stahl, E., O’Reilly, P., Huckins, L. M., & Brennand, K. J. (2023). Non-additive effects of schizophrenia risk genes reflect convergent downstream function. medRxiv. 10.1101/2023.03.20.23287497

Dobin, A., Davis, C. A., Schlesinger, F., Drenkow, J., Zaleski, C., Jha, S., Batut, P., Chaisson, M., & Gingeras, T. R. (2013). STAR: ultrafast universal RNA-seq aligner. Bioinformatics, 29(1), 15–21. 10.1093/bioinformatics/bts635

Dobrindt, K., Zhang, H., Das, D., Abdollahi, S., Prorok, T., Ghosh, S., Weintraub, S., Genovese, G., Powell, S. K., Lund, A., Akbarian, S., Eggan, K., McCarroll, S., Duan, J., Avramopoulos, D., & Brennand, K. J. (2021). Publicly Available hiPSC Lines with Extreme Polygenic Risk Scores for Modeling Schizophrenia. Complex Psychiatry, 6(3-4), 68–82. 10.1159/000512716

Dong, F., Mao, J., Chen, M., Yoon, J., & Mao, Y. (2020). Schizophrenia Risk ZNF804A Interacts with its Associated Proteins to Modulate Dendritic Morphology and Synaptic Development. 10.21203/rs.3.rs-113453/v1

Esslinger, C., Walter, H., Kirsch, P., Erk, S., Schnell, K., Arnold, C., Haddad, L., Mier, D., Opitz von Boberfeld, C., Raab, K., Witt, S. H., Rietschel, M., Cichon, S., & Meyer-Lindenberg, A. (2009). Neural mechanisms of a genome-wide supported psychosis variant. Science, 324(5927), 605. 10.1126/science.1167768

Forrest, M. P., Parnell, E., & Penzes, P. (2018). Dendritic structural plasticity and neuropsychiatric disease. Nat Rev Neurosci, 19(4), 215–234. 10.1038/nrn.2018.16

Forsyth, J. K., & Lewis, D. A. (2017). Mapping the Consequences of Impaired Synaptic Plasticity in Schizophrenia through Development: An Integrative Model for Diverse Clinical Features. Trends Cogn Sci, 21(10), 760–778. 10.1016/j.tics.2017.06.006

Gandal, M. J., Zhang, P., Hadjimichael, E., Walker, R. L., Chen, C., Liu, S., Won, H., van Bakel, H., Varghese, M., Wang, Y., Shieh, A. W., Haney, J., Parhami, S., Belmont, J., Kim, M., Moran Losada, P., Khan, Z., Mleczko, J., Xia, Y., … Geschwind, D. H. (2018). Transcriptome-wide isoform-level dysregulation in ASD, schizophrenia, and bipolar disorder. Science, 362(6420). 10.1126/science.aat8127

Grunwald, L.-M., Stock, R., Haag, K., Buckenmaier, S., Eberle, M.-C., Wildgruber, D., Storchak, H., Kriebel, M., Weißgraeber, S., Mathew, L., Singh, Y., Loos, M., Li, K. W., Kraushaar, U., Fallgatter, A. J., & Volkmer, H. (2019). Comparative characterization of human induced pluripotent stem cells (hiPSC) derived from patients with schizophrenia and autism. Translational psychiatry, 9(1), 179. 10.1038/s41398-019-0517-3

Gusev, A., Mancuso, N., Won, H., Kousi, M., Finucane, H. K., Reshef, Y., Song, L., Safi, A., Schizophrenia Working Group of the Psychiatric Genomics, C., McCarroll, S., Neale, B. M., Ophoff, R. A., O’Donovan, M. C., Crawford, G. E., Geschwind, D. H., Katsanis, N., Sullivan, P. F., Pasaniuc, B., & Price, A. L. (2018). Transcriptome-wide association study of schizophrenia and chromatin activity yields mechanistic disease insights. NatureGenetics, 50(4), 538–548. 10.1038/s41588-018-0092-1

Hill, M. J., & Bray, N. J. (2012). Evidence that schizophrenia risk variation in the ZNF804A gene exerts its effects during fetal brain development. The American Journal of Psychiatry, 169(12), 1301–1308. 10.1176/appi.ajp.2012.11121845

Hill, M. J., Jeffries, A. R., Dobson, R. J. B., Price, J., & Bray, N. J. (2012). Knockdown of the psychosis susceptibility gene ZNF804A alters expression of genes involved in cell adhesion. Human Molecular Genetics, 21(5), 1018–1024. 10.1093/hmg/ddr532

Huang, Y., Huang, J., Zhou, Q.-X., Yang, C.-X., Yang, C.-P., Mei, W.-Y., Zhang, L., Zhang, Q., Hu, L., Hu, Y.-Q., Song, N.-N., Wu, S.-X., Xu, L., & Ding, Y.-Q. (2020). ZFP804A mutant mice display sex-dependent schizophrenia-like behaviors. Molecular Psychiatry. 10.1038/s41380-020-00972-4

Huber, K. M., Kayser, M. S., & Bear, M. F. (2000). Role for rapid dendritic protein synthesis in hippocampal mGluR-dependent long-term depression. Science, 288(5469), 1254–1257. 10.1126/science.288.5469.1254

Ishii, T., Ishikawa, M., Fujimori, K., Maeda, T., Kushima, I., Arioka, Y., Mori, D., Nakatake, Y., Yamagata, B., Nio, S., Kato, T. A., Yang, N., Wernig, M., Kanba, S., Mimura, M., Ozaki, N., & Okano, H. (2019). In Vitro Modeling of the Bipolar Disorder and Schizophrenia Using Patient-Derived Induced Pluripotent Stem Cells with Copy Number Variations of PCDH15 and RELN. eNeuro, 6(5). 10.1523/ENEURO.0403-18.2019

Kang, H., & Schuman, E. M. (1996). A requirement for local protein synthesis in neurotrophin-induced hippocampal synaptic plasticity. Science, 273(5280), 1402–1406. 10.1126/science.273.5280.1402

Klockmeier, K., Silva Ramos, E., Rasko, T., Marti Pastor, A., & Wanker, E. E. (2021). Schizophrenia risk candidate protein ZNF804A interacts with STAT2 and influences interferon-mediated gene transcription in mammalian cells. J Mol Biol, 433(19), 167184. 10.1016/j.jmb.2021.167184

Lencz, T., Szeszko, P. R., DeRosse, P., Burdick, K. E., Bromet, E. J., Bilder, R. M., & Malhotra, A. K. (2010). A schizophrenia risk gene, ZNF804A, influences neuroanatomical and neurocognitive phenotypes. Neuropsychopharmacology, 35(11), 2284–2291. 10.1038/npp.2010.102

Maroof, A. M., Keros, S., Tyson, J. A., Ying, S.-W., Ganat, Y. M., Merkle, F. T., Liu, B., Goulburn, A., Stanley, E. G., Elefanty, A. G., Widmer, H. R., Eggan, K., Goldstein, P. A., Anderson, S. A., & Studer, L. (2013). Directed differentiation and functional maturation of cortical interneurons from human embryonic stem cells. Cell Stem Cell, 12(5), 559–572. 10.1016/j.stem.2013.04.008

Mitsumori, K., Takei, Y., & Hirokawa, N. (2017). Components of RNA granules affect their localization and dynamics in neuronal dendrites. Molecular Biology of the Cell, 28(11), 1412–1417. 10.1091/mbc.E16-07-0497

Murray, R. M., Bora, E., Modinos, G., & Vernon, A. (2022). Schizophrenia: A developmental disorder with a risk of non-specific but avoidable decline. Schizophrenia Research, 243, 181–186. 10.1016/j.schres.2022.03.005

Nehme, R., Zuccaro, E., Ghosh, S. D., Li, C., Sherwood, J. L., Pietilainen, O., Barrett, L. E., Limone, F., Worringer, K. A., Kommineni, S., Zang, Y., Cacchiarelli, D., Meissner, A., Adolfsson, R., Haggarty, S., Madison, J., Muller, M., Arlotta, P., Fu, Z., … Eggan, K. (2018). Combining NGN2 Programming with Developmental Patterning Generates Human Excitatory Neurons with NMDAR-Mediated Synaptic Transmission. Cell reports, 23(8), 2509–2523. 10.1016/j.celrep.2018.04.066

O’Donovan, M. C., Craddock, N., Norton, N., Williams, H., Peirce, T., Moskvina, V., Nikolov, I., Hamshere, M., Carroll, L., Georgieva, L., Dwyer, S., Holmans, P., Marchini, J. L., Spencer, C. C. A., Howie, B., Leung, H.-T., Hartmann, A. M., Möller, H.-J., Morris, D. W., … Molecular Genetics of Schizophrenia, C. (2008). Identification of loci associated with schizophrenia by genome-wide association and follow-up. Nature Genetics, 40(9), 1053–1055. 10.1038/ng.201

Pavlinek, A., Matuleviciute, R., Sichlinger, L., Dutan Polit, L., Armeniakos, N., Vernon, A. C., & Srivastava, D. P. (2022). Interferon-γ exposure of human iPSC-derived neurons alters major histocompatibility complex I and synapsin protein expression. Frontiers in psychiatry, 13, 836217. 10.3389/fpsyt.2022.836217

Pawlowski, M., Ortmann, D., Bertero, A., Tavares, J. M., Pedersen, R. A., Vallier, L., & Kotter, M. R. N. (2017). Inducible and Deterministic Forward Programming of Human Pluripotent Stem Cells into Neurons, Skeletal Myocytes, and Oligodendrocytes. Stem cell reports, 8(4), 803–812. 10.1016/j.stemcr.2017.02.016

Penzes, P., Cahill, M. E., Jones, K. A., VanLeeuwen, J.-E., & Woolfrey, K. M. (2011). Dendritic spine pathology in neuropsychiatric disorders. Nature Neuroscience, 14(3), 285–293. 10.1038/nn.2741

Perez-Riverol, Y., Bai, J., Bandla, C., García-Seisdedos, D., Hewapathirana, S., Kamatchinathan, S., Kundu, D. J., Prakash, A., Frericks-Zipper, A., Eisenacher, M., Walzer, M., Wang, S., Brazma, A., & Vizcaíno, J. A. (2022). The PRIDE database resources in 2022: a hub for mass spectrometry-based proteomics evidences. Nucleic Acids Research, 50(D1), D543–D552. 10.1093/nar/gkab1038

Pirbhoy, P. S., Farris, S., & Steward, O. (2016). Synaptic activation of ribosomal protein S6 phosphorylation occurs locally in activated dendritic domains. Learning & Memory, 23(6), 255–269. 10.1101/lm.041947.116

Polioudakis, D., de la Torre-Ubieta, L., Langerman, J., Elkins, A. G., Shi, X., Stein, J. L., Vuong, C. K., Nichterwitz, S., Gevorgian, M., Opland, C. K., Lu, D., Connell, W., Ruzzo, E. K., Lowe, J. K., Hadzic, T., Hinz, F. I., Sabri, S., Lowry, W. E., Gerstein, M. B., … Geschwind, D. H. (2019). A Single-Cell Transcriptomic Atlas of Human Neocortical Development during Mid-gestation. Neuron, 103(5), 785–801.e788. 10.1016/j.neuron.2019.06.011

Puighermanal, E., Biever, A., Pascoli, V., Melser, S., Pratlong, M., Cutando, L., Rialle, S., Severac, D., Boubaker-Vitre, J., Meyuhas, O., Marsicano, G., Lüscher, C., & Valjent, E. (2017). Ribosomal Protein S6 Phosphorylation Is Involved in Novelty-Induced Locomotion, Synaptic Plasticity and mRNA Translation. Frontiers in Molecular Neuroscience, 10, 419. 10.3389/fnmol.2017.00419

Räsänen, N., Tiihonen, J., Koskuvi, M., Lehtonen, Š., & Koistinaho, J. (2022). The iPSC perspective on schizophrenia. Trends in Neurosciences, 45(1), 8–26. 10.1016/j.tins.2021.11.002

Rasetti, R., Sambataro, F., Chen, Q., Callicott, J. H., Mattay, V. S., & Weinberger, D. R. (2011). Altered cortical network dynamics: a potential intermediate phenotype for schizophrenia and association with ZNF804A. Arch Gen Psychiatry, 68(12), 1207–1217. 10.1001/archgenpsychiatry.2011.103

Ritchie, M. E., Phipson, B., Wu, D., Hu, Y., Law, C. W., Shi, W., & Smyth, G. K. (2015). limma powers differential expression analyses for RNA-sequencing and microarray studies. Nucleic Acids Research, 43(7), e47. 10.1093/nar/gkv007

Robicsek, O., Karry, R., Petit, I., Salman-Kesner, N., Müller, F. J., Klein, E., Aberdam, D., & Ben-Shachar, D. (2013). Abnormal neuronal differentiation and mitochondrial dysfunction in hair follicle-derived induced pluripotent stem cells of schizophrenia patients. Molecular Psychiatry, 18(10), 1067–1076. 10.1038/mp.2013.67

Ruvinsky, I., & Meyuhas, O. (2006). Ribosomal protein S6 phosphorylation: from protein synthesis to cell size. Trends in Biochemical Sciences, 31(6), 342–348. 10.1016/j.tibs.2006.04.003

Sando, R., Bushong, E., Zhu, Y., Huang, M., Considine, C., Phan, S., Ju, S., Uytiepo, M., Ellisman, M., & Maximov, A. (2017). Assembly of excitatory synapses in the absence of glutamatergic neurotransmission. Neuron, 94(2), 312–321.e313. 10.1016/j.neuron.2017.03.047

Schizophrenia Working Group of the Psychiatric Genomics, C. (2014). Biological insights from 108 schizophrenia-associated genetic loci. Nature, 511(7510), 421–427. 10.1038/nature13595

Schmidt, E. K., Clavarino, G., Ceppi, M., & Pierre, P. (2009). SUnSET, a nonradioactive method to monitor protein synthesis. Nature Methods, 6(4), 275–277. 10.1038/nmeth.1314

Schrode, N., Ho, S. M., Yamamuro, K., Dobbyn, A., Huckins, L., Matos, M. R., Cheng, E., Deans, P. J. M., Flaherty, E., Barretto, N., Topol, A., Alganem, K., Abadali, S., Gregory, J., Hoelzli, E., Phatnani, H., Singh, V., Girish, D., Aronow, B., … Brennand, K. J. (2019). Synergistic effects of common schizophrenia risk variants. Nature Genetics, 51(10), 1475–1485. 10.1038/s41588-019-0497-5

Sey, N. Y. A., Hu, B., Mah, W., Fauni, H., McAfee, J. C., Rajarajan, P., Brennand, K. J., Akbarian, S., & Won, H. (2020). A computational tool (H-MAGMA) for improved prediction of brain-disorder risk genes by incorporating brain chromatin interaction profiles. Nature Neuroscience, 23(4), 583–593. 10.1038/s41593-020-0603-0

Sherman, B. T., Hao, M., Qiu, J., Jiao, X., Baseler, M. W., Lane, H. C., Imamichi, T., & Chang, W. (2022). DAVID: a web server for functional enrichment analysis and functional annotation of gene lists (2021 update). Nucleic Acids Research, 50(W1), W216–W221. 10.1093/nar/gkac194

Shum, C., Dutan, L., Annuario, E., Warre-Cornish, K., Taylor, S. E., Taylor, R. D., Andreae, L. C., Buckley, N. J., Price, J., Bhattacharyya, S., & Srivastava, D. P. (2020). Δ9-tetrahydrocannabinol and 2-AG decreases neurite outgrowth and differentially affects ERK1/2 and Akt signaling in hiPSC-derived cortical neurons. Molecular and Cellular Neurosciences, 103, 103463. 10.1016/j.mcn.2019.103463

Sigler, A., Oh, W. C., Imig, C., Altas, B., Kawabe, H., Cooper, B. H., Kwon, H.-B., Rhee, J.-S., & Brose, N. (2017). Formation and maintenance of functional spines in the absence of presynaptic glutamate release. Neuron, 94(2), 304–311.e304. 10.1016/j.neuron.2017.03.029

Singh, T., Poterba, T., Curtis, D., Akil, H., Al Eissa, M., Barchas, J. D., Bass, N., Bigdeli, T. B., Breen, G., Bromet, E. J., Buckley, P. F., Bunney, W. E., Bybjerg-Grauholm, J., Byerley, W. F., Chapman, S. B., Chen, W. J., Churchhouse, C., Craddock, N., Cusick, C. M., … Daly, M. J. (2022). Rare coding variants in ten genes confer substantial risk for schizophrenia. Nature, 604(7906), 509–516. 10.1038/s41586-022-04556-w

Skene, N. G., Bryois, J., Bakken, T. E., Breen, G., Crowley, J. J., Gaspar, H. A., Giusti-Rodriguez, P., Hodge, R. D., Miller, J. A., Muñoz-Manchado, A. B., O’Donovan, M. C., Owen, M. J., Pardiñas, A. F., Ryge, J., Walters, J. T. R., Linnarsson, S., Lein, E. S., Major Depressive Disorder Working Group of the Psychiatric Genomics, C., Sullivan, P. F., & Hjerling-Leffler, J. (2018). Genetic identification of brain cell types underlying schizophrenia. Nature Genetics, 50(6), 825–833. 10.1038/s41588-018-0129-5

Sriretnakumar, V., Zai, C. C., Wasim, S., Barsanti-Innes, B., Kennedy, J. L., & So, J. (2019). Copy number variant syndromes are frequent in schizophrenia: Progressing towards a CNV-schizophrenia model. Schizophrenia Research, 209, 171–178. 10.1016/j.schres.2019.04.026

Südhof, T. C. (2018). Towards an understanding of synapse formation. Neuron, 100(2), 276–293. 10.1016/j.neuron.2018.09.040

Sullivan, P. F., Daly, M. J., & O’Donovan, M. (2012). Genetic architectures of psychiatric disorders: the emerging picture and its implications. Nature Reviews. Genetics, 13(8), 537–551. 10.1038/nrg3240

Tao, R., Cousijn, H., Jaffe, A. E., Burnet, P. W. J., Edwards, F., Eastwood, S. L., Shin, J. H., Lane, T. A., Walker, M. A., Maher, B. J., Weinberger, D. R., Harrison, P. J., Hyde, T. M., & Kleinman, J. E. (2014). Expression of ZNF804A in human brain and alterations in schizophrenia, bipolar disorder, and major depressive disorder: a novel transcript fetally regulated by the psychosis risk variant rs1344706. JAMA psychiatry, 71(10), 1112–1120. 10.1001/jamapsychiatry.2014.1079

Treutlein, B., Lee, Q. Y., Camp, J. G., Mall, M., Koh, W., Shariati, S. A. M., Sim, S., Neff, N. F., Skotheim, J. M., Wernig, M., & Quake, S. R. (2016). Dissecting direct reprogramming from fibroblast to neuron using single-cell RNA-seq. Nature, 534(7607), 391–395. 10.1038/nature18323

Trubetskoy, V., Pardiñas, A. F., Qi, T., Panagiotaropoulou, G., Awasthi, S., Bigdeli, T. B., Bryois, J., Chen, C.-Y., Dennison, C. A., Hall, L. S., Lam, M., Watanabe, K., Frei, O., Ge, T., Harwood, J. C., Koopmans, F., Magnusson, S., Richards, A. L., Sidorenko, J., … Schizophrenia Working Group of the Psychiatric Genomics, C. (2022). Mapping genomic loci implicates genes and synaptic biology in schizophrenia. Nature, 604(7906), 502–508. 10.1038/s41586-022-04434-5

UniProt, C. (2023). Uniprot: the universal protein knowledgebase in 2023. Nucleic Acids Research, 51(D1), D523–D531. 10.1093/nar/gkac1052

Verhage, M., Maia, A. S., Plomp, J. J., Brussaard, A. B., Heeroma, J. H., Vermeer, H., Toonen, R. F., Hammer, R. E., van den Berg, T. K., Missler, M., Geuze, H. J., & Südhof, T. C. (2000). Synaptic assembly of the brain in the absence of neurotransmitter secretion. Science, 287(5454), 864–869. 10.1126/science.287.5454.864

Wingett, S. W., & Andrews, S. (2018). FastQ Screen: A tool for multi-genome mapping and quality control. F1000Research, 7, 1338. 10.12688/f1000research.15931.2

Wingo, A. P., Liu, Y., Gerasimov, E. S., Vattathil, S. M., Liu, J., Cutler, D. J., Epstein, M. P., Blokland, G. A. M., Thambisetty, M., Troncoso, J. C., Duong, D. M., Bennett, D. A., Levey, A. I., Seyfried, N. T., & Wingo, T. S. (2023). Sex differences in brain protein expression and disease. Nat Med, 29(9), 2224–2232. 10.1038/s41591-023-02509-y

Wong, H. H.-W., Lin, J. Q., Ströhl, F., Roque, C. G., Cioni, J.-M., Cagnetta, R., Turner-Bridger, B., Laine, R. F., Harris, W. A., Kaminski, C. F., & Holt, C. E. (2017). RNA Docking and Local Translation Regulate Site-Specific Axon Remodeling In Vivo. Neuron, 95(4), 852–868.e858. 10.1016/j.neuron.2017.07.016

Wong, H. H.-W., Watt, A. J., & Sjöström, P. J. (2023). Synapse-specific burst coding sustained by local axonal translation. Neuron. 10.1016/j.neuron.2023.10.011

Zappulo, A., van den Bruck, D., Ciolli Mattioli, C., Franke, V., Imami, K., McShane, E., Moreno-Estelles, M., Calviello, L., Filipchyk, A., Peguero-Sanchez, E., Müller, T., Woehler, A., Birchmeier, C., Merino, E., Rajewsky, N., Ohler, U., Mazzoni, E. O., Selbach, M., Akalin, A., & Chekulaeva, M. (2017). RNA localization is a key determinant of neurite-enriched proteome. Nature Communications, 8(1), 583. 10.1038/s41467-017-00690-6

Zhang, F., Chen, Q., Ye, T., Lipska, B. K., Straub, R. E., Vakkalanka, R., Rujescu, D., St Clair, D., Hyde, T. M., Bigelow, L., Kleinman, J. E., & Weinberger, D. R. (2011). Evidence of sex-modulated association of ZNF804A with schizophrenia. Biological Psychiatry, 69(10), 914–917. 10.1016/j.biopsych.2011.01.003

Zhou, D., Xiao, X., & Li, M. (2020). The schizophrenia risk isoform ZNF804AE3E4 affects dendritic spine. Schizophrenia Research. 10.1016/j.schres.2019.12.038

Zhou, Y., Dong, F., Lanz, T. A., Reinhart, V., Li, M., Liu, L., Zou, J., Xi, H. S., & Mao, Y. (2018). Interactome analysis reveals ZNF804A, a schizophrenia risk gene, as a novel component of protein translational machinery critical for embryonic neurodevelopment. Molecular Psychiatry, 23(4), 952–962. 10.1038/mp.2017.166

