## Supplemental Material for "Schizophrenia risk gene ZNF804A controls ribosome localization and synaptogenesis in developing human neurons"

**This PDF file includes:**

Supplemental methods
Supplemental text
Supplemental figures - S1 to S17
Tables S1 to S17

#### Supplemental Methods

##### Genotyping

Initial genotyping to identify potential CRISPR/Cas9 candidates involved PCR amplification using primers flanking the excision site, and DNA from crude cell lysates. A total of 96 clones were prepared for genotyping, where 20µl of each clone's cell suspension was placed in individual PCR tubes, and the remaining 40µl were evenly divided into two pre-warmed 96-well plates. Initial genotyping involved treating each clone's 20µl cell suspension with a PBS-based solution containing 0.5µl of proteinase K (20mg/ml; Promega; A505B), allowing cell lysis and genomic DNA (gDNA) extraction at 56°C for 1h, followed by stopping the reaction at 95°C for 10min, and using 1µl of crude lysates for PCR. 15 promising clones were expanded and underwent final genotyping by extracting gDNA using Promega ReliaPrep™ gDNA Tissue Miniprep System (A2051). To validate genotypes, PCRs were performed on extracted gDNAs from clones using two distinct primer pairs (**table S1**) and CloneAmp HiFi Polymerase (Takara; 639298).

##### Sanger sequencing

To confirm the precise excision sites, 5µl of PCR products and 5µl of forward primers (final concentration: 0.3µM; **table S1**) were submitted to Eurofins Genomics LLC or Source BioScience, and genotype was confirmed by aligning sequences to the amplicon sequence using the Clustal Omega online tool (https://www.ebi.ac.uk/Tools/msa/clustalo/). Sequencing was performed in duplicate using two distinct forward primers to validate the excision sites or wildtype genotypes.

##### Off-target analysis

Potential off-target sites for both sgRNAs were evaluated using IDT's CRISPR/Cas9 guide RNA design checker (www.eu.idtdna.com/site/order/designtool/). The top four sequences with the highest off-target cleavage risk were selected for further examination. Primers (**table S1**) were designed to target regions located approximately 250-850bp both up- and downstream of predicted off-target sites. PCR and Sanger sequencing were carried out as above.

**Real-time quantitative PCR**

RNA samples were collected in TRIzol reagent (Thermo Fisher Scientific, 15596026) and RNA was extracted using chloroform (372978, Sigma) and isopropanol as described before. RNA purity was ensured through sodium acetate (Sigma, S2889) and 100% molecular grade ethanol treatments overnight, and the RNA samples were resuspended in RNase/DNase-free water and stored at -80°C. To analyze ZNF804A gene expression in NPCs, RNA was extracted at seven timepoints, totaling 92 samples, which included NPCs derived from three hiPSC lines and three clones per line (except for 014_01, which lacked a D20 sample due to technical issues). The mRNA extraction was carried out in nine batches, each time using samples from different donors to control for potential batch effects. mRNA from D7 neurons (n = 6) and CRISPR/Cas9 edited lines (n = 18) was extracted in two batches on the same day.

Reverse transcription to produce complementary DNA was conducted using SuperScript III (Invitrogen, 18080-044) and RT-qPCR was performed using MicroAmpTM Optical 384-well reaction plates (4343814, Applied Biosystems) and 5x HOT FIREPol® EvaGreen®

qPCR Mix Plus (ROX) (final concentration 1X; Solis BioDyne 08-24-00020) and gene expression was measured by a QuantStudio7 Flex RT-qPCR system and QuantStudio RT-PCR software (Thermo Scientific, v1.3), based on a protocol optimised for the dyes and primers used (**table S1**). Each condition was assessed with three technical replicates, and the raw data for the cycle threshold (Ct) was analyzed using the $2^{-\Delta\Delta Ct}$ method to calculate the FC in gene expression relative to the control conditions (*54*).

#### Supplementary Text

##### Off-target analysis

We evaluated potential off-target effects. We confirmed pluripotency and self-renewal properties in the selected clones using RT-qPCR and immunocytochemistry (ICC) along with confocal microscopy (**fig. S6**). Using *in silico* modeling to identify potential CRISPR/Cas9 off-target sites for both sgRNAs (**table S2**), we selected the four most high-risk sequences for each sgRNA, isolated genomic DNA from the hiPSCs of each clone, and conducted Sanger sequencing of genomic regions spanning the potential off-target sites. We identified point mutations at the Cas9 cleavage site in off-target sequence 1 of sgRNA1 and off-target sequence 2 of sgRNA2. Specifically, within off-target sequence 1 of sgRNA1 cytosine was substituted for thymine (C → T) and within off-target sequence 2 of sgRNA2 thymine was substituted for guanine (T → G). Interestingly, these off-target effects were observed in all six clones (**fig. S7**). Genomic sequences located both up- and downstream of each off-target substitution displayed no irregularities. In-depth analysis of off-target sequence 1 of sgRNA1 indicated that it resides within a sizable non-coding region on chromosome 1, with the nearest coding region positioned over 17k nucleotides downstream (ring finger protein 220) and more than 32k nucleotides upstream (exoribonuclease family member 3) of the point mutation. Similarly, off-target sequence 2 of sgRNA2 is located in a vast intronic region on chromosome 20, with the closest downstream coding region being over 600k nucleotides away (Jagged1) and the nearest upstream coding region positioned more than 500k nucleotides away (BTB domain containing 3). Considering the significant distances to coding regions, the isolated editing effects, and the uniformity of the off-target effects among all clones, we proceeded with further investigations.

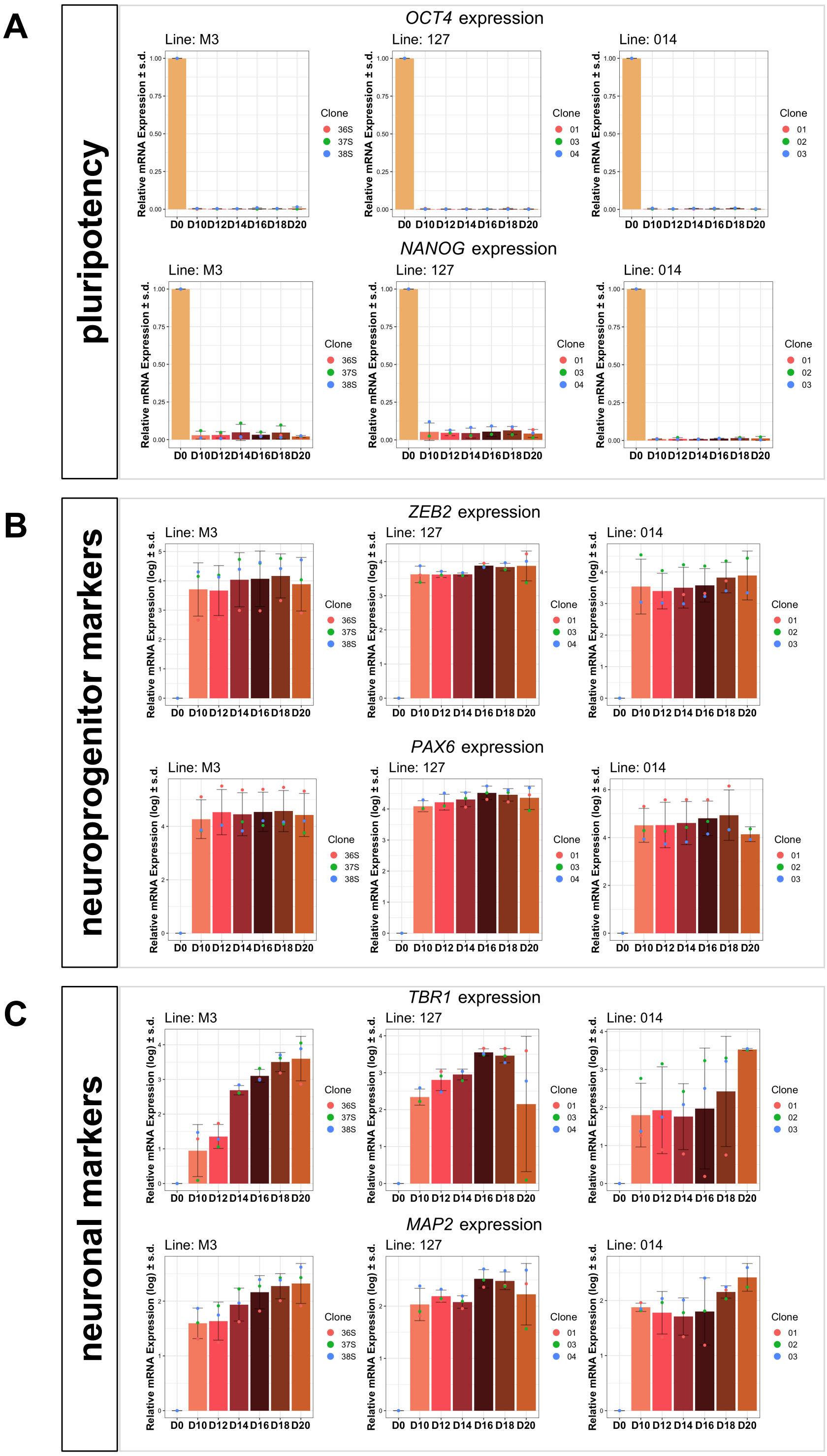

**Fig. S1 Fate marker expression of cell lines at different developmental timepoints.** Barcharts of RT-qPCR-derived log10-transformed $2^{-\Delta\Delta Ct}$ of (A) pluripotency markers octamer-binding transcription factor 4 (*OCT4*) and homeobox protein NANOG (*NANOG*), (B) neuroprogenitor cell (NPC) markers zinc finger E-box binding homeobox 2 (*ZEB2*)

and paired box 6 (*PAX6*), and (C) neuronal markers T-box brain transcription factor 1 (*TBR1*) and microtubule associated protein 2 (*MAP2*) in each cell line used in this study (M3, 127 and 014) at seven timepoints between days (D)0-D20. Overlaying dotplots show colour-coded clones. Cells show expression profiles of NPCs appropriate to their developmental timepoint.
Error bars: mean ± standard deviation (SD)

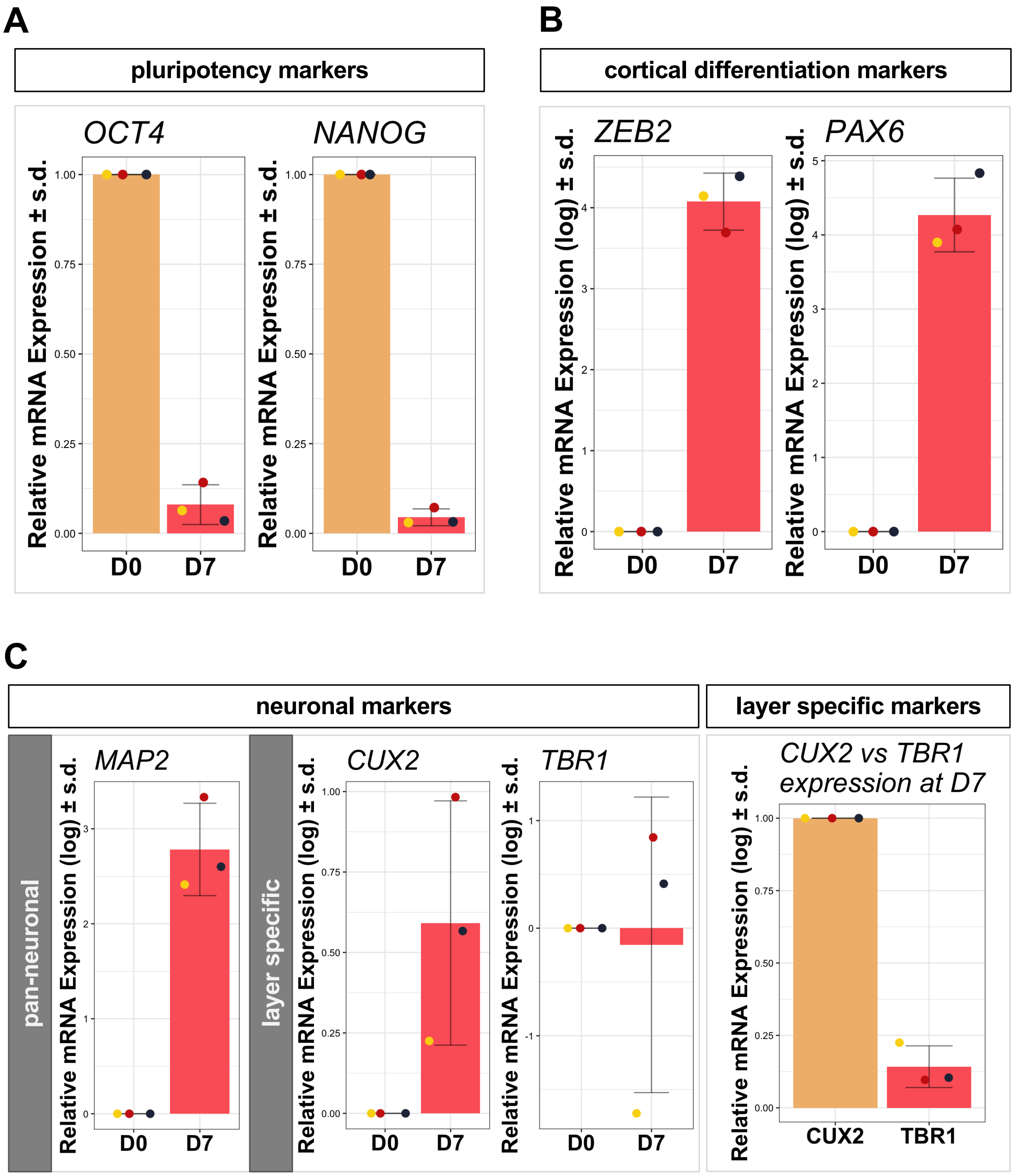

**Fig. S2 ioGlutamatergic cell-derived neurons resemble immature upper layer cortical neurons.** Barcharts of RT-qPCR-derived log10-transformed $2^{-\Delta\Delta Ct}$ of (A) pluripotency markers octamer-binding transcription factor 4 (*OCT4*) and homeobox protein NANOG (*NANOG*) (B) neuroprogenitor cell (NPC) markers zinc finger E-box binding homeobox 2 (*ZEB2*) and paired box 6 (*PAX6*)*,* and (C) pan-neuronal marker microtubule associated protein 2 (*MAP2*), layer VI specific marker T-box brain transcription factor 1 (*TBR1*), and upper layer specific maker cut like homebox 2 (*CUX2*) in human induced pluripotent stem cells (hiPSCs, D0) and developing neurons seven days post neural induction (D7). Biological replicates (i.e. distinct differentiations) are plotted in overlaying dotplots. High *MAP2* transcript abundance at D7 indicates post-mitotic neuronal fate. Biological replicates showed variable layer-specific marker expression; *CUX2* expression was more abundant compared to *TBR1* expression at D7. Error bars: mean ± standard deviation (SD)

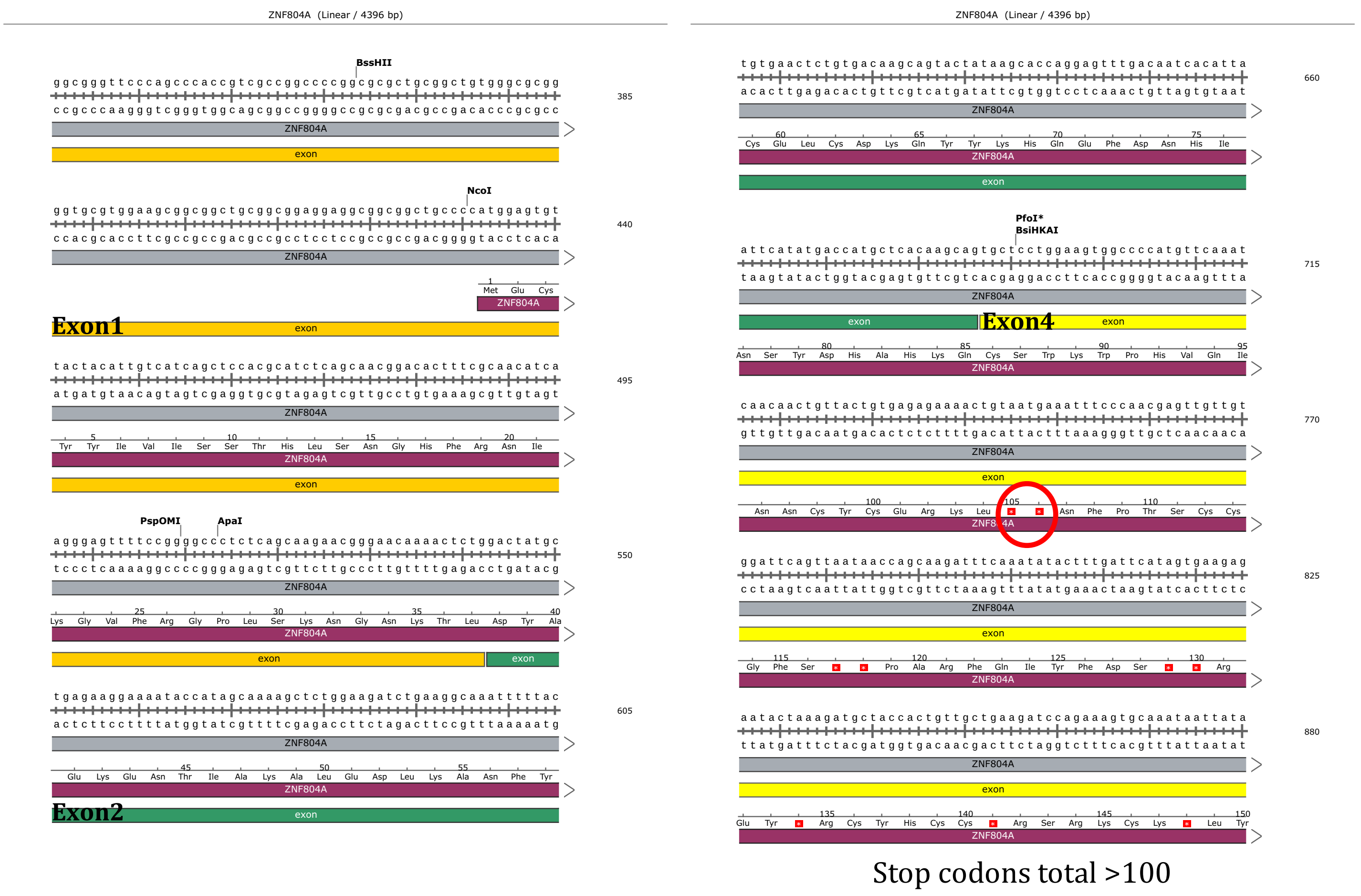

**Fig. S3. *In silico* design of mutant *ZNF804A* model.** SnapGene (www.snapgene.com) modelling of exon 3 excision of *ZNF804A* showing putative premature stop codons in exon 4 (first stop condons marked with red circle; in total > 100).

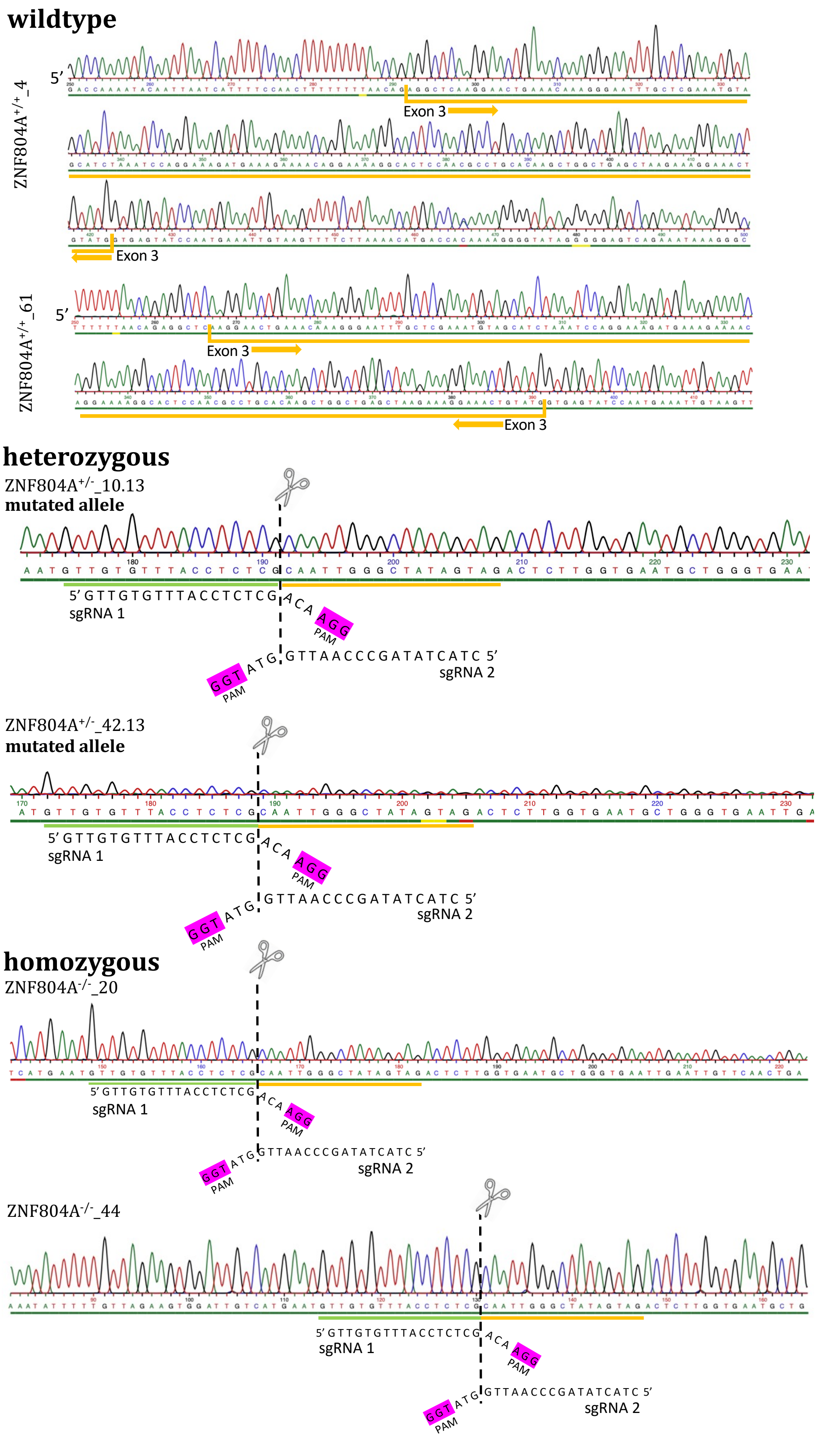

**Fig. S4 Sanger sequences of *ZNF804A* mutant clones.** Exon 3 excision (heterozygous.

ZNF804A$^{+/-}$_10.23, ZNF804A$^{+/-}$_42.13; homozygous: ZNF804A$^{-/-}$_44, ZNF804A$^{-/-}$_20.) or unaltered genotype (wildtype clones ZNF804A$^{+/+}$_4, ZNF804A$^{+/+}$_61) was confirmed by sanger sequencing with two distinct forward primers.

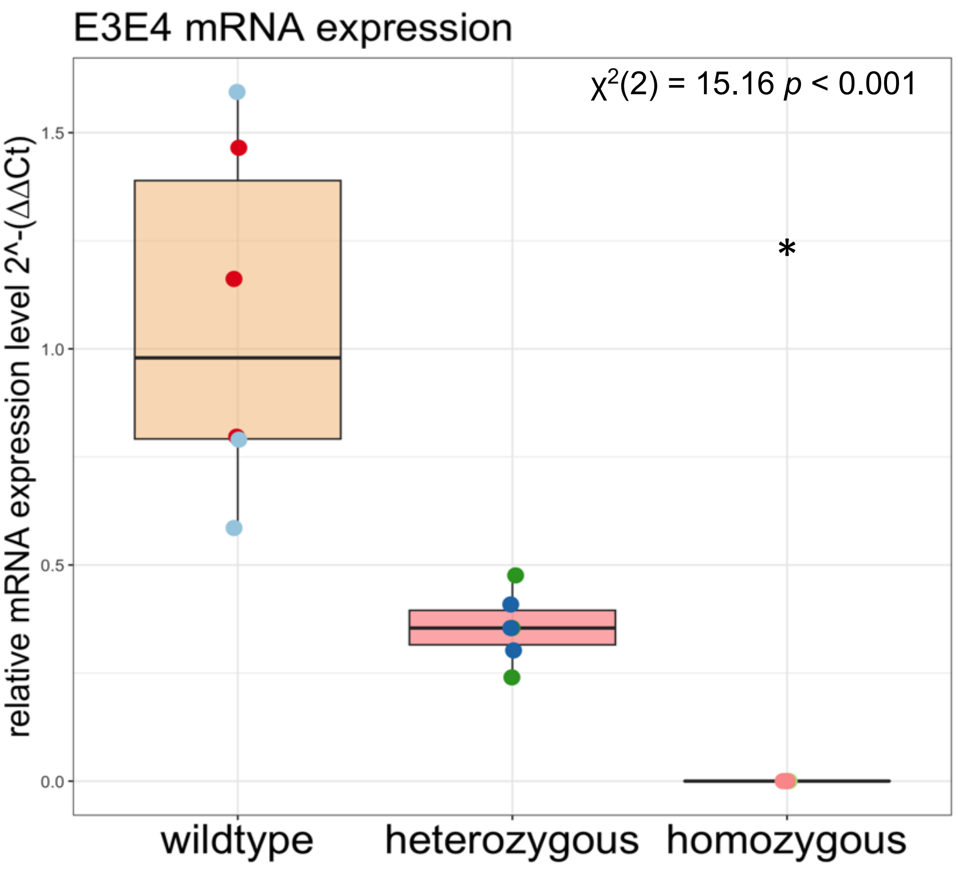

**Fig. S5 mRNA validation of ZNF804A mutant clones.** Boxplots showing RT-qPCR-derived log10-transformed $2^{-\Delta\Delta Ct}$ values of schizophrenia risk variant of zinc-finger protein 804A (*ZNF804A*), *ZNF804A* $^{E3E4}$, in each clone of three genotypes (ZNF804A$^{+/+}$, ZNF804A$^{+/-}$, ZNF804A$^{-/-}$) in D7 developing glutamatergic neurons (n = 18; 3 individual differentiations per clone). Clones are plotted in overlaying dotplots. * p.adj. < 0.05.

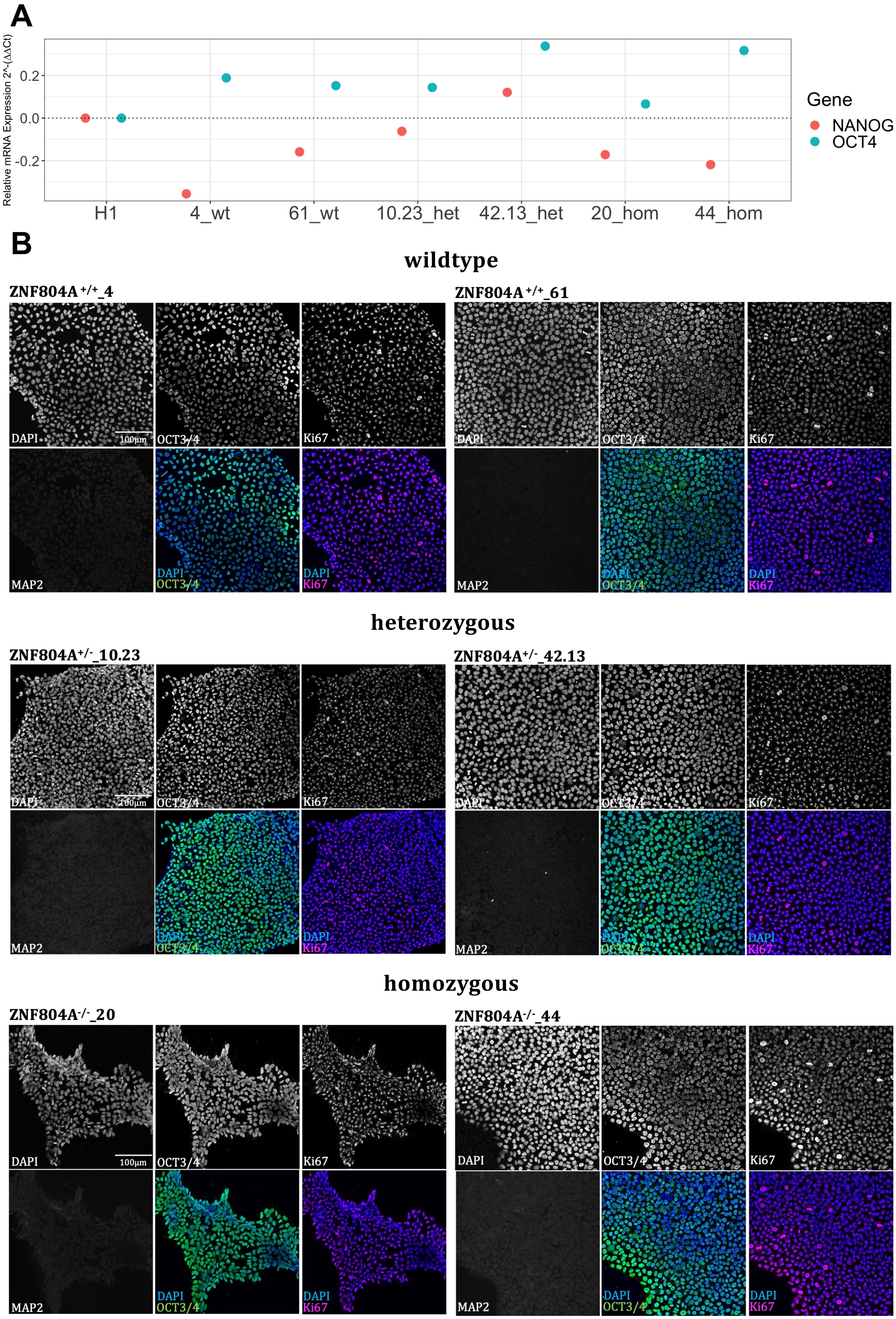
A
Relative mRNA Expression 2^-(ΔΔCt)
0.2
0.0
-0.2
H1
4_wt
61_wt
10.23_het
42.13_het
20_hom
44_hom
Gene
NANOG
OCT4
B
wildtype
ZNF804A +/+_4
ZNF804A +/+_61
DAPI
100μm
OCT3/4
Ki67
MAP2
heterozygous
ZNF804A+/-_10.23
ZNF804A+/-_42.13
homozygous
ZNF804A-/-_20
ZNF804A-/-_44

**Fig. S6 Genome editing did not interfere with pluripotency properties.** (A) Dotplot of RT-qPCR-derived $2^{-\Delta\Delta Ct}$ of values pluripotency markers octamer-binding transcription factor 4 (*OCT4*) and homeobox protein NANOG (*NANOG*) in human induced pluripotent stem cells (hiPSCs) of each genome edited (heterozygous: ZNF804A$^{+/-}$_10.23, ZNF804A$^{+/-}$_42.13; homozygous: ZNF804A$^{-/-}$_44, ZNF804A$^{-/-}$_20.) or wildtype clone (wildtype: ZNF804A$^{+/+}$_4, ZNF804A$^{+/+}$_61) compared to the expression in a male human embryonic stem cell line (hESC – H1). High expression of pluripotency markers, similar to H1 line, indicate clones retained pluripotency properties. (B) Representative confocal images of hiPSCs from each clone. Immunostaining shows pluripotency marker OCT3/4 (green), proliferation marker Ki67 (magenta), and pan-neuronal marker microtubule-associated protein 2 (MAP2). Nuclei were stained with DAPI (blue). Global expression of OCT3/4 and Ki67 confirm unaltered hiPSC properties. MAP2 was not expressed.

| sgRNA sequence 1 - Intron 2<br>**GTTGTGTTTACCTCTCGACA** | | |
|---|---|---|
| **Potential off-target sequence** | | **Locus** |
| 1 | *AATGTGTGTACCTCTTGACA* | chr1:-44387600 |
| 2 | *TTTCTGTTTACCTCTCTACA* | chr7:+122403326 |
| 3 | *ATTGTGTGTGCCTCTTGACA* | chr5:+110971250 |
| 4 | *CTTGAGTTTTCCTCTCTACA* | chr3:-117476849 |

| sgRNA sequence 2 - Intron 3<br>**CTACTATAGCCCAATTGGTA** | | |
|---|---|---|
| **Potential off-target sequence** | | **Locus** |
| 1 | *CACCTA-AGCCCAATTGGTA* | chr12:+111474134 |
| 2 | *CTTCTATA-CCCAATTGGTA* | chr20:+11338400 |
| 3 | *CTTCTAT-GCCTAATTGGTA* | chr20:+23807904 |
| 4 | *TTACTAG-GCCCAATTGGTA* | chr20:-39488186 |

### wildtype

**ZNF804A $^{+/+}$_4** **ZNF804A $^{+/+}$_61**

#### sgRNA1

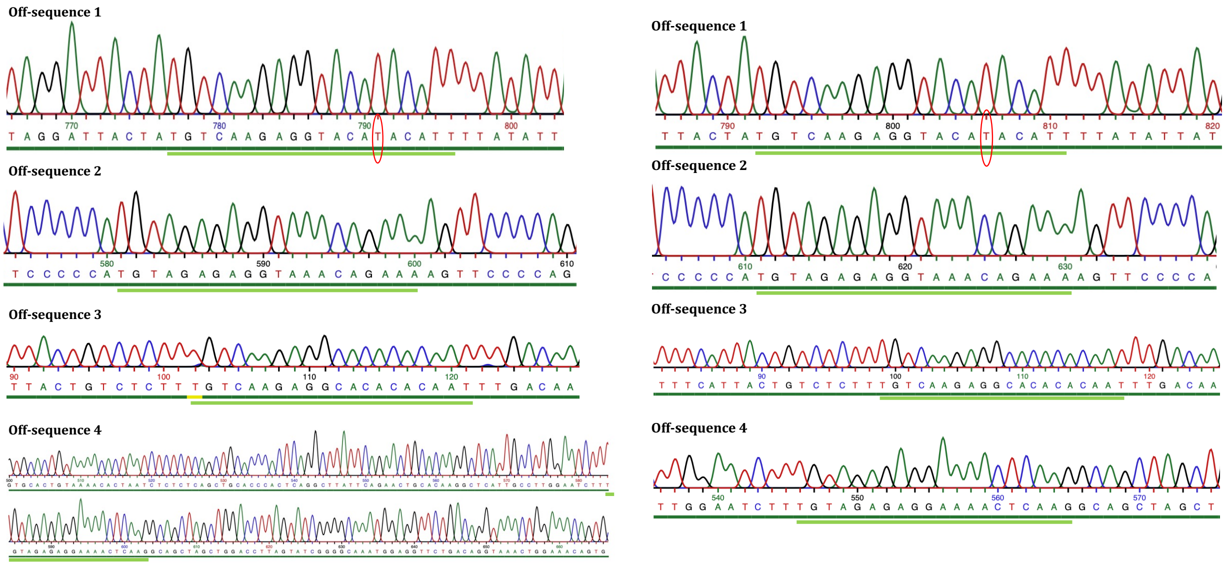

#### sgRNA2

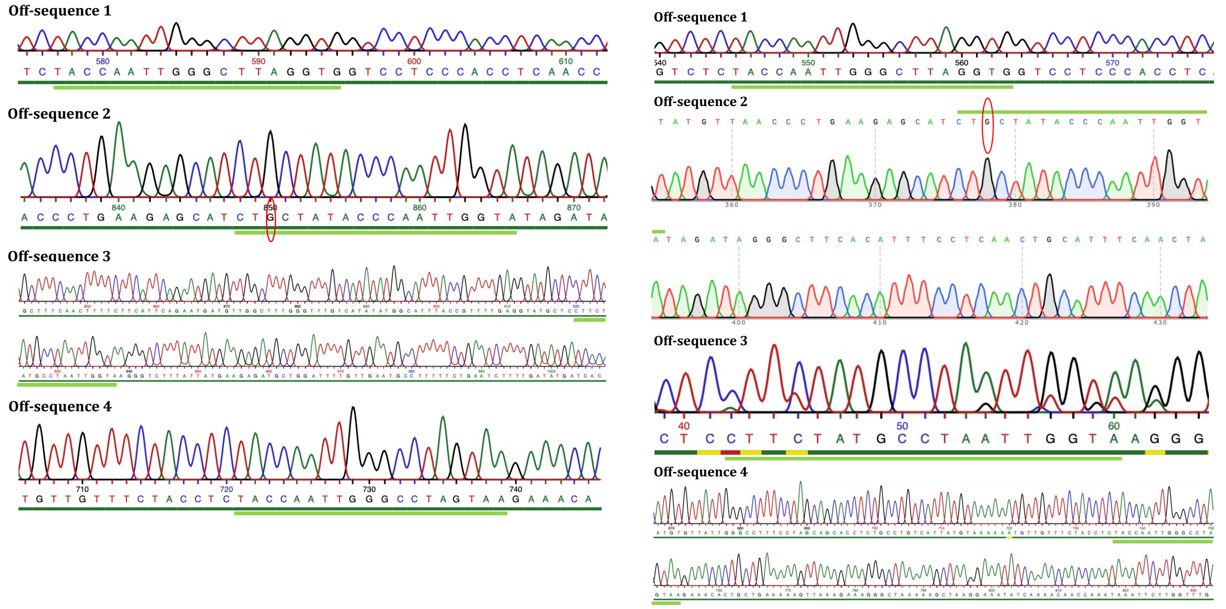

### heterozygous

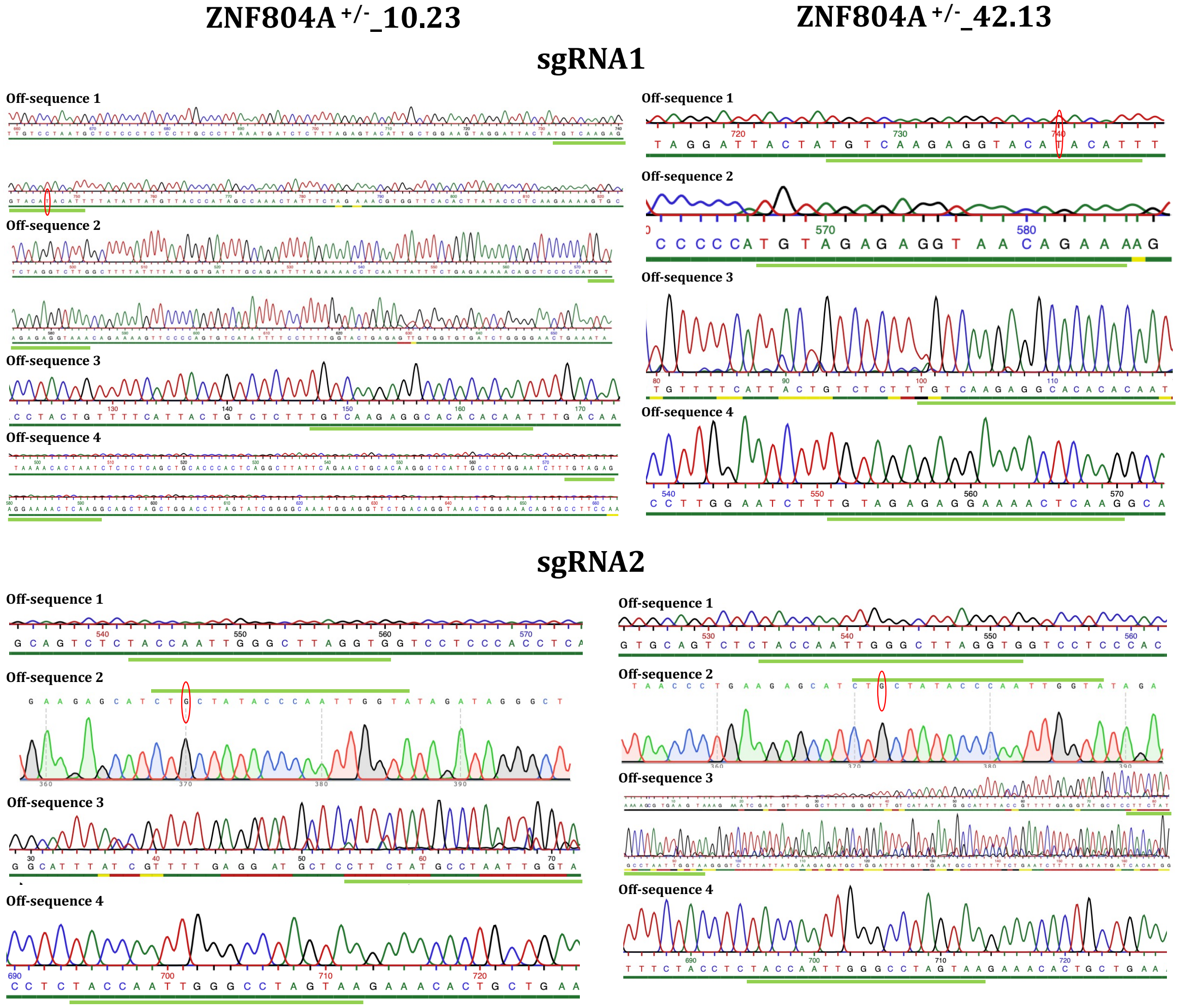

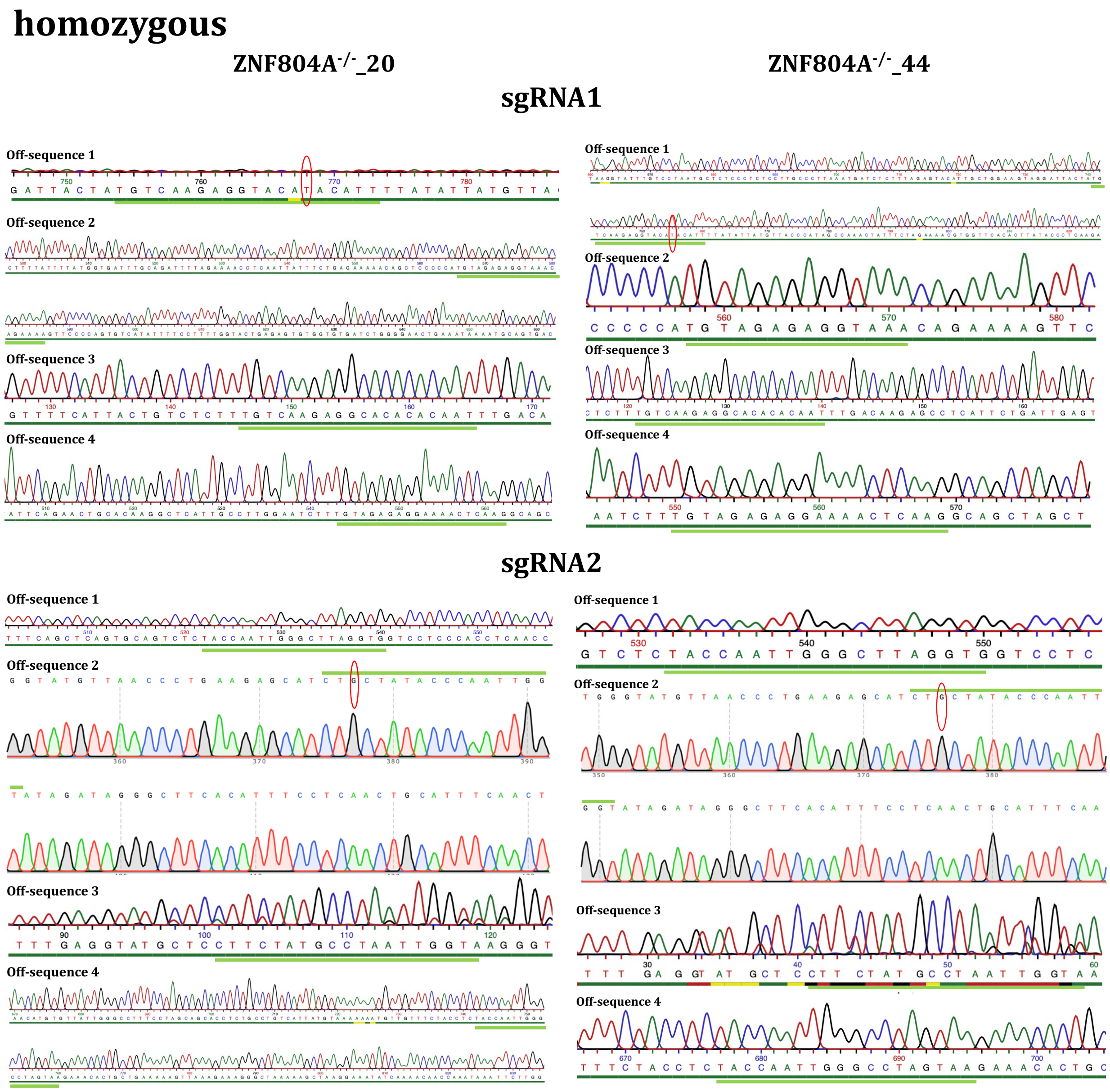

**Fig. S7 Potential off-target effect screening of gene edited clones.** Top-four predicted off-target sites of each single guide RNA (sgRNA) (table) were tested in clones using sanger sequencing. Sanger sequences show clear sequences except for two substitutions in off-target sequence 1 of sgRNA1 (C → T) and off-target sequence 2 of sgRNA2 (T → G). Off-target effects were isogenic. Substitutions are indicated by red circle and predicted off-target sites are indicated by green line.

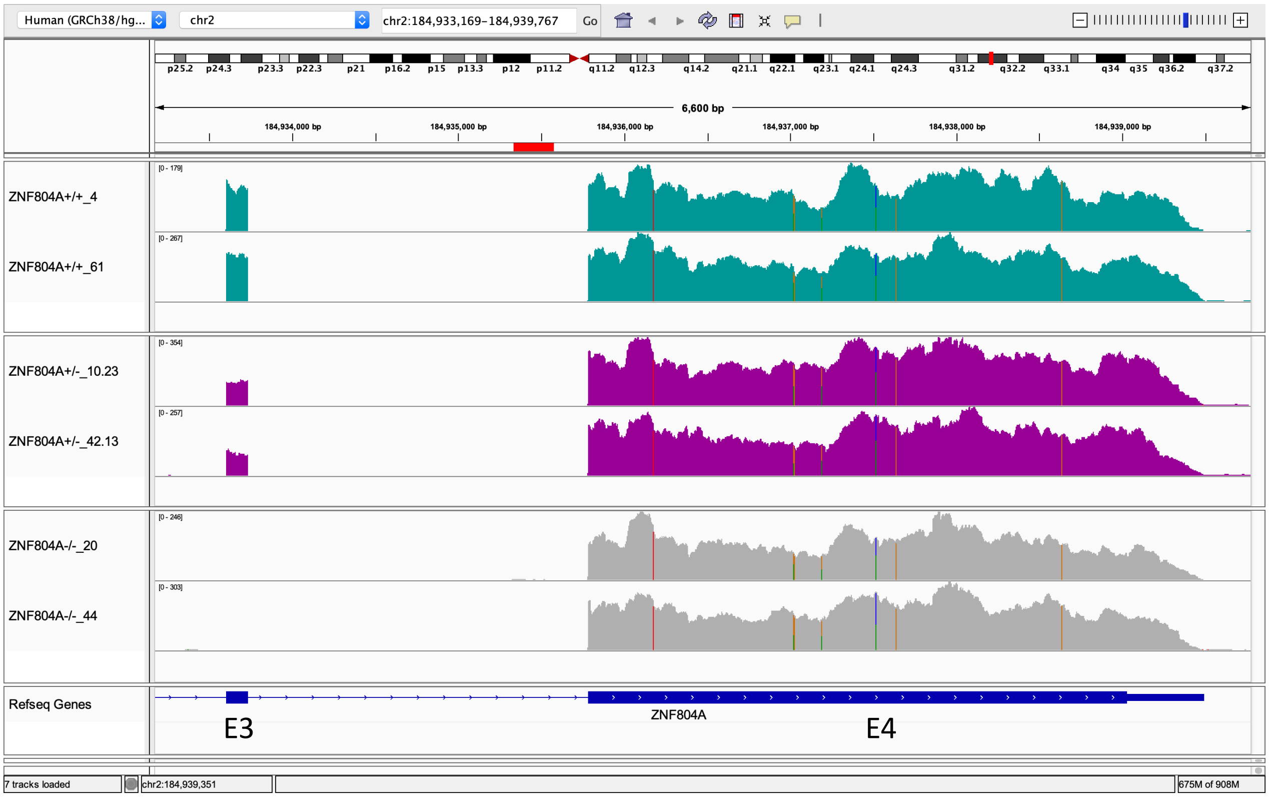

**Fig. S8 Integrative genome viewer (IGV) tracks of each ZNF804A mutant clone at 6000 base pair resolution.** Representative alignments of each clone; wildtype alignments (ZNF804A$^{+/+}$_4 and _61) are shown as turquois, heterozygous alignments (ZNF804A$^{+/-}$_10.23 and _42.13) as magenta and homozygous alignments (ZNF804A$^{-/-}$_20 and _44) as grey polygons. Reference genome sequence of ZNF804A is plotted on the bottom in blue. Exon 3 deletion was confirmed in mutation clones, whereas exon 4 seems to be transcribed.

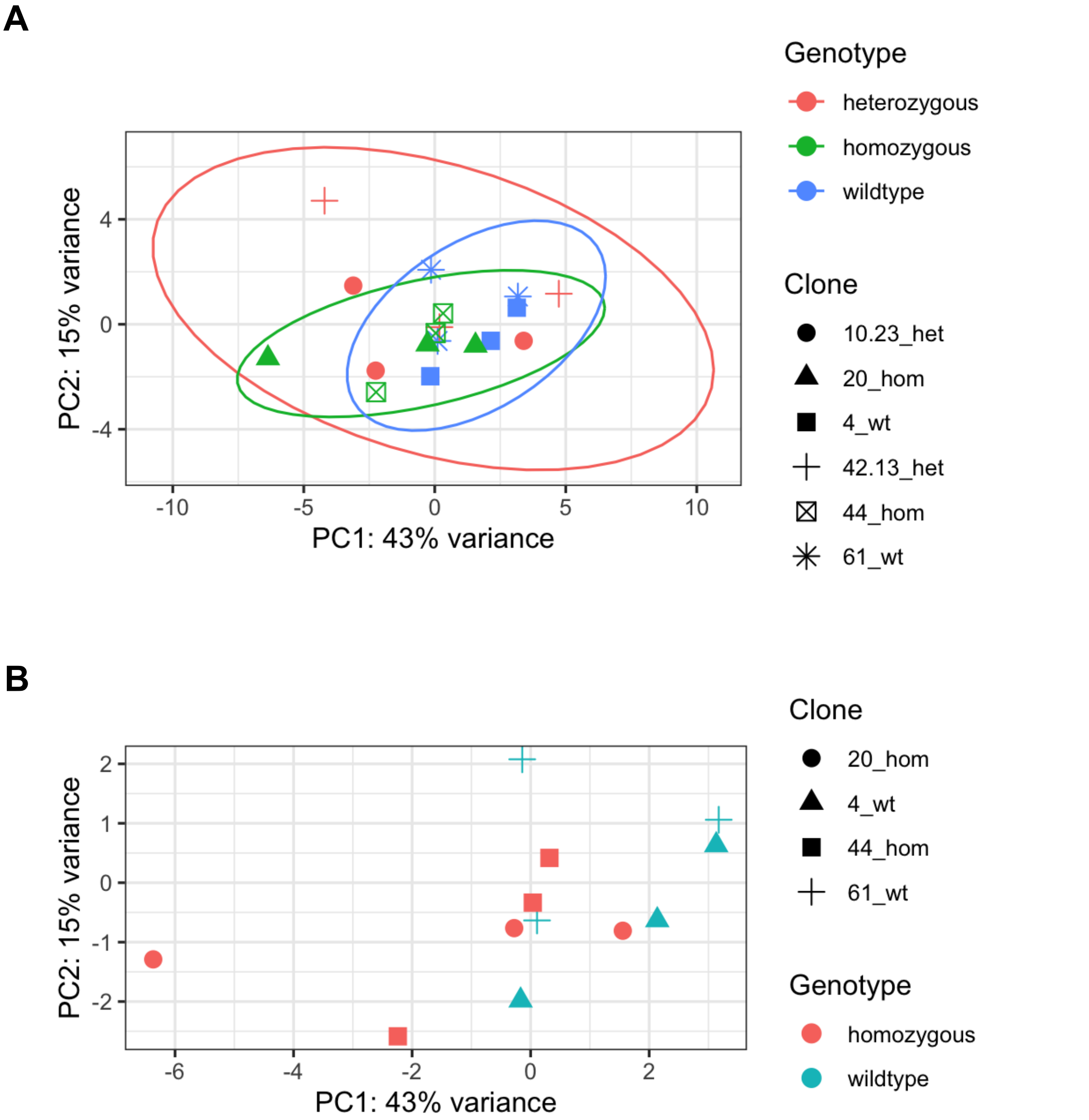

**Fig. S9 Principal component analyses (PCA) of transcriptomic data.** PCA plot of all (A) or ZNF804A$^{+/+}$ and ZNF804A$^{-/-}$ samples only (B). PC1 (43%) is plotted on x-axis, PC2 (15%) is plotted on y-axis. Color-coding according to genotype, shape-coding according to shapes. Samples cluster slightly according to ZNF804A$^{+/+}$ and ZNF804A$^{-/-}$ genotypes.

**A**

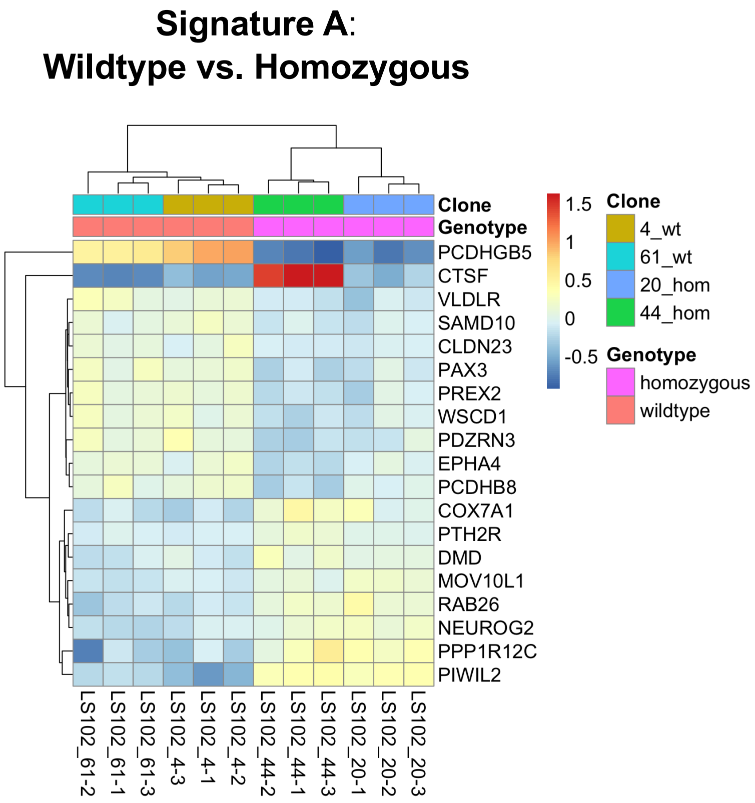

**B**

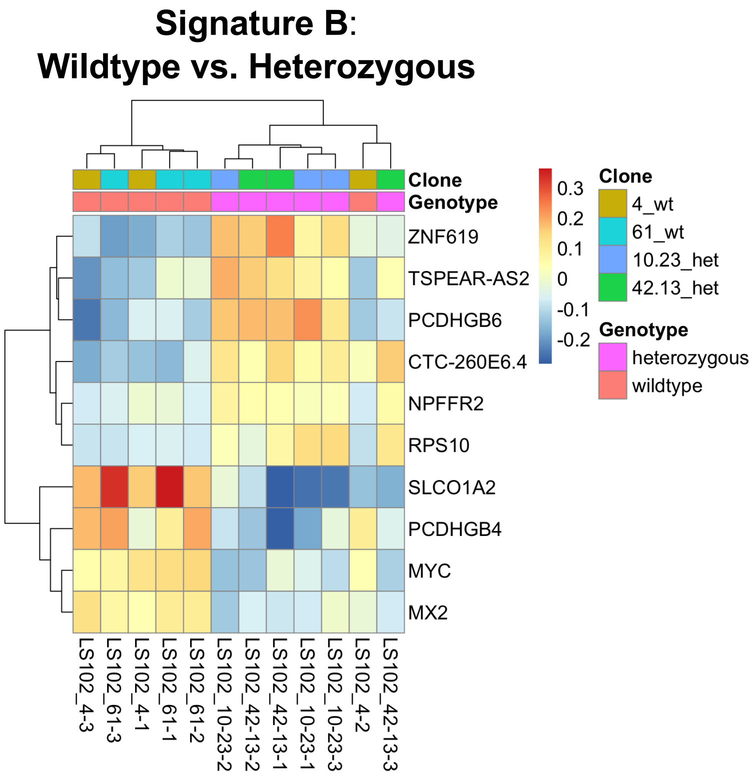

**Fig. S10 Supervised heatmaps of top protein coding DEGs of Signature A and B.**

(A) Heatmap of top 20 (10 up-, 10 down-regulated) protein coding DEGs of Signature A. Dendrogram clustering occurs according to genotype. (B) Heatmap of all DEGs of Signature B. Dendrogram clustering mainly occurs according to genotype.

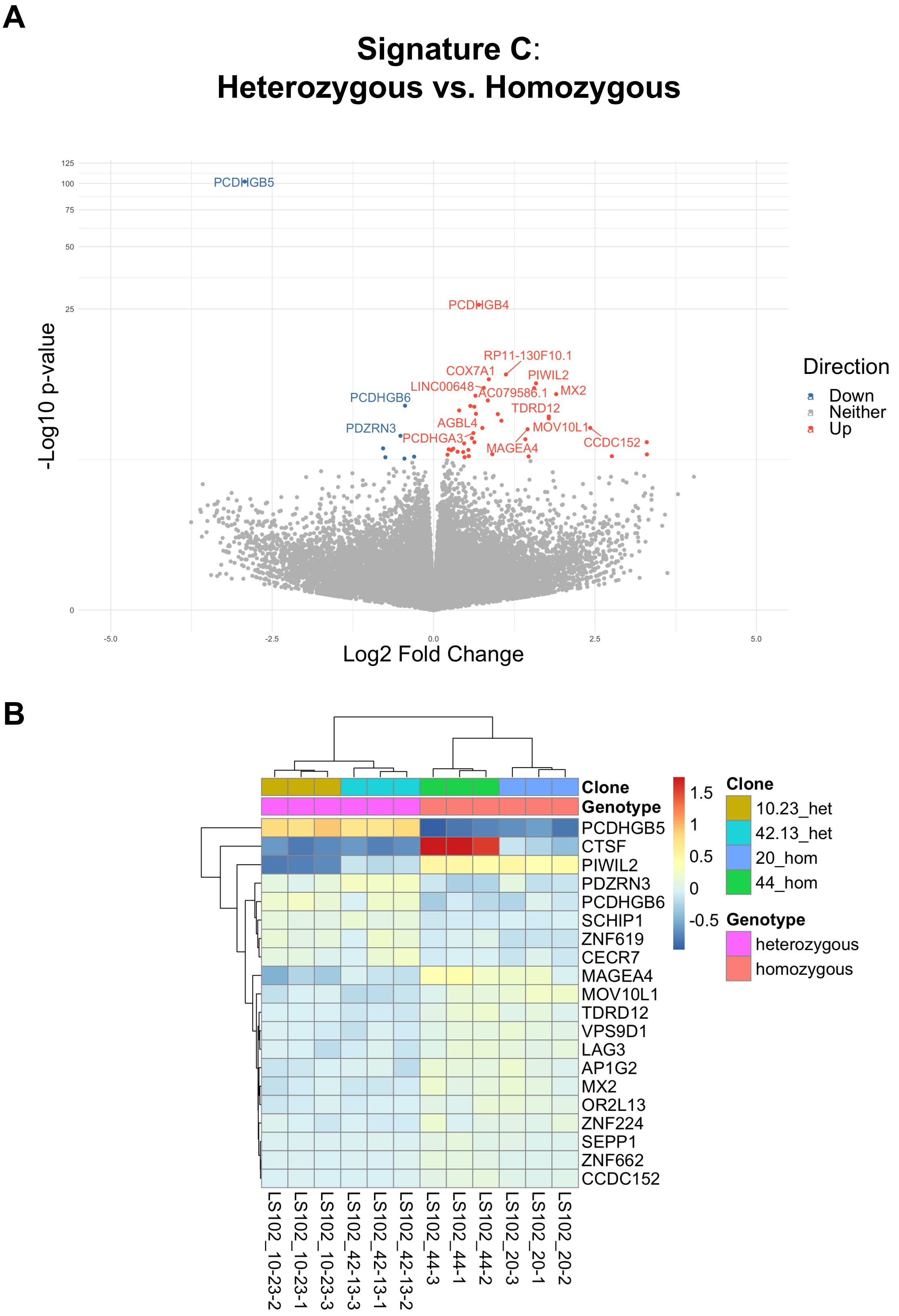

**Fig. S11 Transcriptomic effects of Signature C**. (A) Volcano plot of differentially expressed genes (DEGs; x-axis= log2 fold change, y-axis = -log10 p-value). Significantly up-regulated DEGs (FDR < 0.05) depicted in red, significantly down-regulated DEGs in blue. (B) Supervised heatmap of top 20 (10 up-, 10 down-regulated) protein-coding DEGs. Dendrogram clustering occurs according to clone and genotype.

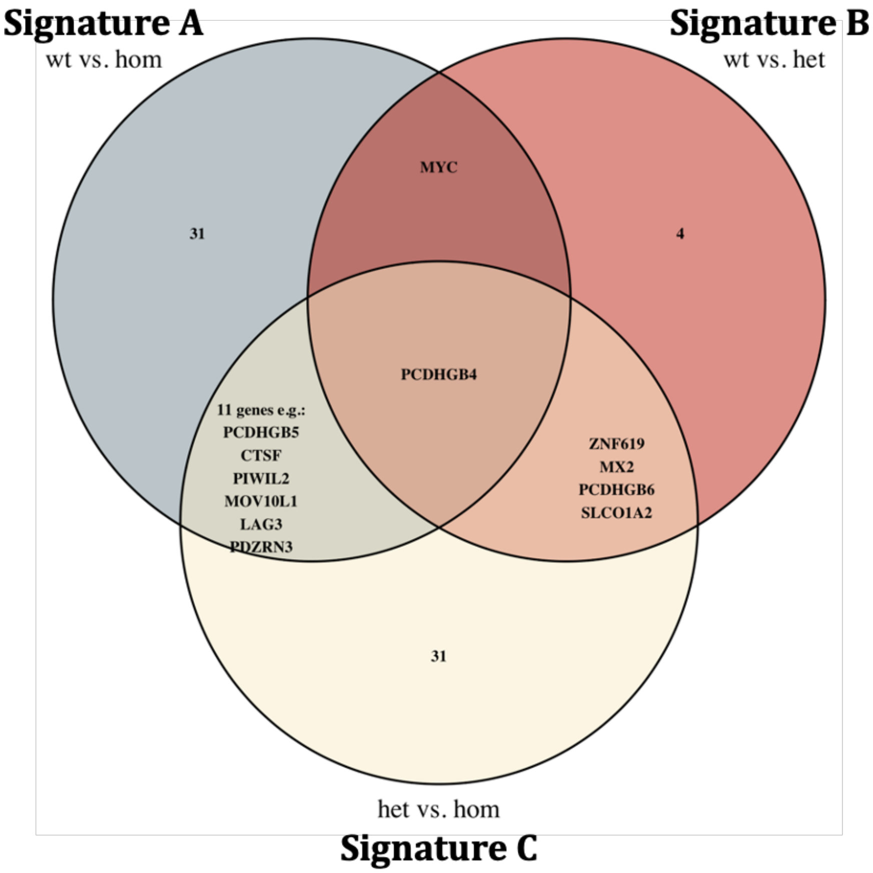

**Fig. S12 Overlapping differentially expressed genes of each Signature plotted in Venn diagram.**

A

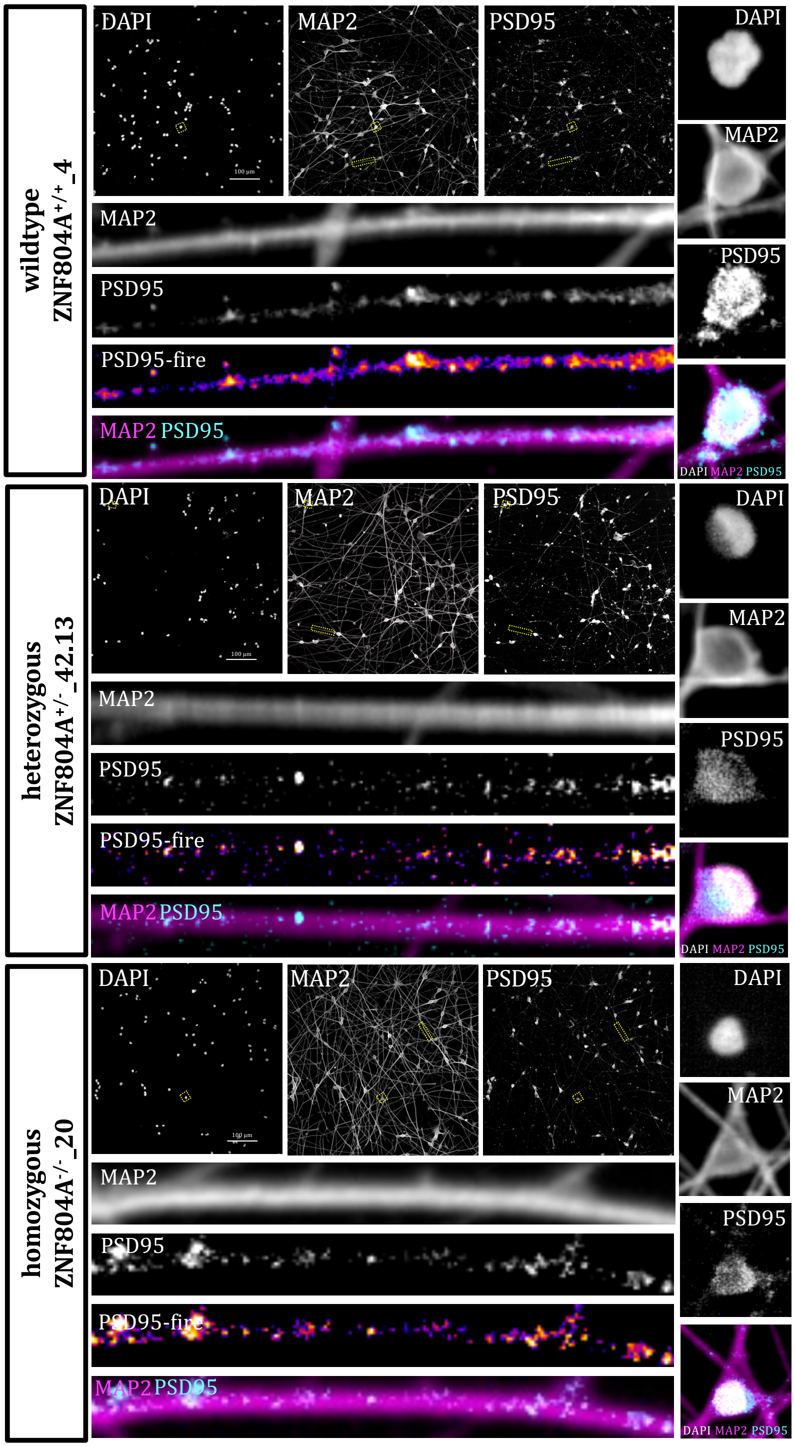
wildtype
ZNF804A+/+_4
DAPI
MAP2
PSD95
100 µm
MAP2
PSD95
PSD95-fire
MAP2 PSD95
DAPI
MAP2
PSD95
DAPI MAP2 PSD95
heterozygous
ZNF804A+/-_42.13
DAPI
MAP2
PSD95
100 µm
MAP2
PSD95
PSD95-fire
MAP2 PSD95
DAPI
MAP2
PSD95
DAPI MAP2 PSD95
homozygous
ZNF804A-/-_20
DAPI
MAP2
PSD95
100 µm
MAP2
PSD95
PSD95-fire
MAP2 PSD95
DAPI
MAP2
PSD95
DAPI MAP2 PSD95

**B**

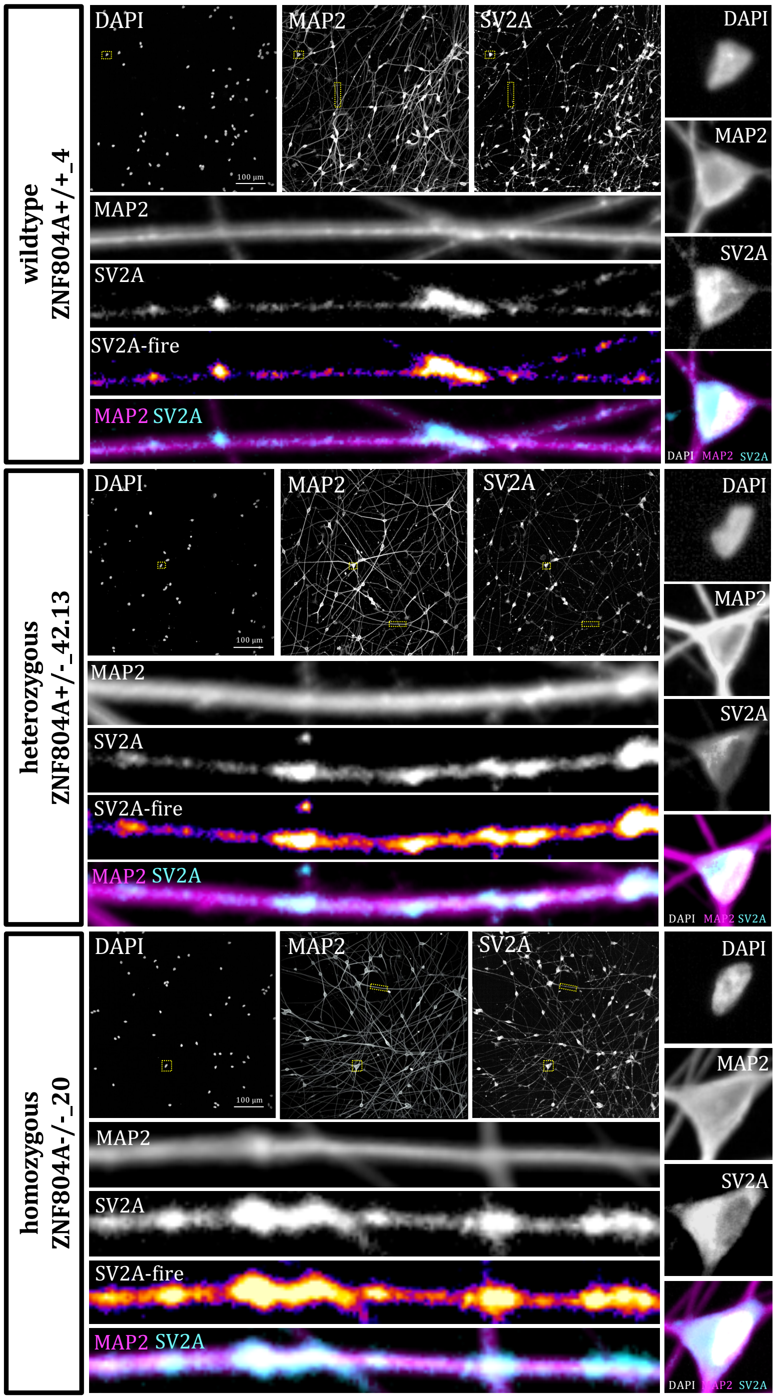
wildtype
ZNF804A+/+_4
DAPI
MAP2
SV2A
100 µm
MAP2
SV2A
SV2A-fire
MAP2 SV2A
DAPI
MAP2
SV2A
DAPI MAP2 SV2A
heterozygous
ZNF804A+/-_42.13
DAPI
MAP2
SV2A
100 µm
MAP2
SV2A
SV2A-fire
MAP2 SV2A
DAPI
MAP2
SV2A
DAPI MAP2 SV2A
homozygous
ZNF804A-/-_20
DAPI
MAP2
SV2A
100 µm
MAP2
SV2A
SV2A-fire
MAP2 SV2A
DAPI
MAP2
SV2A

C

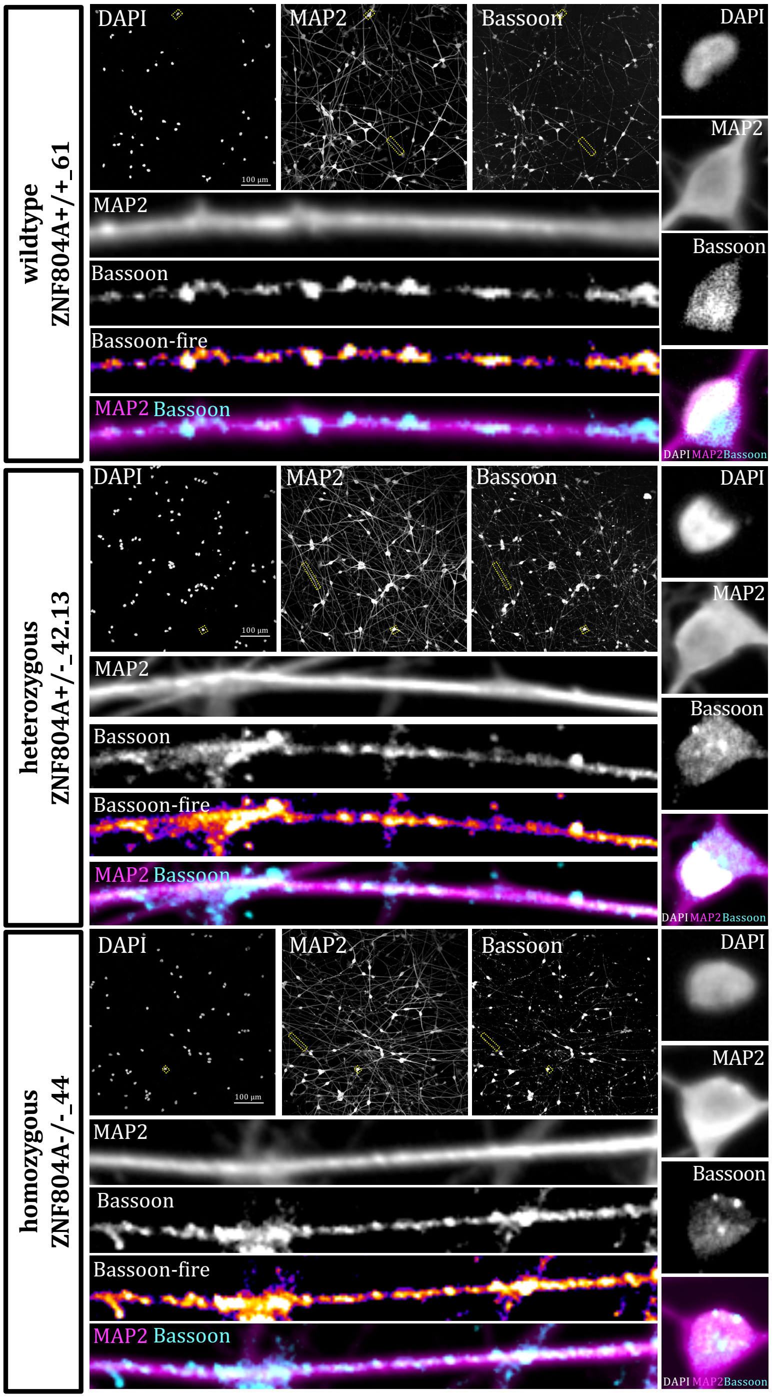
wildtype
ZNF804A+/+_61
DAPI
MAP2
Bassoon
MAP2
Bassoon
Bassoon-fire
MAP2 Bassoon
DAPI
MAP2
Bassoon
DAPI MAP2 Bassoon
heterozygous
ZNF804A+/-_42.13
DAPI
MAP2
Bassoon
MAP2
Bassoon
Bassoon-fire
MAP2 Bassoon
DAPI
MAP2
Bassoon
DAPI MAP2 Bassoon
homozygous
ZNF804A-/-_44
DAPI
MAP2
Bassoon
MAP2
Bassoon
Bassoon-fire
MAP2 Bassoon
DAPI
MAP2
Bassoon
DAPI MAP2 Bassoon

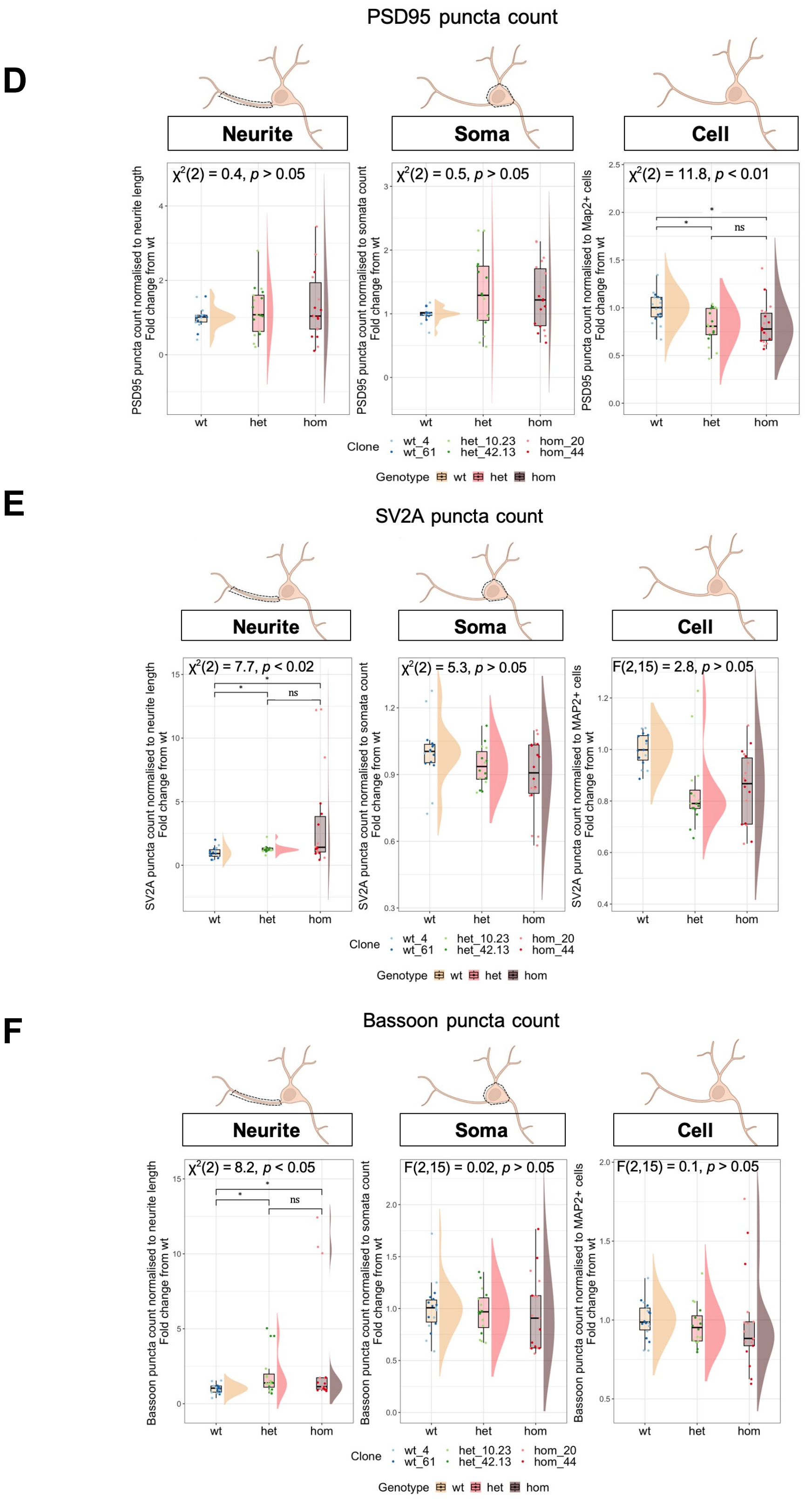

D
PSD95 puncta count
Neurite
Soma
Cell
χ2(2) = 0.4, p > 0.05
χ2(2) = 0.5, p > 0.05
χ2(2) = 11.8, p < 0.01
PSD95 puncta count normalised to neurite length Fold change from wt
PSD95 puncta count normalised to somata count Fold change from wt
PSD95 puncta count normalised to Map2+ cells Fold change from wt
ns
wt
het
hom
Clone
wt_4
wt_61
het_10.23
het_42.13
hom_20
hom_44
Genotype
wt
het
hom
E
SV2A puncta count
Neurite
Soma
Cell
χ2(2) = 7.7, p < 0.02
χ2(2) = 5.3, p > 0.05
F(2,15) = 2.8, p > 0.05
SV2A puncta count normalised to neurite length Fold change from wt
SV2A puncta count normalised to somata count Fold change from wt
SV2A puncta count normalised to MAP2+ cells Fold change from wt
ns
wt
het
hom
Clone
wt_4
wt_61
het_10.23
het_42.13
hom_20
hom_44
Genotype
wt
het
hom
F
Bassoon puncta count
Neurite
Soma
Cell
χ2(2) = 8.2, p < 0.05
F(2,15) = 0.02, p > 0.05
F(2,15) = 0.1, p > 0.05
Bassoon puncta count normalised to neurite length Fold change from wt
Bassoon puncta count normalised to somata count Fold change from wt
Bassoon puncta count normalised to MAP2+ cells Fold change from wt
ns
wt
het
hom
Clone
wt_4
wt_61
het_10.23
het_42.13
hom_20
hom_44
Genotype
wt
het
hom

**Fig. S13 Synaptic proteins are increased in and juxtaposed to MAP2$^{+}$ neurites.** Representative confocal images (A - C) of ZNF804A$^{+/+}$, ZNF804A$^{+/-}$, and ZNF804A$^{-/-}$ developing glutamatergic neurons (Day 7) from high-content confocal microscopy. Cells were immunostained for pan-neuronal marker MAP2 (magenta) to identify neurite, somata and whole cell region morphology. An example of post-synaptic density protein 95 (PSD95; A), pre-synaptic synaptic vesicle glycoprotein 2A (SV2A; B) and Bassoon (C) puncta are shown in neurite and somata regions in cyan and fire LUT. (D-F) Quantification of synaptic puncta counts in neurites, somata and whole cell regions of MAP2$^{+}$ neurons plotted in raincloud plots. Boxplots (left), corresponding density plots (right) and overlaying dotplots of technical replicates for each clones give clear visualisation of data variability. Statistical analysis on averaged technical replicates normalised to neurite length, somata or cell counts and as ratio to the respective wildtype counts acquired from images, which were imaged in the same plate, showed (D) a significant decrease of PSD95 puncta in whole cell regions (Kruskal-Wallis: $\chi^2(2)=11.8$, $p = 0.003$, $n = 18$) in ZNF804A$^{+/-}$ and ZNF804A$^{-/-}$ (Dunn's post-hoc tests with holm corrections: $p.adj < 0.05$) neurons, and (E) significant increases of SV2A (Kruskal-Wallis: $\chi^2(2) = 7.7$, $p = 0.02$, $n = 18$) and (F) Bassoon (Kruskal-Wallis: $\chi^2(2) = 8.2$, $p = 0.02$, $n = 18$) puncta lining neurite regions of ZNF804A$^{+/-}$ and ZNF804A$^{-/-}$ neurons (Dunn's post-hoc tests with holm corrections: $p.adj. < 0.05$).
*$p.adj. < 0.05$

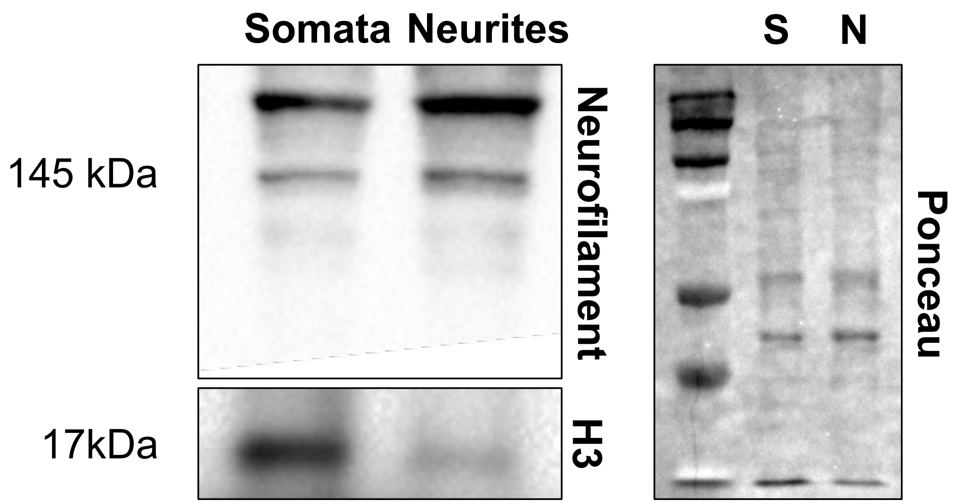

**Fig. S14 Validation of compartment separation assay.** Representative Western blot showing enrichment of Neurofilament (pan-axonal ~145-160 kDa) in neurite and histone 3 (H3 ~17 kDa) in soma sections. Loading control is shown on the right by ponceau staining.

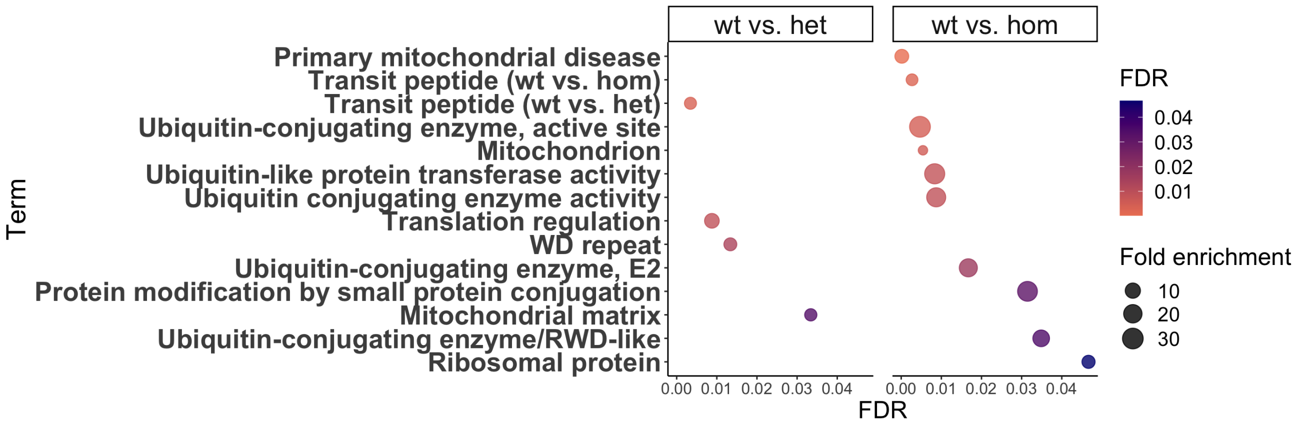

**Fig. S15** GO analysis of nominally differentially expressed proteins in ZNF804A+/- (left) and ZNF804A-/- (right) neurites. X-axis displays FDR, y-axis lists terms with significant enrichment (FDR < 0.05). Icon size represents fold enrichment.

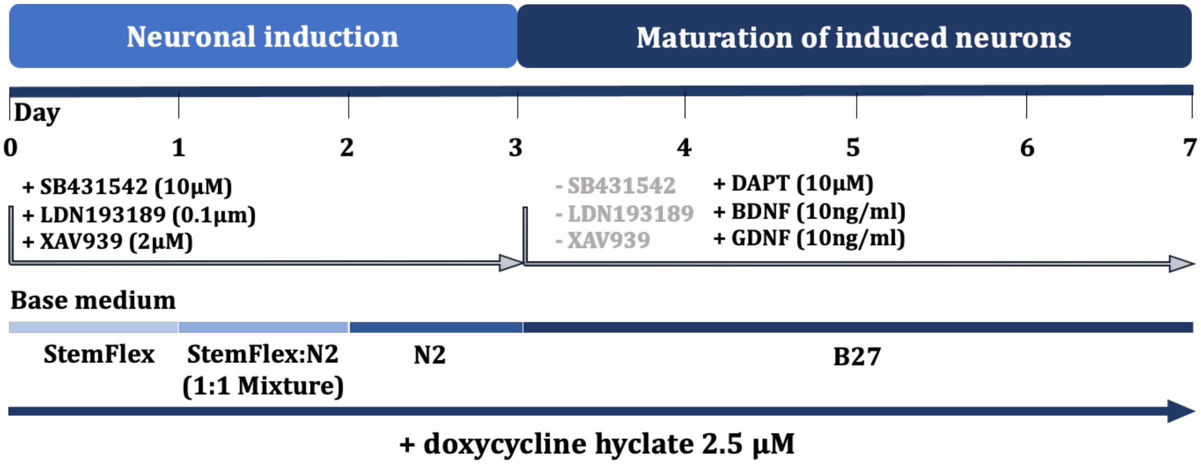

**Fig. S16 Experimental timeline of differentiation protocol using a combination of *NGN2* forward programming and SMAD and Wnt inhibition.** Cells were exposed to neuronal induction medium for 3 days with 3 different types of base media (D1 – D3 neuralisation medium) supplemented with doxycycline hyclate to activate Tet-ON-controlled *NGN2* trasngene expression and SMAD and Wnt inhibition for patterned differentiation into glutamatergic forebrain neurons. On D4 neuralisation medium was changed to B27 based maturation medium supplemented with DAPT and neurotrophic factors (BDNF and GDNF) until D7, when cells were processed for downstream analysis.

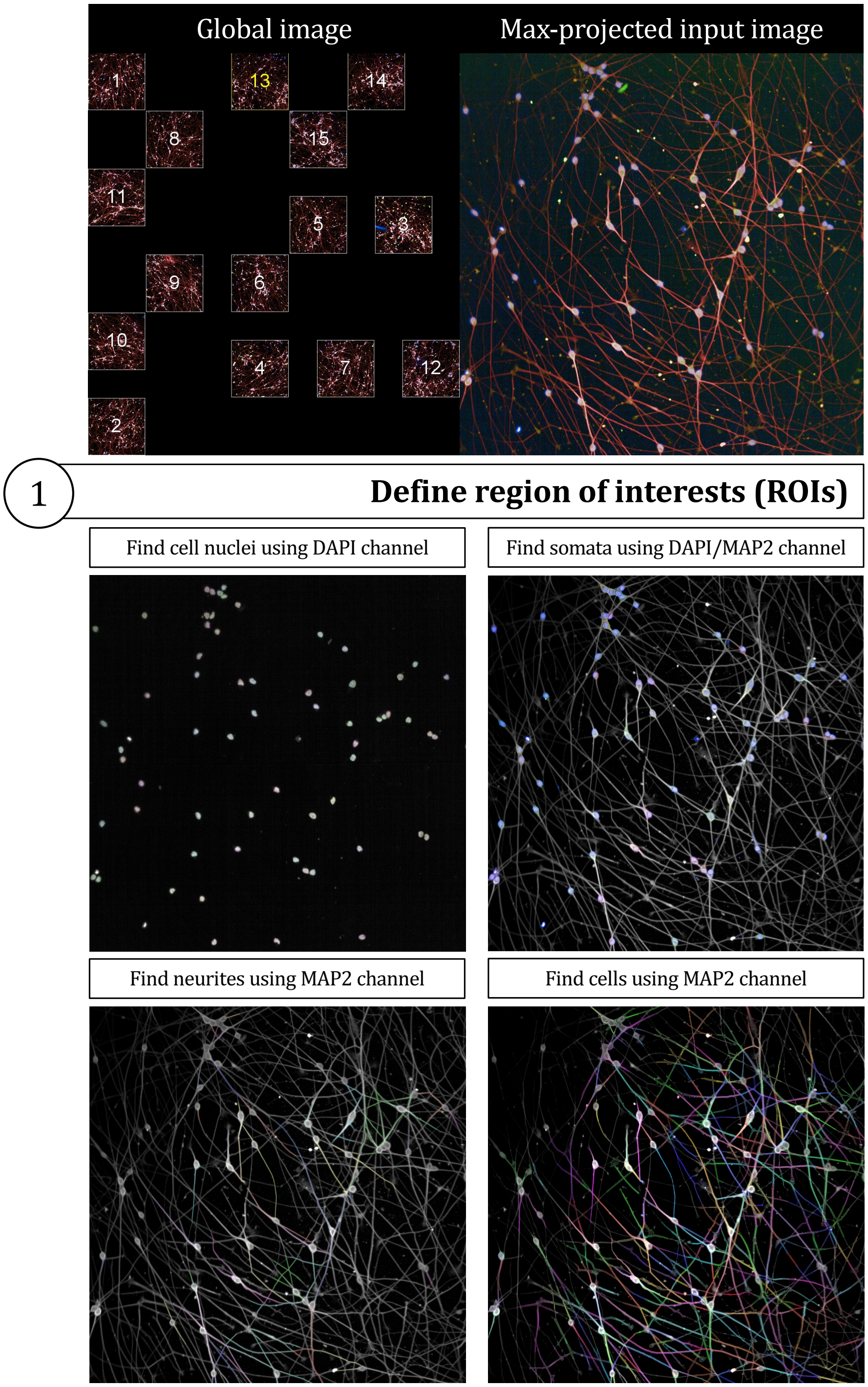
Global image
Max-projected input image
1
13
14
8
15
11
5
3
9
6
10
4
7
12
2
1
Define region of interests (ROIs)
Find cell nuclei using DAPI channel
Find somata using DAPI/MAP2 channel
Find neurites using MAP2 channel
Find cells using MAP2 channel

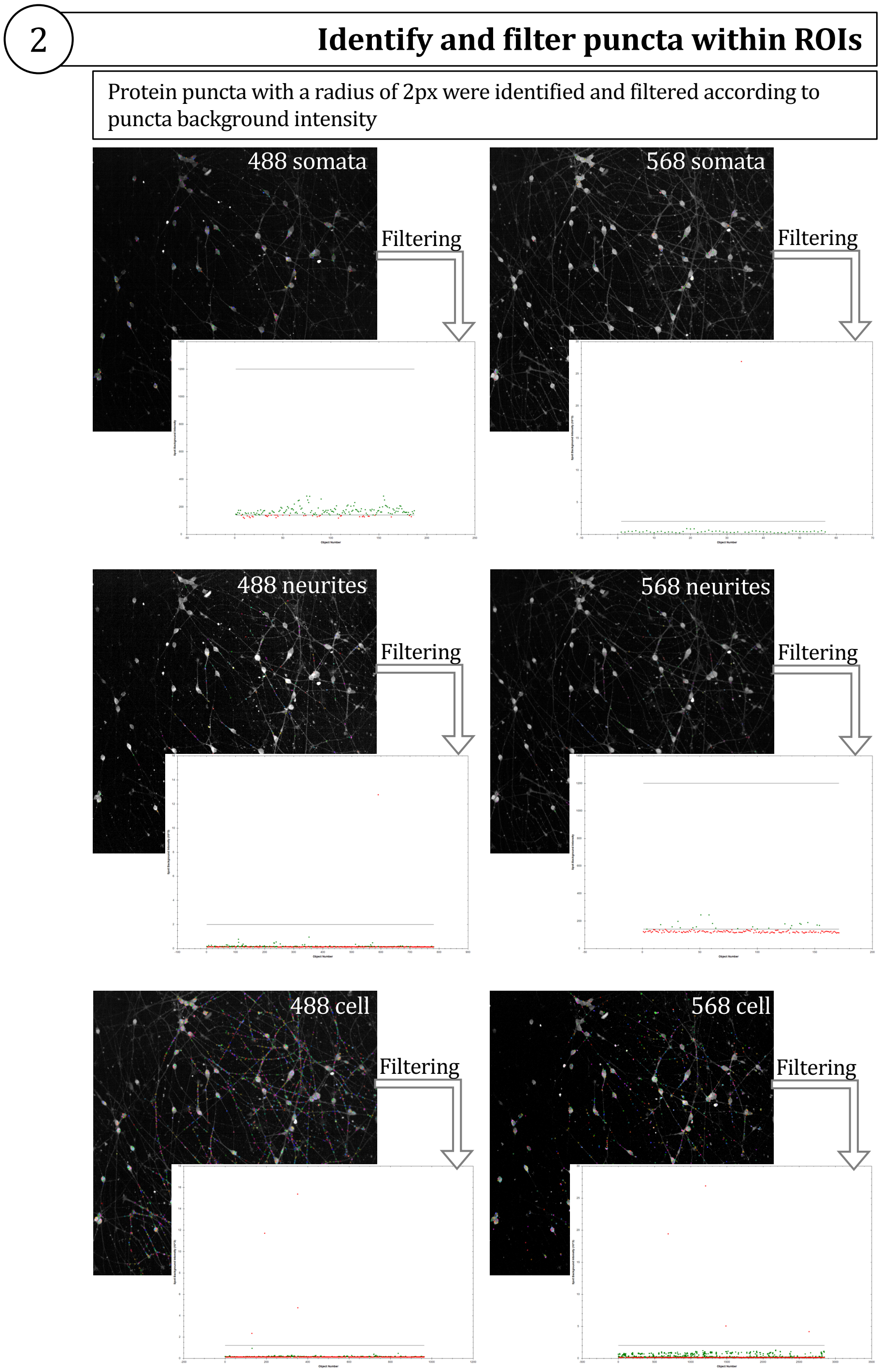

**Fig. S17 Exemplary diagram of automated analysis pipeline.** Automated analysis pipeline was instructed to (i) define regions of interests (ROIs) and (ii) detect and filter puncta within ROIs. DAPI and MAP2 channels were basis of somata identification, MAP2 channels helped to outline cells and identify neurites. Puncta detection was based on radii not exceeding 5 pixels (px) (~1.5μM) within predefined ROIs. Filtering of detected puncta was based on "Spot Background Intensity". Further analyses were conducted only on puncta that met the filtering criteria.

| Primers for ZNF804A expression analysis throughout cellular cortical development | | |
|---|---|---|
| | ***ZNF804A full-length (E2E3)*** | **Amplicon Size** |
| **Forward** | GACAAGCAGTACTATAAGCACCA | 154 |
| **Reverse** | GTGCCTTTTCCTGTTTTCTTTCA | |

| | | |
|---|---|---|
| | ***E3E4*** | **Amplicon Size** |
| **Forward** | CAGGAAAAGGCACTCCAACG | 75 |
| **Reverse** | GCCACTTCCAGGAGCACATA | |

| Genotyping primers flanking excision site in ZNF804A locus | | |
|---|---|---|
| | ***Primer sequence I*** | **Amplicon size** |
| **Forward** | TTCACAAGCCAAAATGCGAG | 1898 (wildtype) |
| **Reverse** | TCAATTCACCCAGCATTCACC | 81 (mutant) |
| | ***Primer sequence II*** | **Amplicon size** |
| **Forward** | GGATTTATAAGGCACGAATCCTTC | 2249 (wildtype) |
| **Reverse** | TGCTCTTAACCAGTTCTATCTGAC | 478 (mutant) |

| Primers for potential off-targets | | | | |
|---|---|---|---|---|
| | | ***sgRNA1*** | | |
| **Ranking** | **Chromosome** | **Potential off-target sequence** | **Primer sequence** | **Amplicon size** |
| 1 | chr1:-44387600 | AATGTGTGTACCTCTTGACA | **F:** CAGGAGTGGGCACATACGC | 1587 |
| | | | **R:** AAACTGTGCGAGTGTGTGCTA | |
| 2 | chr7:+122403326 | TTTCTGTTTACCTCTCTACA | **F:** ATGTTCACCATCAAGCAATTAGG | 1399 |
| | | | **R:** CCCCCAACTCAGCTATAAAGT | |
| 3 | chr5:+110971250 | ATTGTGTGTGCCTCTTGACA | **F:** TGGCTTCCAACACTCTGCTT | 606 |
| | | | **R:** ATCCCCTTGGTTTGCTGAGG | |
| 4 | chr3:-117476849 | CTTGAGTTTTCCTCTCTACA | **F:** ACACATGAGCCAGACAGCAA | 1723 |
| | | | **R:** AGTGTGATCGTTGAAGCCAGT | |
| | | ***sgRNA2*** | | |
| **Ranking** | **Locus** | **Potential off-target sequence** | **Primer sequence** | **Amplicon size** |
| 1 | chr12:+111474134 | CACCTA-AGCCCAATTGGTA | **F:** GCCAGTCATGTTTCATCCCTAC | 1121 |
| | | | **R:** TAAGGCGCTAAAGGGACATATTATC | |
| 2 | chr20:+11338400 | CTTCTATA-CCCAATTGGTA | **F:** AACACAAATGCCTTAAACTGGAT | 534 |
| | | | **R:** CAGAGAAATCTGCAATCAGCATCA | |
| 3 | chr20:+23807904 | CTTCTAT-GCCTAATTGGTA | **F:** TCGTAGTGGGTATCCTTGCC | 532 |
| | | | **R:** GAACTGCTGCCAATCTTACTGA | |
| 4 | chr20:-39488186 | TTACTAG-GCCCAATTGGTA | **F:** ACTATGGCACTCCTGTTGTTAG | 1674 |
| | | | **R:** TGCCCTCCCAAATACATCAATG | |

**Table S1. Primers used in this study.** Forward and reverse primer sequences including amplicon sizes ordered by experiments, for which they were used in this study.

| Three-way ANOVA | | | | | |
|---|---|---|---|---|---|
| | **Df** | **Sum Sq** | **Mean Sq** | **F value** | **Pr(>F)** |
| **Isoform** | 1 | 3.083 | 3.0831 | 15.1689 | 0.0001707 |
| **Timepoint** | 6 | 135.685 | 22.6141 | 111.2606 | < 0.0000000000000002 |
| **Line** | 2 | 11.249 | 5.6247 | 27.6731 | 1.981E-10 |
| **Isoform:Timepoint** | 6 | 0.57 | 0.095 | 0.4672 | 0.8313185 |
| **Residuals** | 108 | 21.951 | 0.2033 | | |

| TukeyHSD post-hoc corrections | | | | | |
|---|---|---|---|---|---|
| *Isoform* | | | | | |
| | **Isoform.diff** | **Isoform.lwr** | **Isoform.upr** | **Isoform.p.adj** | **Significance** |
| **E3E4-E2E3** | -0.315366281 | -0.475868181 | -0.154864382 | 0.000170685 | * |
| *Line* | | | | | |
| | **Line.diff** | **Line.lwr** | **Line.upr** | **Line.p.adj** | **Significance** |
| **127-014** | 0.727992093 | 0.491290145 | 0.964694041 | 1.47478E-10 | * |
| **M3-014** | 0.48602143 | 0.249319482 | 0.722723378 | 1.09677E-05 | ns |
| **M3-127** | -0.241970663 | -0.475768183 | -0.008173143 | 0.040738696 | * |
| *Timepoint* | | | | | |
| | **Timepoint.diff** | **Timepoint.lwr** | **Timepoint.upr** | **Timepoint.p.adj** | **Significance** |
| **D10-D0** | 1.118591507 | 0.667055284 | 1.570127729 | 5.26516E-10 | * |
| **D12-D0** | 1.606261533 | 1.15472531 | 2.057797755 | 6.06182E-14 | * |
| **D14-D0** | 2.2318497 | 1.780313477 | 2.683385922 | 1.15463E-14 | * |
| **D16-D0** | 2.618286469 | 2.166750246 | 3.069822691 | 1.15463E-14 | * |
| **D18-D0** | 3.023978176 | 2.572441953 | 3.475514398 | 1.15463E-14 | * |
| **D20-D0** | 3.134436137 | 2.669003252 | 3.599869022 | 1.15463E-14 | * |
| **D12-D10** | 0.487670026 | 0.036133803 | 0.939206248 | 0.025420684 | * |
| **D14-D10** | 1.113258193 | 0.661721971 | 1.564794415 | 6.28686E-10 | * |
| **D16-D10** | 1.499694962 | 1.048158739 | 1.951231184 | 7.41629E-14 | * |
| **D18-D10** | 1.905386669 | 1.453850446 | 2.356922891 | 1.38778E-14 | * |
| **D20-D10** | 2.01584463 | 1.550411746 | 2.481277515 | 1.26565E-14 | * |
| **D14-D12** | 0.625588167 | 0.174051945 | 1.07712439 | 0.001209944 | * |
| **D16-D12** | 1.012024936 | 0.560488714 | 1.463561159 | 1.72762E-08 | * |
| **D18-D12** | 1.417716643 | 0.966180421 | 1.869252865 | 9.69225E-14 | * |
| **D20-D12** | 1.528174605 | 1.06274172 | 1.993607489 | 7.74936E-14 | * |
| **D16-D14** | 0.386436769 | -0.065099454 | 0.837972991 | 0.14524709 | ns |
| **D18-D14** | 0.792128476 | 0.340592253 | 1.243664698 | 1.43132E-05 | * |
| **D20-D14** | 0.902586437 | 0.437153553 | 1.368019322 | 1.22523E-06 | * |
| **D18-D16** | 0.405691707 | -0.045844516 | 0.857227929 | 0.108278359 | ns |
| **D20-D16** | 0.516149668 | 0.050716784 | 0.981582553 | 0.019634721 | * |
| **D20-D18** | 0.110457962 | -0.354974923 | 0.575890846 | 0.991547485 | ns |

**Table S2. Results of statistical analysis for Fig. 1B - C.** Summary results table (top) of three-way ANOVA for isoform, timepoint, line and isoform:timepoint interaction effects on *ZNF804A* expression in neuroprogenitor cells (NPCs). Results table of Tukey's test for post-hoc analysis (bottom) with difference in means (*.diff), confidence levels (*.lwr, *.upr) and the adjusted p-values for all possible pairs for each tested variable (isoform, line and timepoint).

| Gene ID | Gene | Regional specificity |
|---|---|---|
| DCX | Doublecortin | pan-neuronal |
| MAP2 | Microtubule-associated protein 2 | pan-neuronal |
| MAPT | microtubule-associated protein tau | pan-neuronal |
| RBFOX3 | RNA Binding Fox-1 Homolog 3 | pan-neuronal |
| EMX1 | Empty Spiracles Homeobox 1 | dorsal forebrain |
| DMRTA1 | Doublesex- And Mab-3-Related Transcription Factor A1 | dorsal forebrain |
| DMRT3 | Doublesex And Mab-3 Related Transcription Factor 3 | dorsal forebrain |
| GSX2 | GS Homeobox 2 | ventral forebrain |
| NKX2-1 | NK2 Homeobox 1 | ventral forebrain |
| DLX2 | Distal-Less Homeobox 2 | ventral forebrain |
| LMX1A | LIM Homeobox transcription factor 1, alpha | midbrain |
| EN1 | Engrailed Homeobox 1 | midbrain |
| EN2 | Engrailed Homeobox 2 | midbrain |
| FGF8 | Fibroblast growth factor 8 | hindbrain |
| GBX2 | Gastrulation Brain Homeobox 2 | hindbrain |
| HOXB1 | Homeobox B1 | hindbrain |
| TLE1 | TLE Family Member 1, Transcriptional Corepressor | upper layer cortical |
| CUX1 | Homeobox protein cut-like 1 | upper layer cortical |
| CUX2 | Homeobox protein cut-like 2 | upper layer cortical |
| LIMCH1 | LIM And Calponin Homology Domains 1 | upper layer cortical |
| RELN | Reelin | upper layer cortical |
| POU3F3 | POU Class 3 Homeobox 3 | upper layer cortical |
| RORB | RAR Related Orphan Receptor B | deep layer cortical |
| BCL11B | B cell leukemia 11b | deep layer cortical |
| TBR1 | T-Box Brain Transcription Factor 1 | deep layer cortical |
| FEZF2 | FEZ Family Zinc Finger 2 | deep layer cortical |

**Table S3. List of genes expressed in all RNAseq samples.**

| DEGs from Signature A: wildtype vs. homozygous | | | | | | |
|---|---|---|---|---|---|---|
| | **baseMean** | **log2FoldChange** | **lfcSE** | **stat** | **pvalue** | **padj** |
| **PCDHGB5** | 841.2775654 | -2.829672243 | 0.13583736 | -20.831326 | 2.25E-96 | 4.72E-92 |
| **RP11-402J6.1** | 35886.59494 | 0.726698803 | 0.10255285 | 7.08609046 | 1.38E-12 | 1.45E-08 |
| **RAB26** | 2627.018296 | 0.547801304 | 0.0957138 | 5.72332642 | 1.04E-08 | 7.30E-05 |
| **COX7A1** | 472.8798777 | 0.672670959 | 0.12414269 | 5.41853059 | 6.01E-08 | 0.00031507 |
| **RP11-469N6.1** | 152.1321969 | 0.967728977 | 0.18157421 | 5.32966087 | 9.84E-08 | 0.00041273 |
| **PAX3** | 237.8921331 | -0.725937085 | 0.13978741 | -5.1931506 | 2.07E-07 | 0.00054206 |
| **PCDHGB4** | 4377.382025 | 0.338447343 | 0.06508236 | 5.2002932 | 1.99E-07 | 0.00054206 |
| **FRRS1L** | 1069.620489 | 0.3598253 | 0.06919512 | 5.20015462 | 1.99E-07 | 0.00054206 |
| **PDZRN3** | 659.4517806 | -0.554365338 | 0.10906087 | -5.083082 | 3.71E-07 | 0.00086539 |
| **CTSF** | 175.5922075 | 3.809005416 | 0.76239938 | 4.99607624 | 5.85E-07 | 0.00112257 |
| **PIWIL2** | 501.6636391 | 1.177053452 | 0.23667733 | 4.97324123 | 6.58E-07 | 0.00112257 |
| **AC079586.1** | 100.6695458 | 1.177845289 | 0.23685725 | 4.97280658 | 6.60E-07 | 0.00112257 |
| **CLDN23** | 30.80333051 | -1.604078536 | 0.32323821 | -4.9625276 | 6.96E-07 | 0.00112257 |
| **SHANK2** | 1232.673387 | 0.403353443 | 0.08252658 | 4.88755788 | 1.02E-06 | 0.00152945 |
| **CADM2** | 2524.144943 | 0.316552043 | 0.06523448 | 4.85252652 | 1.22E-06 | 0.00159786 |
| **RP11-78I14.1** | 359.6856588 | -0.75382643 | 0.15532864 | -4.8531064 | 1.22E-06 | 0.00159786 |
| **VLDLR** | 1217.511073 | -0.453277445 | 0.09514099 | -4.7642707 | 1.90E-06 | 0.00233834 |
| **WSCD1** | 2223.867446 | -0.385006329 | 0.08230469 | -4.6778175 | 2.90E-06 | 0.00337834 |
| **CTD-2587H24.1** | 3300.085988 | -0.802622988 | 0.17303483 | -4.6385055 | 3.51E-06 | 0.0038738 |
| **LINC00654** | 213.9538259 | 0.532420439 | 0.11901686 | 4.47348746 | 7.70E-06 | 0.00806979 |
| **PPP1R12C** | 5800.403771 | 0.864355832 | 0.1946821 | 4.43983218 | 9.00E-06 | 0.00899133 |
| **RIMS4** | 6640.279721 | 0.347495714 | 0.07864186 | 4.41871168 | 9.93E-06 | 0.00946559 |
| **GALNT18** | 970.5535812 | -0.322866913 | 0.07377896 | -4.3761379 | 1.21E-05 | 0.0105595 |
| **CCDC106** | 689.6239271 | -0.300460548 | 0.06865982 | -4.3760751 | 1.21E-05 | 0.0105595 |
| **MOV10L1** | 40.68937834 | 1.425698243 | 0.33032901 | 4.31599469 | 1.59E-05 | 0.01332925 |
| **MAPK3** | 2882.577186 | -0.175613624 | 0.04110414 | -4.272407 | 1.93E-05 | 0.01361529 |
| **SMPD3** | 9317.349852 | 0.422276406 | 0.09861932 | 4.28188329 | 1.85E-05 | 0.01361529 |
| **KCNK15** | 670.3204444 | 0.397860711 | 0.09296336 | 4.27975822 | 1.87E-05 | 0.01361529 |
| **PTH2R** | 18.47137094 | 1.980243475 | 0.46042835 | 4.30087215 | 1.70E-05 | 0.01361529 |
| **NEUROG2** | 209769.0483 | 0.42567169 | 0.09966975 | 4.27082111 | 1.95E-05 | 0.01361529 |
| **EPS8L1** | 1908.28559 | -0.279644186 | 0.06720715 | -4.1609291 | 3.17E-05 | 0.02144357 |
| **EPB41** | 14730.17842 | -0.167877927 | 0.04091169 | -4.1034219 | 4.07E-05 | 0.02668053 |
| **DMD** | 106.0841267 | 0.781249043 | 0.19079936 | 4.09461037 | 4.23E-05 | 0.02687588 |
| **ST6GALNAC5** | 1381.964056 | -0.26984769 | 0.06629471 | -4.0704257 | 4.69E-05 | 0.02894724 |
| **MYC** | 2308.080018 | -0.185719066 | 0.04593313 | -4.0432485 | 5.27E-05 | 0.03071129 |
| **CTB-12A17.3** | 8625.86689 | -0.17809998 | 0.04398422 | -4.0491786 | 5.14E-05 | 0.03071129 |
| **EPHA4** | 1068.251706 | -0.346056156 | 0.0859088 | -4.0281804 | 5.62E-05 | 0.0310236 |
| **PCDHB8** | 906.3981575 | -0.412753684 | 0.1024016 | -4.0307345 | 5.56E-05 | 0.0310236 |
| **PREX2** | 159.0307004 | -0.796913069 | 0.19986937 | -3.9871696 | 6.69E-05 | 0.0359586 |
| **SAMD10** | 139.3981786 | -0.629365312 | 0.15811711 | -3.9803744 | 6.88E-05 | 0.03607713 |
| **TTC39A** | 171.766541 | 0.389703864 | 0.09825241 | 3.96635442 | 7.30E-05 | 0.0373321 |
| **HEG1** | 4276.678802 | -0.228912446 | 0.05788455 | -3.9546379 | 7.67E-05 | 0.03827605 |
| **LAG3** | 467.2268409 | 0.27737664 | 0.07089509 | 3.91249411 | 9.13E-05 | 0.04354174 |
| **RP4-665J23.1** | 114.3694552 | -0.464728412 | 0.11871719 | -3.9145839 | 9.06E-05 | 0.04354174 |

| DEGs from Signature B: wildtype vs. heterozygous | | | | | | |
|---|---|---|---|---|---|---|
| | **baseMean** | **log2FoldChange** | **lfcSE** | **stat** | **pvalue** | **padj** |
| **MYC** | 2308.080018 | -0.272323359 | 0.04610451 | -5.9066532 | 3.49E-09 | 7.89E-05 |
| **PCDHGB4** | 4377.382025 | -0.362832783 | 0.06542408 | -5.54586 | 2.93E-08 | 3.31E-04 |
| **ZNF619** | 196.728203 | 0.616716877 | 0.11474311 | 5.37476182 | 7.67E-08 | 4.34E-04 |
| **TSPEAR-AS2** | 70.8751824 | 0.987735381 | 0.18268266 | 5.40683707 | 6.41E-08 | 0.00043351 |
| **CTC-260E6.4** | 86.93013908 | 0.821049194 | 0.16282258 | 5.04260025 | 4.59E-07 | 0.002077 |
| **MX2** | 33.39716097 | -1.489467348 | 0.30838081 | -4.8299613 | 1.37E-06 | 0.0051467 |
| **RPS10** | 74.02250928 | 0.674262634 | 0.14068737 | 4.79263074 | 1.65E-06 | 0.00531756 |
| **PCDHGB6** | 5902.346688 | 0.364077533 | 0.07647315 | 4.76085461 | 1.93E-06 | 0.00544902 |
| **NPFFR2** | 9.598915161 | 2.865225204 | 0.6123969 | 4.67870625 | 2.89E-06 | 0.00725351 |
| **SLCO1A2** | 627.4470502 | -0.647005421 | 0.14066151 | -4.5997333 | 4.23E-06 | 0.00956603 |

| DEGs from Signature C: heterozygous vs. homozygous | | | | | | |
|---|---|---|---|---|---|---|
| | **baseMean** | **log2FoldChange** | **lfcSE** | **stat** | **pvalue** | **padj** |
| **PCDHGB5** | 841.2775654 | -2.925058143 | 0.13580697 | -21.538351 | 6.81E-103 | 1.99E-98 |
| **PCDHGB4** | 4377.382025 | 0.701280126 | 0.06529313 | 10.7404887 | 6.57E-27 | 9.58E-23 |
| **CTSF** | 175.5922075 | 6.180194833 | 0.7907352 | 7.81575783 | 5.46E-15 | 5.31E-11 |
| **RP11-130F10.1** | 148.9185444 | 1.120119689 | 0.15850885 | 7.06660652 | 1.59E-12 | 1.1579E-08 |
| **COX7A1** | 472.8798777 | 0.854959399 | 0.12474079 | 6.85388786 | 7.19E-12 | 4.1931E-08 |
| **PIWIL2** | 501.6636391 | 1.586437718 | 0.2374336 | 6.68160583 | 2.36E-11 | 1.1491E-07 |
| **LINC00648** | 414.0604868 | 0.777677786 | 0.11982076 | 6.49034245 | 8.56E-11 | 3.3358E-07 |
| **AC079586.1** | 100.6695458 | 1.556282943 | 0.24015235 | 6.48039865 | 9.15E-11 | 3.3358E-07 |
| **MX2** | 33.39716097 | 1.898941947 | 0.30451868 | 6.23588008 | 4.49E-10 | 1.4562E-06 |
| **RP11-71L14.3** | 491.678133 | 0.648461697 | 0.10507031 | 6.17169294 | 6.76E-10 | 1.9709E-06 |
| **SLCO1A2** | 627.4470502 | 0.840162399 | 0.14042764 | 5.98288499 | 2.19E-09 | 5.8137E-06 |
| **PCDHGB6** | 5902.346688 | -0.442198199 | 0.07649683 | -5.7806085 | 7.44E-09 | 1.7751E-05 |
| **SMPD3** | 9317.349852 | 0.569252933 | 0.09865126 | 5.77035624 | 7.91E-09 | 1.7751E-05 |
| **PCDHGA8** | 217.6532312 | 0.630537337 | 0.10986198 | 5.73935887 | 9.50E-09 | 1.9803E-05 |
| **PCDHA11** | 659.3835204 | 0.395789684 | 0.07067869 | 5.59984497 | 2.15E-08 | 4.1724E-05 |
| **LINC00654** | 213.9538259 | 0.656487273 | 0.11990435 | 5.47509155 | 4.37E-08 | 7.9728E-05 |
| **RP11-469N6.1** | 152.1321969 | 0.994373451 | 0.181983 | 5.46410072 | 4.65E-08 | 7.9838E-05 |
| **TDRD12** | 24.29487706 | 1.784421756 | 0.33182941 | 5.37752737 | 7.55E-08 | 0.00012239 |
| **MOV10L1** | 40.68937834 | 1.785665439 | 0.33630411 | 5.30967481 | 1.10E-07 | 0.00016862 |
| **OR2L13** | 42.8337835 | 1.050470373 | 0.20100826 | 5.22600606 | 1.73E-07 | 0.00025265 |
| **CCDC152** | 10.03946324 | 2.426607875 | 0.48816661 | 4.97086001 | 6.67E-07 | 0.00088853 |
| **AGBL4** | 148.1557937 | 0.755431557 | 0.15200323 | 4.96983875 | 6.70E-07 | 0.00088853 |
| **MAGEA4** | 129.7154867 | 1.455656741 | 0.29549921 | 4.92609347 | 8.39E-07 | 0.00106402 |
| **PCDHGA3** | 104.1982301 | 0.616453875 | 0.12858523 | 4.79412681 | 1.63E-06 | 0.00198594 |
| **PDZRN3** | 659.4517806 | -0.513362333 | 0.10920454 | -4.7009248 | 2.59E-06 | 0.00302205 |
| **TRIM58** | 347.1960593 | 0.5934447 | 0.12833761 | 4.62409049 | 3.76E-06 | 0.00422148 |
| **RP11-366F6.2** | 22.02862623 | 1.421430283 | 0.30984434 | 4.58756259 | 4.48E-06 | 0.00484526 |
| **ZNF454** | 139.261743 | 0.632460718 | 0.14081963 | 4.4912825 | 7.08E-06 | 0.0071228 |
| **ZNF662** | 6.262576192 | 3.301732519 | 0.73514861 | 4.49124503 | 7.08E-06 | 0.0071228 |
| **PCDHGA1** | 252.9631299 | 0.472922897 | 0.10621237 | 4.45261602 | 8.48E-06 | 0.00824891 |
| **TSPEAR-AS1** | 57.26790318 | -0.781688069 | 0.18212418 | -4.2920609 | 1.77E-05 | 0.01665838 |
| **LAG3** | 467.2268409 | 0.30456985 | 0.07112233 | 4.28233765 | 1.85E-05 | 0.0168596 |
| **VPS9D1** | 1181.966402 | 0.234613536 | 0.05506773 | 4.26045379 | 2.04E-05 | 0.01803468 |
| **DOC2B** | 472.3308145 | 0.538843329 | 0.12702414 | 4.24205469 | 2.21E-05 | 0.01900322 |
| **AP1G2** | 2089.123612 | 0.274625257 | 0.06485953 | 4.23415423 | 2.29E-05 | 0.01912128 |
| **ZNF568** | 703.9677385 | 0.372280887 | 0.08891861 | 4.18676022 | 2.83E-05 | 0.02292956 |
| **SPINK5** | 363.8298164 | 0.458577367 | 0.10980455 | 4.17630584 | 2.96E-05 | 0.02335979 |
| **SLCO4A1-AS1** | 33.8243245 | 0.909765526 | 0.22145828 | 4.10806726 | 3.99E-05 | 0.0304652 |
| **RP11-333A23.4** | 7.777936735 | 3.303983704 | 0.8052005 | 4.10330558 | 4.07E-05 | 0.0304652 |
| **ZNF224** | 1493.140943 | 0.215727737 | 0.05269533 | 4.09386805 | 4.24E-05 | 0.03093952 |
| **PCDHGA2** | 248.0287569 | 0.545575436 | 0.13479056 | 4.04757883 | 5.18E-05 | 0.03602524 |
| **SEPP1** | 6.986410251 | 2.760719576 | 0.68215581 | 4.04705134 | 5.19E-05 | 0.03602524 |
| **CTD-3006G17.2** | 21.5522394 | 1.471578038 | 0.36432642 | 4.03917469 | 5.36E-05 | 0.03639011 |
| **SCHIP1** | 475.1726663 | -0.301213689 | 0.07470559 | -4.0320099 | 5.53E-05 | 0.03666514 |
| **LGI1** | 363.1476344 | 0.479045917 | 0.11927514 | 4.01630966 | 5.91E-05 | 0.03798391 |
| **CECR7** | 66.53543161 | -0.746737082 | 0.18606914 | -4.0132237 | 5.99E-05 | 0.03798391 |
| **ZNF619** | 196.728203 | -0.451350249 | 0.11361107 | -3.9727663 | 7.10E-05 | 0.04409487 |

**Table S4. Differentially expressed genes (DEGs) of Signatures A, B and C.**

| Category | Term | Count | % | PValue | List Total | Pop Hits | Pop Total | Fold Enrichmen | Bonferroni | Benjamini | FDR |
|---|---|---|---|---|---|---|---|---|---|---|---|
| UP_KW_CELLULAR_COMPONENT | KW-0770~Synapse | 9 | 20.00 | 1.25E-05 | 40 | 507 | 17615 | 7.82 | 2.37E-04 | 2.37E-04 | 2.37E-04 |
| UP_KW_CELLULAR_COMPONENT | KW-0472~Membrane | 29 | 64.44 | 0.001299399 | 40 | 8221 | 17615 | 1.55 | 0.024401973 | 0.012344291 | 0.012344291 |
| UP_KW_DOMAIN | KW-1133~Transmembrane helix | 24 | 53.33 | 0.00198362 | 36 | 5824 | 14495 | 1.66 | 0.033191684 | 0.020271021 | 0.019078608 |
| UP_KW_DOMAIN | KW-0812~Transmembrane | 24 | 53.33 | 0.002384826 | 36 | 5893 | 14495 | 1.64 | 0.039777703 | 0.020271021 | 0.019078608 |
| UP_KW_CELLULAR_COMPONENT | KW-0965~Cell junction | 6 | 13.33 | 0.004533527 | 40 | 496 | 17615 | 5.33 | 0.082711142 | 0.026155438 | 0.026155438 |
| UP_KW_CELLULAR_COMPONENT | KW-1003~Cell membrane | 17 | 37.78 | 0.005506408 | 40 | 3854 | 17615 | 1.94 | 0.09959522 | 0.026155438 | 0.026155438 |
| UP_KW_DOMAIN | KW-0732~Signal | 19 | 42.22 | 0.006454173 | 36 | 4354 | 14495 | 1.76 | 0.104234441 | 0.036573648 | 0.034422257 |
| UP_KW_BIOLOGICAL_PROCESS | KW-0130~Cell adhesion | 6 | 13.33 | 0.001383988 | 21 | 489 | 11223 | 6.56 | 0.044674572 | 0.045671601 | 0.045671601 |
| GOTERM_CC_DIRECT | GO:0070161~anchoring junction | 7 | 15.56 | 4.08E-04 | 43 | 481 | 20562 | 6.96 | 0.052801878 | 0.054235936 | 0.054235936 |
| UP_SEQ_FEATURE | TRANSMEM:Helical | 23 | 51.11 | 2.43E-04 | 43 | 5358 | 20559 | 2.05 | 0.05854352 | 0.06031982 | 0.06031982 |
| UP_KW_MOLECULAR_FUNCTION | KW-0217~Developmental protein | 7 | 15.56 | 0.006527799 | 22 | 991 | 11693 | 3.75 | 0.156570663 | 0.084861387 | 0.084861387 |
| UP_KW_MOLECULAR_FUNCTION | KW-9996~Developmental protein | 7 | 15.56 | 0.006527799 | 22 | 991 | 11693 | 3.75 | 0.156570663 | 0.084861387 | 0.084861387 |
| UP_KW_PTM | KW-0325~Glycoprotein | 19 | 42.22 | 0.010346776 | 34 | 4731 | 14106 | 1.67 | 0.108105503 | 0.124161306 | 0.124161306 |
| GOTERM_BP_DIRECT | GO:0007156~homophilic cell adhesion via plasma membrane adhesion molecules | 5 | 11.11 | 3.54E-04 | 39 | 172 | 19308 | 14.39 | 0.147426646 | 0.155877481 | 0.155531087 |
| GOTERM_BP_DIRECT | GO:0007155~cell adhesion | 7 | 15.56 | 6.93E-04 | 39 | 555 | 19308 | 6.24 | 0.267918185 | 0.155877481 | 0.155531087 |
| UP_KW_CELLULAR_COMPONENT | KW-0628~Postsynaptic cell membrane | 3 | 6.67 | 0.043469122 | 40 | 150 | 17615 | 8.81 | 0.570185672 | 0.165182664 | 0.165182664 |
| GOTERM_BP_DIRECT | GO:0007399~nervous system development | 6 | 13.33 | 0.001253181 | 39 | 415 | 19308 | 7.16 | 0.431233292 | 0.187977094 | 0.187559368 |
| INTERPRO | IPR013761:Sterile alpha motif/pointed domain | 4 | 8.89 | 0.002282243 | 41 | 126 | 19204 | 14.87 | 0.267095418 | 0.310385034 | 0.310385034 |
| GOTERM_CC_DIRECT | GO:0016021~integral component of membrane | 20 | 44.44 | 0.006291487 | 43 | 5415 | 20562 | 1.77 | 0.568035187 | 0.381717249 | 0.381717249 |
| GOTERM_CC_DIRECT | GO:0071546~pi-body | 2 | 4.44 | 0.014212962 | 43 | 7 | 20562 | 136.62 | 0.851011501 | 0.381717249 | 0.381717249 |
| GOTERM_CC_DIRECT | GO:0014069~postsynaptic density | 4 | 8.89 | 0.017119912 | 43 | 267 | 20562 | 7.16 | 0.899406163 | 0.381717249 | 0.381717249 |
| GOTERM_CC_DIRECT | GO:0005886~plasma membrane | 18 | 40.00 | 0.017215586 | 43 | 5071 | 20562 | 1.70 | 0.900700147 | 0.381717249 | 0.381717249 |
| GOTERM_CC_DIRECT | GO:0045202~synapse | 5 | 11.11 | 0.017220327 | 43 | 488 | 20562 | 4.90 | 0.900763832 | 0.381717249 | 0.381717249 |
| GOTERM_CC_DIRECT | GO:0005883~neurofilament | 2 | 4.44 | 0.020243688 | 43 | 10 | 20562 | 95.64 | 0.934127621 | 0.384630063 | 0.384630063 |
| GOTERM_BP_DIRECT | GO:0048710~regulation of astrocyte differentiation | 2 | 4.44 | 0.003932421 | 39 | 2 | 19308 | 495.08 | 0.830190456 | 0.442397316 | 0.441414211 |
| GOTERM_CC_DIRECT | GO:0030424~axon | 4 | 8.89 | 0.037221063 | 43 | 361 | 20562 | 5.30 | 0.993557785 | 0.549862446 | 0.549862446 |
| GOTERM_CC_DIRECT | GO:0098978~glutamatergic synapse | 4 | 8.89 | 0.040389151 | 43 | 373 | 20562 | 5.13 | 0.995844198 | 0.549862446 | 0.549862446 |
| GOTERM_CC_DIRECT | GO:0045211~postsynaptic membrane | 3 | 6.67 | 0.041343041 | 43 | 158 | 20562 | 9.08 | 0.996359094 | 0.549862446 | 0.549862446 |
| INTERPRO | IPR013164:Cadherin, N-terminal | 3 | 6.67 | 0.008099333 | 41 | 65 | 19204 | 21.62 | 0.669118273 | 0.550754655 | 0.550754655 |
| GOTERM_CC_DIRECT | GO:0043186~P granule | 2 | 4.44 | 0.045963277 | 43 | 23 | 20562 | 41.58 | 0.998085051 | 0.555737801 | 0.555737801 |
| UP_SEQ_FEATURE | DOMAIN:Cadherin 6 | 3 | 6.67 | 0.012202475 | 43 | 82 | 20559 | 17.49 | 0.952395546 | 0.563335442 | 0.563335442 |
| UP_SEQ_FEATURE | TOPO_DOM:Cytoplasmic | 15 | 33.33 | 0.015802056 | 43 | 3807 | 20559 | 1.88 | 0.980749212 | 0.563335442 | 0.563335442 |
| UP_SEQ_FEATURE | DOMAIN:SAM | 3 | 6.67 | 0.016134413 | 43 | 95 | 20559 | 15.10 | 0.982296012 | 0.563335442 | 0.563335442 |
| UP_SEQ_FEATURE | DOMAIN:Cadherin 5 | 3 | 6.67 | 0.019132082 | 43 | 104 | 20559 | 13.79 | 0.991693628 | 0.563335442 | 0.563335442 |
| UP_SEQ_FEATURE | DOMAIN:Cadherin 3 | 3 | 6.67 | 0.02161299 | 43 | 111 | 20559 | 12.92 | 0.995567506 | 0.563335442 | 0.563335442 |
| UP_SEQ_FEATURE | DOMAIN:Cadherin 4 | 3 | 6.67 | 0.02161299 | 43 | 111 | 20559 | 12.92 | 0.995567506 | 0.563335442 | 0.563335442 |
| UP_SEQ_FEATURE | CARBOHYD:N-linked (GlcNAc...) asparagine | 16 | 35.56 | 0.02237572 | 43 | 4376 | 20559 | 1.75 | 0.99634699 | 0.563335442 | 0.563335442 |
| UP_SEQ_FEATURE | DOMAIN:Cadherin 2 | 3 | 6.67 | 0.022715139 | 43 | 114 | 20559 | 12.58 | 0.996648413 | 0.563335442 | 0.563335442 |
| UP_SEQ_FEATURE | DOMAIN:Cadherin 1 | 3 | 6.67 | 0.022715139 | 43 | 114 | 20559 | 12.58 | 0.996648413 | 0.563335442 | 0.563335442 |
| INTERPRO | IPR001660:Sterile alpha motif domain | 3 | 6.67 | 0.0163872 | 41 | 94 | 19204 | 14.95 | 0.894296988 | 0.59563336 | 0.59563336 |
| INTERPRO | IPR020894:Cadherin conserved site | 3 | 6.67 | 0.023144062 | 41 | 113 | 19204 | 12.44 | 0.958604721 | 0.59563336 | 0.59563336 |
| INTERPRO | IPR002126:Cadherin | 3 | 6.67 | 0.025083401 | 41 | 118 | 19204 | 11.91 | 0.96840814 | 0.59563336 | 0.59563336 |
| INTERPRO | IPR015919:Cadherin-like | 3 | 6.67 | 0.026277942 | 41 | 121 | 19204 | 11.61 | 0.973260018 | 0.59563336 | 0.59563336 |
| GOTERM_CC_DIRECT | GO:0032991~macromolecular complex | 5 | 11.11 | 0.054468919 | 43 | 703 | 20562 | 3.40 | 0.999418054 | 0.603697188 | 0.603697188 |
| UP_KW_LIGAND | KW-0730~Sialic acid | 2 | 4.44 | 0.055636003 | 16 | 26 | 6833 | 32.85 | 0.467237515 | 0.611996038 | 0.611996038 |
| GOTERM_CC_DIRECT | GO:0005794~Golgi apparatus | 6 | 13.33 | 0.08141168 | 43 | 1141 | 20562 | 2.51 | 0.999987552 | 0.782531197 | 0.782531197 |
| GOTERM_CC_DIRECT | GO:0048786~presynaptic active zone | 2 | 4.44 | 0.082371705 | 43 | 42 | 20562 | 22.77 | 0.999989169 | 0.782531197 | 0.782531197 |
| INTERPRO | IPR001478:PDZ domain | 3 | 6.67 | 0.043279716 | 41 | 159 | 19204 | 8.84 | 0.99756355 | 0.840863054 | 0.840863054 |
| GOTERM_CC_DIRECT | GO:0005856~cytoskeleton | 4 | 8.89 | 0.09797018 | 43 | 541 | 20562 | 3.54 | 0.999998892 | 0.868668926 | 0.868668926 |
| UP_SEQ_FEATURE | DOMAIN:PDZ | 3 | 6.67 | 0.039017281 | 43 | 153 | 20559 | 9.37 | 0.999948303 | 0.879662327 | 0.879662327 |
| GOTERM_BP_DIRECT | GO:0045727~positive regulation of translation | 3 | 6.67 | 0.011878866 | 39 | 84 | 19308 | 17.68 | 0.99538061 | 1 | 1 |
| SMART | SM00454:SAM | 3 | 6.67 | 0.024825248 | 31 | 85 | 10386 | 11.82 | 0.685375427 | 1 | 1 |
| GOTERM_BP_DIRECT | GO:0065003~macromolecular complex assembly | 3 | 6.67 | 0.035976155 | 39 | 152 | 19308 | 9.77 | 0.999999931 | 1 | 1 |
| GOTERM_BP_DIRECT | GO:0043046~DNA methylation involved in gamete generation | 2 | 4.44 | 0.038653262 | 39 | 20 | 19308 | 49.51 | 0.99999998 | 1 | 1 |
| GOTERM_BP_DIRECT | GO:0034587~piRNA metabolic process | 2 | 4.44 | 0.038653262 | 39 | 20 | 19308 | 49.51 | 0.99999998 | 1 | 1 |
| SMART | SM00112:CA | 3 | 6.67 | 0.043904414 | 31 | 116 | 10386 | 8.66 | 0.873217181 | 1 | 1 |
| KEGG_PATHWAY | hsa04514:Cell adhesion molecules | 3 | 6.67 | 0.045967732 | 19 | 157 | 8164 | 8.21 | 0.999098644 | 1 | 1 |
| KEGG_PATHWAY | hsa05160:Hepatitis C | 3 | 6.67 | 0.045967732 | 19 | 157 | 8164 | 8.21 | 0.999098644 | 1 | 1 |
| BIOCARTA | h_cdmacPathway:Cadmium induces DNA synthesis and proliferation in macrophages | 2 | 4.44 | 0.061315725 | 7 | 17 | 1623 | 27.28 | 0.989494734 | 1 | 1 |
| GOTERM_BP_DIRECT | GO:0007628~adult walking behavior | 2 | 4.44 | 0.062993588 | 39 | 33 | 19308 | 30.00 | 1 | 1 | 1 |
| GOTERM_MF_DIRECT | GO:0005509~calcium ion binding | 5 | 11.11 | 0.063155519 | 39 | 751 | 18869 | 3.22 | 0.99969326 | 1 | 1 |
| GOTERM_BP_DIRECT | GO:1905606~regulation of presynapse assembly | 2 | 4.44 | 0.064840864 | 39 | 34 | 19308 | 29.12 | 1 | 1 | 1 |
| GOTERM_BP_DIRECT | GO:0051973~positive regulation of telomerase activity | 2 | 4.44 | 0.064840864 | 39 | 34 | 19308 | 29.12 | 1 | 1 | 1 |
| INTERPRO | IPR011993:Pleckstrin homology-like domain | 4 | 8.89 | 0.065008618 | 41 | 445 | 19204 | 4.21 | 0.999892889 | 1 | 1 |
| UP_SEQ_FEATURE | DOMAIN:Ig-like C2-type 2 | 3 | 6.67 | 0.066246487 | 43 | 206 | 20559 | 6.96 | 0.999999959 | 1 | 1 |
| UP_SEQ_FEATURE | DOMAIN:Ig-like C2-type 1 | 3 | 6.67 | 0.066246487 | 43 | 206 | 20559 | 6.96 | 0.999999959 | 1 | 1 |
| UP_SEQ_FEATURE | TOPO_DOM:Extracellular | 11 | 24.44 | 0.066855444 | 43 | 2929 | 20559 | 1.80 | 0.999999965 | 1 | 1 |
| KEGG_PATHWAY | hsa05205:Proteoglycans in cancer | 3 | 6.67 | 0.073780915 | 19 | 205 | 8164 | 6.29 | 0.999989027 | 1 | 1 |
| GOTERM_BP_DIRECT | GO:0048813~dendrite morphogenesis | 2 | 4.44 | 0.074024183 | 39 | 39 | 19308 | 25.39 | 1 | 1 | 1 |
| SMART | SM00228:PDZ | 3 | 6.67 | 0.074170696 | 31 | 156 | 10386 | 6.44 | 0.971131617 | 1 | 1 |
| KEGG_PATHWAY | hsa05216:Thyroid cancer | 2 | 4.44 | 0.078588478 | 19 | 37 | 8164 | 23.23 | 0.999994947 | 1 | 1 |
| BIOCARTA | h_gleevecPathway:Inhibition of Cellular Proliferation by Gleevec | 2 | 4.44 | 0.082193885 | 7 | 23 | 1623 | 20.16 | 0.997919977 | 1 | 1 |
| UP_SEQ_FEATURE | DOMAIN:PDZ 1 | 2 | 4.44 | 0.082383239 | 43 | 42 | 20559 | 22.77 | 0.999999999 | 1 | 1 |
| UP_SEQ_FEATURE | DOMAIN:PDZ 2 | 2 | 4.44 | 0.082383239 | 43 | 42 | 20559 | 22.77 | 0.999999999 | 1 | 1 |
| KEGG_PATHWAY | hsa05219:Bladder cancer | 2 | 4.44 | 0.086726009 | 19 | 41 | 8164 | 20.96 | 0.999998652 | 1 | 1 |
| UP_KW_BIOLOGICAL_PROCESS | KW-0524~Neurogenesis | 3 | 6.67 | 0.09102796 | 21 | 286 | 11223 | 5.61 | 0.957128635 | 1 | 1 |
| INTERPRO | IPR003599:Immunoglobulin subtype | 4 | 8.89 | 0.092254773 | 41 | 517 | 19204 | 3.62 | 0.999998081 | 1 | 1 |
| GOTERM_BP_DIRECT | GO:0008284~positive regulation of cell proliferation | 4 | 8.89 | 0.093225812 | 39 | 550 | 19308 | 3.60 | 1 | 1 | 1 |
| GOTERM_BP_DIRECT | GO:0008344~adult locomotory behavior | 2 | 4.44 | 0.093919474 | 39 | 50 | 19308 | 19.80 | 1 | 1 | 1 |
| GOTERM_MF_DIRECT | GO:0070888~E-box binding | 2 | 4.44 | 0.09417202 | 39 | 49 | 18869 | 19.75 | 0.999995283 | 1 | 1 |
| INTERPRO | IPR003598:Immunoglobulin subtype 2 | 3 | 6.67 | 0.097994065 | 41 | 254 | 19204 | 5.53 | 0.99999919 | 1 | 1 |

**Table S5 Gene ontology (GO) terms and statistics for differentially expressed genes (FDR 7%) in ZNF804A$^{-/-}$ neurons.** Green colour-coding identifies terms plotted in dotplot (FDR < 0.05).

**Table S6 Log2-transformed label free quantitation (LFQ) scores for 5393 identified proteins in each sample.**

| ID | logFC | AveExpr | t | P.Value | adj.P.Val | B |
|---|---|---|---|---|---|---|
| SERPINC1 | 4.37 | 12.22 | 10.27 | 1.16E-11 | 3.41E-10 | 16.58 |
| DNAH8 | 3.71 | 14.18 | 9.02 | 2.62E-10 | 4.26E-09 | 13.46 |
| MYOD1 | 3.63 | 12.00 | 3.49 | 0.003558966 | 0.007598053 | -2.00 |
| ALB | 3.61 | 19.77 | 8.87 | 3.86E-10 | 5.92E-09 | 13.07 |
| H2AC20;H2AC19 | 3.50 | 14.68 | 6.03 | 9.79E-07 | 5.10E-06 | 5.25 |
| HPX | 3.45 | 11.57 | 3.50 | 0.002272567 | 0.005146582 | -1.93 |
| JCHAIN | 3.16 | 8.32 | 2.47 | 0.029191517 | 0.048813568 | -4.07 |
| CDC40 | 3.09 | 14.82 | 5.74 | 3.23E-06 | 1.47E-05 | 4.16 |
| INS-IGF2 | 3.02 | 12.30 | 4.12 | 0.000288355 | 0.000820257 | -0.28 |
| ABCD3 | 2.97 | 10.52 | 5.42 | 6.37E-06 | 2.69E-05 | 3.41 |
| TFAP2D | 2.77 | 11.44 | 4.41 | 0.000129633 | 0.000400408 | 0.50 |
| FN1 | 2.68 | 7.64 | 3.17 | 0.00683182 | 0.01346881 | -2.79 |
| MYEF2 | 2.59 | 12.04 | 3.64 | 0.001245063 | 0.002985652 | -1.58 |
| TF | 2.50 | 14.26 | 8.03 | 3.57E-09 | 3.92E-08 | 10.85 |
| CINP | 2.50 | 9.68 | 6.85 | 6.85E-07 | 3.69E-06 | 5.98 |
| ZNF512 | 2.44 | 11.69 | 9.14 | 1.90E-10 | 3.22E-09 | 13.78 |
| TYMP | 2.38 | 11.47 | 2.52 | 0.017893063 | 0.031792409 | -4.16 |
| SCAF1 | 2.37 | 9.29 | 7.23 | 5.25E-07 | 2.94E-06 | 6.31 |
| HP1BP3 | 2.27 | 11.49 | 15.58 | 1.65E-16 | 1.24E-13 | 27.63 |
| MED25 | 2.26 | 8.98 | 4.61 | 0.00013423 | 0.000411942 | 0.69 |
| PDCD11 | 2.23 | 9.13 | 7.22 | 2.36E-07 | 1.47E-06 | 6.98 |
| BRIX1 | 2.23 | 10.42 | 4.59 | 6.55E-05 | 0.000218299 | 1.09 |
| PODXL | 2.21 | 12.84 | 11.66 | 4.59E-13 | 2.58E-11 | 19.79 |
| RRP1B | 2.20 | 11.45 | 4.45 | 9.77E-05 | 0.000312001 | 0.70 |
| NUP210 | 2.19 | 10.22 | 10.10 | 2.50E-11 | 6.29E-10 | 15.84 |
| SCYL3 | 2.19 | 8.90 | 2.66 | 0.014167801 | 0.025927422 | -3.78 |
| CPSF4 | 2.17 | 10.39 | 4.08 | 0.000376237 | 0.001035491 | -0.44 |
| ARL6IP4 | 2.14 | 8.75 | 5.26 | 2.44E-05 | 8.91E-05 | 2.34 |
| FGG | 2.11 | 9.61 | 3.33 | 0.003065945 | 0.00667294 | -2.37 |
| SDAD1 | 2.10 | 7.90 | 4.96 | 9.99E-05 | 0.000318181 | 1.17 |
| NCOA6 | 2.07 | 8.55 | 3.35 | 0.002637705 | 0.005842539 | -2.28 |
| ALDH1A1 | 2.06 | 9.48 | 5.37 | 8.26E-06 | 3.39E-05 | 3.18 |
| PRPF38B | 2.05 | 9.95 | 4.92 | 5.65E-05 | 0.000191873 | 1.52 |
| BAZ1B | 1.99 | 10.54 | 5.72 | 2.71E-06 | 1.26E-05 | 4.26 |
| INTS5 | 1.99 | 10.23 | 8.93 | 3.32E-10 | 5.22E-09 | 13.22 |
| PRPF6 | 1.99 | 11.62 | 4.05 | 0.000303514 | 0.00085873 | -0.41 |
| NOP56 | 1.96 | 11.85 | 11.38 | 8.56E-13 | 4.13E-11 | 19.17 |
| GLE1 | 1.94 | 10.83 | 11.03 | 1.61E-11 | 4.46E-10 | 16.40 |
| TCOF1 | 1.94 | 11.79 | 3.05 | 0.004601734 | 0.009487156 | -3.01 |
| LIMK2 | 1.94 | 7.36 | 2.75 | 0.016628692 | 0.029797799 | -3.60 |
| NUMA1 | 1.92 | 11.28 | 2.80 | 0.00859338 | 0.01663582 | -3.60 |
| U2AF1 | 1.92 | 13.26 | 3.47 | 0.001509601 | 0.003559894 | -1.96 |
| ZBTB34 | 1.92 | 11.51 | 3.87 | 0.000882879 | 0.002190107 | -1.12 |
| PRCC | 1.90 | 12.55 | 9.12 | 2.00E-10 | 3.34E-09 | 13.73 |
| NOP9 | 1.89 | 8.47 | 5.04 | 6.17E-05 | 0.000207833 | 1.58 |
| ZMYM3 | 1.89 | 9.36 | 4.62 | 9.18E-05 | 0.000295444 | 0.93 |
| NOC3L | 1.89 | 9.42 | 5.87 | 5.50E-06 | 2.37E-05 | 3.83 |
| H1-5 | 1.88 | 14.51 | 9.59 | 6.22E-11 | 1.28E-09 | 14.90 |
| MEX3B | 1.84 | 8.04 | 4.05 | 0.000624862 | 0.001630911 | -0.71 |

| | | | | | | |
|---|---|---|---|---|---|---|
| ZMYM4 | 1.84 | 9.47 | 6.14 | 1.47E-06 | 7.25E-06 | 5.00 |
| PHF14 | 1.83 | 10.28 | 3.21 | 0.003031159 | 0.006605437 | -2.62 |
| DDX51 | 1.83 | 6.88 | 3.07 | 0.008261524 | 0.016040668 | -2.97 |
| TOP1 | 1.83 | 12.26 | 10.29 | 1.10E-11 | 3.29E-10 | 16.63 |
| CDK7 | 1.81 | 9.66 | 6.70 | 2.36E-07 | 1.47E-06 | 6.75 |
| POU4F1 | 1.81 | 8.80 | 4.76 | 6.89E-05 | 0.000227793 | 1.26 |
| CDK1 | 1.79 | 9.63 | 7.71 | 1.06E-08 | 9.80E-08 | 9.79 |
| CTSZ | 1.79 | 10.73 | 5.92 | 1.75E-06 | 8.50E-06 | 4.73 |
| H3-7 | 1.78 | 17.07 | 9.35 | 1.13E-10 | 2.09E-09 | 14.30 |
| INO80 | 1.78 | 12.74 | 6.20 | 6.96E-07 | 3.73E-06 | 5.61 |
| HMBOX1 | 1.77 | 6.41 | 4.03 | 0.001427356 | 0.003382647 | -1.19 |
| RRS1 | 1.77 | 10.16 | 6.28 | 6.31E-07 | 3.46E-06 | 5.74 |
| H1-10 | 1.76 | 13.45 | 6.98 | 6.58E-08 | 4.79E-07 | 7.94 |
| MAD1L1 | 1.75 | 11.00 | 3.52 | 0.001314738 | 0.003135558 | -1.82 |
| NUP42 | 1.74 | 9.54 | 2.51 | 0.01832817 | 0.032510535 | -4.19 |
| CENPV | 1.73 | 11.09 | 3.26 | 0.002622292 | 0.005813301 | -2.48 |
| APOB | 1.73 | 9.21 | 2.54 | 0.017240982 | 0.030748174 | -4.11 |
| UTP25 | 1.73 | 8.06 | 3.06 | 0.008433463 | 0.016344307 | -2.99 |
| TOP2B | 1.72 | 13.05 | 8.93 | 3.34E-10 | 5.22E-09 | 13.22 |
| SUN1 | 1.72 | 9.56 | 5.46 | 7.92E-06 | 3.28E-05 | 3.28 |
| CFAP20 | 1.72 | 11.37 | 7.54 | 1.69E-08 | 1.44E-07 | 9.33 |
| APOC3 | 1.71 | 11.15 | 3.02 | 0.005146429 | 0.010504792 | -3.08 |
| DDX24 | 1.70 | 9.76 | 6.26 | 1.13E-05 | 4.48E-05 | 3.47 |
| STAG1 | 1.70 | 10.26 | 3.44 | 0.001701328 | 0.003960467 | -2.05 |
| PNN | 1.70 | 13.01 | 3.49 | 0.001431561 | 0.003391083 | -1.90 |
| DEK | 1.70 | 12.73 | 4.30 | 0.000150646 | 0.000457512 | 0.27 |
| NHP2 | 1.68 | 12.15 | 7.28 | 2.81E-08 | 2.29E-07 | 8.79 |
| H1-4;H1-3;H1-2 | 1.68 | 15.74 | 9.23 | 1.52E-10 | 2.65E-09 | 14.00 |
| CREBRF | 1.66 | 13.32 | 2.78 | 0.008945229 | 0.017212996 | -3.63 |
| RER1 | 1.65 | 12.15 | 3.21 | 0.003000184 | 0.006549322 | -2.61 |
| DDX21 | 1.65 | 12.60 | 11.15 | 1.45E-12 | 6.46E-11 | 18.65 |
| LAS1L | 1.65 | 11.79 | 3.36 | 0.002024826 | 0.004633464 | -2.24 |
| ZCCHC17 | 1.65 | 10.35 | 10.77 | 3.48E-12 | 1.34E-10 | 17.77 |
| H4-16 | 1.64 | 17.82 | 7.80 | 6.67E-09 | 6.57E-08 | 10.22 |
| YTHDC1 | 1.64 | 10.53 | 3.45 | 0.001813976 | 0.004196668 | -2.03 |
| RBM27 | 1.64 | 9.57 | 6.50 | 3.42E-07 | 2.04E-06 | 6.35 |
| CCNK | 1.63 | 11.24 | 3.05 | 0.004550099 | 0.009397099 | -3.00 |
| SMPD4 | 1.63 | 9.78 | 3.65 | 0.001022544 | 0.002503469 | -1.51 |
| EBNA1BP2 | 1.62 | 10.81 | 8.01 | 3.79E-09 | 4.12E-08 | 10.79 |
| BRD1 | 1.62 | 9.83 | 5.76 | 2.75E-06 | 1.27E-05 | 4.28 |
| SNRNP27 | 1.62 | 10.78 | 2.71 | 0.010614767 | 0.020046723 | -3.79 |
| RBM39 | 1.61 | 14.31 | 12.86 | 3.34E-14 | 4.28E-12 | 22.40 |
| IGHG1 | 1.61 | 13.31 | 2.79 | 0.008764312 | 0.016910649 | -3.62 |
| KLHL7 | 1.61 | 8.52 | 3.11 | 0.00448471 | 0.009279952 | -2.83 |
| EMG1 | 1.61 | 10.47 | 7.84 | 7.49E-09 | 7.17E-08 | 10.14 |
| DDX18 | 1.60 | 10.58 | 6.96 | 6.87E-08 | 4.97E-07 | 7.89 |
| MPHOSPH6 | 1.60 | 8.38 | 4.41 | 0.000334614 | 0.000937132 | -0.04 |
| GTF2H4 | 1.60 | 8.35 | 2.80 | 0.010629103 | 0.020066585 | -3.51 |
| HIC2 | 1.60 | 11.11 | 9.50 | 7.70E-11 | 1.54E-09 | 14.68 |
| BRD3 | 1.60 | 11.03 | 9.96 | 3.46E-11 | 7.89E-10 | 15.51 |

| | | | | | | |
|---|---|---|---|---|---|---|
| RSBN1L | 1.60 | 9.23 | 5.43 | 2.57E-05 | 9.35E-05 | 2.42 |
| EMSY | 1.59 | 8.22 | 6.07 | 4.06E-06 | 1.81E-05 | 4.17 |
| CCDC183 | 1.59 | 16.07 | 6.99 | 1.09E-07 | 7.45E-07 | 7.53 |
| KRT3 | 1.58 | 13.94 | 2.55 | 0.015640803 | 0.02829779 | -4.15 |
| FUCA1 | 1.58 | 11.63 | 2.89 | 0.006836235 | 0.013472461 | -3.38 |
| ZBTB10 | 1.57 | 8.11 | 2.74 | 0.012265666 | 0.022792479 | -3.63 |
| SLC35E1 | 1.57 | 8.42 | 3.79 | 0.001220581 | 0.002934982 | -1.35 |
| CLN5 | 1.57 | 10.23 | 2.88 | 0.007047017 | 0.013851487 | -3.41 |
| TMPO | 1.57 | 13.90 | 10.23 | 1.28E-11 | 3.68E-10 | 16.48 |
| ZNF346 | 1.55 | 9.54 | 3.49 | 0.003564692 | 0.007607187 | -2.16 |
| SYMPK | 1.55 | 11.82 | 6.74 | 1.30E-07 | 8.66E-07 | 7.26 |
| TRIM33 | 1.55 | 10.37 | 2.46 | 0.019272173 | 0.033898893 | -4.34 |
| ZC3H11A | 1.55 | 10.76 | 8.20 | 2.86E-09 | 3.26E-08 | 11.10 |
| TERF2 | 1.55 | 8.92 | 3.97 | 0.00041359 | 0.001127076 | -0.66 |
| RBM12B | 1.55 | 12.04 | 11.54 | 6.03E-13 | 3.26E-11 | 19.52 |
| ATOX1 | 1.55 | 9.78 | 2.74 | 0.012238367 | 0.022753143 | -3.56 |
| XAB2 | 1.54 | 9.46 | 5.74 | 3.67E-06 | 1.65E-05 | 4.06 |
| FTSJ3 | 1.54 | 7.46 | 4.42 | 0.000197306 | 0.000580754 | 0.29 |
| ADAR | 1.54 | 12.93 | 9.02 | 2.61E-10 | 4.26E-09 | 13.46 |
| ZNF423 | 1.54 | 7.73 | 2.78 | 0.015504607 | 0.028109457 | -3.38 |
| INTS8 | 1.53 | 9.65 | 5.71 | 3.13E-06 | 1.43E-05 | 4.15 |
| CAT | 1.53 | 10.89 | 4.18 | 0.000207951 | 0.000610037 | -0.04 |
| SAFB2 | 1.53 | 12.03 | 4.44 | 9.97E-05 | 0.000317584 | 0.68 |
| COIL | 1.52 | 9.36 | 5.28 | 1.79E-05 | 6.76E-05 | 2.58 |
| NUP35 | 1.52 | 11.30 | 5.62 | 3.25E-06 | 1.48E-05 | 4.05 |
| KRT5 | 1.51 | 13.86 | 2.65 | 0.012288776 | 0.022814568 | -3.93 |
| COMMD3-BMI1 | 1.51 | 11.29 | 9.92 | 3.81E-11 | 8.56E-10 | 15.42 |
| THOC1 | 1.50 | 10.92 | 3.64 | 0.000958922 | 0.002363122 | -1.52 |
| PRR36 | -1.50 | 11.68 | -5.84 | 1.92E-06 | 9.29E-06 | 4.60 |
| KIAA0513 | -1.50 | 10.86 | -11.68 | 6.72E-13 | 3.50E-11 | 19.44 |
| AS3MT | -1.50 | 9.44 | -3.41 | 0.00261326 | 0.00580062 | -2.04 |
| NIBAN2 | -1.51 | 9.47 | -3.04 | 0.00508685 | 0.010387217 | -3.00 |
| PPP1R17 | -1.51 | 12.94 | -3.03 | 0.004864521 | 0.009975875 | -3.07 |
| ATG12 | -1.51 | 8.19 | -2.82 | 0.010879912 | 0.020481211 | -3.36 |
| MAPK8IP3 | -1.51 | 11.82 | -6.30 | 4.55E-07 | 2.59E-06 | 6.01 |
| HEBP2 | -1.52 | 10.95 | -6.51 | 3.96E-07 | 2.29E-06 | 6.25 |
| MAPT | -1.52 | 15.00 | -8.95 | 3.16E-10 | 5.03E-09 | 13.27 |
| RAB3B | -1.52 | 10.37 | -6.04 | 2.17E-06 | 1.03E-05 | 4.66 |
| CARHSP1 | -1.52 | 12.90 | -3.07 | 0.004307434 | 0.00896998 | -2.95 |
| ATE1 | -1.53 | 9.59 | -2.98 | 0.00541148 | 0.01097866 | -3.17 |
| RAB3D | -1.53 | 11.10 | -3.40 | 0.002102692 | 0.004801192 | -2.12 |
| BAG3 | -1.53 | 9.43 | -2.52 | 0.018295653 | 0.032463816 | -4.12 |
| RAP2C | -1.53 | 11.47 | -2.65 | 0.014757571 | 0.026912974 | -3.75 |
| PHPT1 | -1.53 | 9.72 | -3.29 | 0.003229105 | 0.006978904 | -2.41 |
| DOCK6 | -1.53 | 10.58 | -4.00 | 0.000399537 | 0.001093316 | -0.59 |
| PPM1H | -1.54 | 10.10 | -7.39 | 3.80E-08 | 2.95E-07 | 8.59 |
| CDH6 | -1.54 | 10.19 | -2.64 | 0.012710292 | 0.023547206 | -3.96 |
| ARFGAP1 | -1.54 | 11.06 | -3.90 | 0.000523077 | 0.001392929 | -0.85 |
| ARHGEF10L | -1.54 | 9.19 | -2.77 | 0.010204301 | 0.019323693 | -3.60 |
| AGTPBP1 | -1.54 | 9.62 | -2.40 | 0.023560168 | 0.040341424 | -4.40 |

| | | | | | | |
|---|---|---|---|---|---|---|
| MVK | -1.55 | 11.39 | -5.30 | 8.37E-06 | 3.43E-05 | 3.12 |
| TUBA4A | -1.55 | 15.37 | -8.19 | 2.33E-09 | 2.73E-08 | 11.27 |
| TUBAL3 | -1.55 | 15.28 | -2.98 | 0.005401324 | 0.010965438 | -3.16 |
| GPC1 | -1.55 | 11.02 | -2.73 | 0.010206176 | 0.019323693 | -3.76 |
| MYCBP2 | -1.55 | 11.30 | -10.67 | 4.46E-12 | 1.59E-10 | 17.53 |
| YES1 | -1.55 | 10.30 | -7.51 | 1.63E-07 | 1.06E-06 | 7.44 |
| DBN1 | -1.55 | 10.70 | -3.43 | 0.002404712 | 0.005401607 | -2.03 |
| ABCF3 | -1.56 | 11.25 | -3.32 | 0.002224322 | 0.005046023 | -2.33 |
| ARPC1A | -1.57 | 14.91 | -4.36 | 0.000127594 | 0.000395271 | 0.44 |
| PPP1R1A | -1.57 | 9.91 | -3.43 | 0.00239817 | 0.005391521 | -2.07 |
| PDLIM4 | -1.57 | 9.90 | -4.05 | 0.00040758 | 0.001112429 | -0.51 |
| PTPRA | -1.57 | 10.01 | -2.66 | 0.012201893 | 0.022701396 | -3.90 |
| RAP1GAP2 | -1.57 | 10.42 | -5.35 | 1.19E-05 | 4.70E-05 | 2.94 |
| TENM3 | -1.58 | 10.89 | -4.23 | 0.000346418 | 0.000967099 | -0.21 |
| GCLM | -1.58 | 10.69 | -2.46 | 0.020038464 | 0.035000696 | -4.30 |
| SNX4 | -1.58 | 12.18 | -3.40 | 0.001805839 | 0.004181524 | -2.13 |
| UNC13A | -1.59 | 9.58 | -4.56 | 0.000125158 | 0.000388642 | 0.73 |
| TRAPPC9 | -1.59 | 9.40 | -2.56 | 0.015849408 | 0.028606248 | -4.10 |
| ADD3 | -1.59 | 9.51 | -5.87 | 2.95E-06 | 1.36E-05 | 4.33 |
| LIMA1 | -1.60 | 12.62 | -10.67 | 4.45E-12 | 1.59E-10 | 17.53 |
| ZP1 | -1.61 | 11.75 | -7.16 | 5.64E-08 | 4.18E-07 | 8.16 |
| EVL | -1.61 | 10.16 | -2.84 | 0.008209818 | 0.015946168 | -3.48 |
| POTEI | -1.61 | 13.28 | -3.66 | 0.000956865 | 0.002359837 | -1.47 |
| KATNB1 | -1.62 | 11.52 | -2.71 | 0.010895814 | 0.020503799 | -3.77 |
| PCSK1 | -1.62 | 10.36 | -2.37 | 0.023823411 | 0.040734159 | -4.53 |
| SNX15 | -1.62 | 10.88 | -2.94 | 0.006112427 | 0.012201631 | -3.26 |
| WASF2 | -1.63 | 10.70 | -2.60 | 0.014274837 | 0.026114203 | -4.04 |
| VPS50 | -1.63 | 9.64 | -2.89 | 0.007409081 | 0.014514285 | -3.37 |
| NAV2 | -1.63 | 10.71 | -6.26 | 7.86E-07 | 4.17E-06 | 5.56 |
| LRCH2 | -1.63 | 11.09 | -2.91 | 0.006481616 | 0.012855572 | -3.33 |
| LPIN1 | -1.63 | 8.60 | -3.65 | 0.001708298 | 0.003973173 | -1.52 |
| PIP4K2A | -1.63 | 12.02 | -3.24 | 0.002791653 | 0.00613261 | -2.54 |
| F8A3 | -1.63 | 9.51 | -2.58 | 0.015747764 | 0.028471697 | -4.02 |
| GSTCD | -1.63 | 8.80 | -3.63 | 0.001657403 | 0.003874313 | -1.58 |
| PPP6R3 | -1.63 | 11.48 | -2.36 | 0.024269024 | 0.041358889 | -4.55 |
| MAPK7 | -1.64 | 9.97 | -3.70 | 0.0008569 | 0.002135745 | -1.36 |
| BICD1 | -1.64 | 12.08 | -5.02 | 1.84E-05 | 6.95E-05 | 2.34 |
| MAP7D1 | -1.64 | 12.52 | -4.58 | 6.69E-05 | 0.000222175 | 1.07 |
| TUBB6 | -1.65 | 11.35 | -6.82 | 1.19E-07 | 8.10E-07 | 7.37 |
| PFKL | -1.65 | 12.13 | -3.77 | 0.000660417 | 0.001710962 | -1.16 |
| SNAP25 | -1.65 | 12.57 | -5.25 | 9.46E-06 | 3.81E-05 | 3.00 |
| CBLB | -1.65 | 10.30 | -2.91 | 0.006850896 | 0.013496291 | -3.32 |
| STX2 | -1.66 | 10.15 | -3.60 | 0.001073386 | 0.002616969 | -1.63 |
| CFL2 | -1.66 | 12.15 | -7.84 | 3.32E-08 | 2.63E-07 | 8.88 |
| ABHD14B | -1.66 | 9.22 | -2.60 | 0.019138769 | 0.033732 | -3.75 |
| DSTN | -1.66 | 14.43 | -8.42 | 1.25E-09 | 1.59E-08 | 11.89 |
| STX18 | -1.67 | 9.57 | -3.21 | 0.004827003 | 0.009906669 | -2.56 |
| TMEM169 | -1.67 | 12.17 | -6.58 | 2.02E-07 | 1.29E-06 | 6.82 |
| LYPLA2 | -1.67 | 12.42 | -3.76 | 0.000705158 | 0.001804627 | -1.20 |
| DOCK4 | -1.67 | 8.74 | -2.53 | 0.017119993 | 0.030553141 | -4.14 |

| | | | | | | |
|---|---|---|---|---|---|---|
| MAP6 | -1.68 | 12.62 | -10.06 | 1.94E-11 | 5.24E-10 | 16.06 |
| UBE2G1 | -1.69 | 10.01 | -3.90 | 0.000829567 | 0.002077476 | -0.98 |
| PPP1R14B | -1.70 | 10.94 | -2.67 | 0.012288078 | 0.022814568 | -3.86 |
| PSME2 | -1.70 | 11.18 | -3.78 | 0.000672556 | 0.001736419 | -1.15 |
| FNBP1 | -1.71 | 9.48 | -2.58 | 0.01555365 | 0.02816921 | -4.04 |
| FNTA | -1.71 | 10.85 | -2.66 | 0.012795763 | 0.023680499 | -3.86 |
| KCTD10 | -1.71 | 9.47 | -4.13 | 0.000293922 | 0.000833387 | -0.26 |
| EIF5A | -1.72 | 13.64 | -3.88 | 0.00049341 | 0.001323317 | -0.88 |
| ITGA11 | -1.72 | 11.22 | -7.59 | 6.00E-08 | 4.42E-07 | 8.29 |
| STMN3 | -1.72 | 11.98 | -5.71 | 3.50E-06 | 1.58E-05 | 4.08 |
| PIR | -1.73 | 8.78 | -4.27 | 0.000517891 | 0.001380518 | -0.22 |
| CDV3 | -1.73 | 12.55 | -3.64 | 0.000949873 | 0.002346325 | -1.51 |
| PIN4 | -1.73 | 11.08 | -4.09 | 0.000274317 | 0.000784148 | -0.31 |
| SNTB2 | -1.73 | 10.08 | -4.59 | 0.000129004 | 0.000398934 | 0.76 |
| SLC18A3 | -1.74 | 11.69 | -3.42 | 0.001953425 | 0.00448963 | -2.09 |
| DAB2IP | -1.75 | 10.89 | -2.92 | 0.006586225 | 0.013033532 | -3.30 |
| CEP41 | -1.75 | 11.30 | -7.44 | 2.18E-08 | 1.81E-07 | 9.07 |
| GYPC | -1.75 | 9.19 | -2.54 | 0.021056225 | 0.036583798 | -3.77 |
| KIFAP3 | -1.75 | 10.52 | -7.70 | 1.09E-08 | 9.99E-08 | 9.77 |
| GNPNAT1 | -1.77 | 9.07 | -4.14 | 0.000465037 | 0.001255551 | -0.45 |
| NQO2 | -1.77 | 11.92 | -4.37 | 0.000121528 | 0.000379387 | 0.48 |
| EDIL3 | -1.78 | 15.58 | -14.55 | 1.12E-15 | 4.21E-13 | 25.75 |
| FSD1L | -1.78 | 12.20 | -4.73 | 4.36E-05 | 0.000151478 | 1.49 |
| SEMA3C | -1.79 | 11.94 | -9.18 | 3.21E-10 | 5.09E-09 | 13.32 |
| NUMB | -1.80 | 8.39 | -3.78 | 0.001494508 | 0.003529055 | -1.14 |
| ENO2 | -1.80 | 13.96 | -8.68 | 6.31E-10 | 8.89E-09 | 12.58 |
| SHMT1 | -1.80 | 8.85 | -2.61 | 0.015578202 | 0.028203953 | -3.86 |
| UBE2L3 | -1.81 | 12.29 | -2.77 | 0.009272299 | 0.017760357 | -3.67 |
| RALGAPA1 | -1.81 | 10.43 | -6.81 | 1.24E-07 | 8.35E-07 | 7.33 |
| ZC2HC1A | -1.81 | 12.20 | -4.78 | 3.78E-05 | 0.000132387 | 1.63 |
| TMEM120B | -1.82 | 6.98 | -2.52 | 0.023522664 | 0.040309224 | -3.92 |
| DISP2 | -1.82 | 9.75 | -3.65 | 0.00096633 | 0.002380262 | -1.50 |
| TUBB8 | -1.82 | 15.53 | -4.07 | 0.000288713 | 0.000820832 | -0.36 |
| SNX24 | -1.83 | 8.74 | -3.77 | 0.001660829 | 0.003879946 | -1.41 |
| ILVBL | -1.83 | 10.15 | -3.14 | 0.004296017 | 0.008953301 | -2.76 |
| SCRN1 | -1.84 | 13.86 | -3.48 | 0.001460871 | 0.003454282 | -1.92 |
| CALD1 | -1.84 | 12.36 | -12.69 | 4.81E-14 | 5.18E-12 | 22.04 |
| LAMA1 | -1.84 | 11.02 | -2.97 | 0.007866767 | 0.015336547 | -2.75 |
| SH3PXD2A | -1.86 | 9.69 | -3.87 | 0.000587448 | 0.001543997 | -0.93 |
| C15orf38-AP3S2;ARPIN | -1.87 | 10.78 | -5.28 | 8.85E-06 | 3.60E-05 | 3.06 |
| MAPK8IP2 | -1.87 | 10.44 | -2.88 | 0.009254384 | 0.017732507 | -3.16 |
| HSPB11 | -1.88 | 9.85 | -5.54 | 9.27E-06 | 3.75E-05 | 3.26 |
| PTGR1 | -1.89 | 11.01 | -3.19 | 0.003278179 | 0.007055941 | -2.64 |
| ACTN2 | -1.89 | 11.09 | -2.62 | 0.013731066 | 0.025163245 | -3.96 |
| KYAT1 | -1.90 | 9.05 | -3.15 | 0.0044391 | 0.009200408 | -2.68 |
| NUDT4B;NUDT4 | -1.90 | 10.12 | -2.62 | 0.013245826 | 0.024427368 | -4.00 |
| CDK16 | -1.90 | 9.82 | -3.21 | 0.003238849 | 0.00699135 | -2.61 |
| RAB35 | -1.90 | 11.55 | -4.53 | 9.29E-05 | 0.000298836 | 0.83 |
| MSTO1 | -1.92 | 9.66 | -2.58 | 0.01606071 | 0.028927998 | -3.96 |
| PARVA | -1.93 | 10.06 | -3.18 | 0.003474673 | 0.007436225 | -2.68 |

| | | | | | | |
|---|---|---|---|---|---|---|
| PLCB3 | -1.94 | 9.18 | -2.73 | 0.013335871 | 0.024576172 | -3.48 |
| LLGL1 | -1.95 | 10.64 | -3.71 | 0.000781582 | 0.001971403 | -1.32 |
| APLP2 | -1.95 | 9.99 | -4.82 | 8.16E-05 | 0.000267099 | 1.24 |
| SYNJ2BP-COX16 | -1.96 | 8.45 | -3.76 | 0.002358775 | 0.005321169 | -1.56 |
| PPP4R1 | -1.96 | 11.01 | -4.41 | 0.000114097 | 0.000359177 | 0.57 |
| LRRC20 | -1.97 | 10.93 | -4.15 | 0.000242845 | 0.000699894 | -0.16 |
| AHCYL2 | -1.97 | 11.05 | -3.27 | 0.002582701 | 0.005744925 | -2.47 |
| EFHD2 | -1.97 | 10.27 | -9.80 | 7.09E-11 | 1.43E-09 | 14.83 |
| TUBB4A | -1.97 | 13.99 | -5.24 | 9.74E-06 | 3.92E-05 | 2.97 |
| RBM43 | -1.98 | 10.85 | -3.25 | 0.003118018 | 0.006764682 | -2.53 |
| VCAN | -1.98 | 10.24 | -6.77 | 6.53E-07 | 3.56E-06 | 6.01 |
| TOMM34 | -1.98 | 9.30 | -2.30 | 0.02820705 | 0.047348192 | -4.67 |
| ADA | -1.98 | 8.35 | -4.58 | 0.000231247 | 0.000670145 | 0.46 |
| KIF3B | -1.98 | 9.58 | -5.68 | 5.64E-06 | 2.43E-05 | 3.71 |
| PPP1R9A | -2.00 | 7.37 | -2.46 | 0.023297766 | 0.039989044 | -4.14 |
| ATG4B | -2.01 | 10.52 | -4.88 | 5.01E-05 | 0.000171852 | 1.59 |
| PPP3R1 | -2.02 | 11.16 | -3.00 | 0.005183523 | 0.010572294 | -3.13 |
| TGM1 | -2.03 | 10.08 | -2.42 | 0.024244765 | 0.041330953 | -4.28 |
| PYGM | -2.03 | 12.52 | -2.34 | 0.026185156 | 0.044251145 | -4.58 |
| GPC4 | -2.04 | 9.10 | -3.33 | 0.002532314 | 0.00564719 | -2.32 |
| FLAD1 | -2.05 | 10.84 | -2.40 | 0.022255341 | 0.038476329 | -4.47 |
| CBARP | -2.05 | 10.69 | -3.24 | 0.002792007 | 0.00613261 | -2.54 |
| BRK1 | -2.06 | 9.47 | -2.94 | 0.010755205 | 0.020275511 | -2.74 |
| UBA5 | -2.07 | 10.73 | -4.50 | 9.51E-05 | 0.000304659 | 0.78 |
| DCX | -2.08 | 9.93 | -4.06 | 0.000604713 | 0.00158304 | -0.60 |
| MRPL2 | -2.08 | 9.53 | -2.41 | 0.022505573 | 0.038812154 | -4.44 |
| PHYHIPL | -2.08 | 11.12 | -3.35 | 0.002084897 | 0.004766774 | -2.26 |
| SEMA6D | -2.09 | 10.44 | -2.98 | 0.005573207 | 0.011279518 | -3.17 |
| ATP11C | -2.09 | 6.43 | -2.35 | 0.026900826 | 0.045358453 | -4.49 |
| ADAMTS4 | -2.09 | 9.38 | -5.41 | 1.46E-05 | 5.62E-05 | 2.89 |
| CFAP410 | -2.09 | 9.28 | -3.33 | 0.005930116 | 0.011891919 | -2.14 |
| SMAD1 | -2.11 | 9.72 | -2.82 | 0.008331685 | 0.016158979 | -3.53 |
| GBE1 | -2.11 | 9.68 | -3.03 | 0.005038282 | 0.010308074 | -3.05 |
| VPS37B | -2.12 | 6.94 | -2.58 | 0.025693968 | 0.043547132 | -3.67 |
| GRK5 | -2.12 | 10.38 | -2.98 | 0.005656402 | 0.011412725 | -3.16 |
| PPP1CB | -2.12 | 12.91 | -4.22 | 0.000195021 | 0.000575641 | 0.05 |
| LRCH1 | -2.14 | 10.77 | -2.52 | 0.017445382 | 0.031081057 | -4.20 |
| ATP6V1G2 | -2.14 | 11.38 | -2.99 | 0.005323166 | 0.010823496 | -3.15 |
| TBCE | -2.15 | 10.72 | -3.31 | 0.002309751 | 0.00521954 | -2.36 |
| ACTR1B | -2.17 | 11.30 | -2.86 | 0.007439058 | 0.014567578 | -3.46 |
| BDNF | -2.17 | 11.45 | -3.05 | 0.00508679 | 0.010387217 | -2.96 |
| SUSD2 | -2.19 | 11.71 | -3.87 | 0.000502921 | 0.001346075 | -0.90 |
| PRUNE1 | -2.19 | 10.24 | -3.81 | 0.000623163 | 0.001627286 | -1.08 |
| NUBP2 | -2.19 | 10.09 | -3.51 | 0.001664653 | 0.003883698 | -1.89 |
| CZIB | -2.22 | 10.91 | -2.99 | 0.005258251 | 0.010708082 | -3.14 |
| PAK5 | -2.23 | 8.11 | -2.48 | 0.022415209 | 0.038689064 | -3.96 |
| TRAPPC2; TRAPPC2B | -2.24 | 8.21 | -3.73 | 0.002229563 | 0.005055729 | -1.44 |
| JPT1 | -2.28 | 11.51 | -4.30 | 0.000155316 | 0.000469794 | 0.27 |
| SMYD5 | -2.28 | 10.38 | -7.86 | 4.21E-08 | 3.26E-07 | 8.70 |
| SYT5 | -2.28 | 9.31 | -3.46 | 0.00201206 | 0.004612287 | -1.97 |

| | | | | | | |
|---|---|---|---|---|---|---|
| PRXL2B | -2.29 | 8.21 | -3.45 | 0.004260153 | 0.008892668 | -2.03 |
| FNTB | -2.34 | 10.21 | -3.59 | 0.001206779 | 0.002905782 | -1.66 |
| AFAP1 | -2.34 | 12.29 | -6.82 | 1.01E-07 | 7.00E-07 | 7.50 |
| FAM89B | -2.35 | 9.97 | -2.71 | 0.013546204 | 0.024902643 | -3.63 |
| NAPRT | -2.36 | 8.94 | -4.44 | 0.000171724 | 0.000515859 | 0.45 |
| RABEP2 | -2.36 | 8.83 | -7.35 | 2.30E-07 | 1.45E-06 | 7.12 |
| POLR3D | -2.38 | 15.02 | -4.76 | 6.98E-05 | 0.000230659 | 1.26 |
| MAGI1 | -2.38 | 8.93 | -3.62 | 0.00110097 | 0.002675529 | -1.57 |
| KCMF1 | -2.39 | 9.39 | -3.22 | 0.00380954 | 0.008057698 | -2.53 |
| PLIN3 | -2.40 | 10.26 | -3.84 | 0.00059815 | 0.001568988 | -1.01 |
| SGIP1 | -2.40 | 9.88 | -4.14 | 0.000348076 | 0.000970696 | -0.29 |
| MADD | -2.41 | 9.93 | -3.50 | 0.001508228 | 0.003558253 | -1.88 |
| STUM | -2.43 | 8.58 | -3.66 | 0.001308695 | 0.003123981 | -1.51 |
| NPDC1 | -2.45 | 7.96 | -2.50 | 0.022066816 | 0.038200675 | -3.93 |
| PITPNA | -2.49 | 11.18 | -3.23 | 0.002935381 | 0.006420688 | -2.56 |
| ATG3 | -2.51 | 10.83 | -3.87 | 0.000505776 | 0.001353029 | -0.90 |
| FAM107B | -2.51 | 9.51 | -3.57 | 0.001184933 | 0.002859466 | -1.70 |
| VASP | -2.58 | 10.64 | -3.75 | 0.000760958 | 0.001927711 | -1.24 |
| NRP1 | -2.59 | 9.28 | -3.01 | 0.00563962 | 0.011387612 | -3.06 |
| ACOT7L | -2.65 | 13.43 | -4.52 | 0.000140321 | 0.00042863 | 0.60 |
| KIF21B | -2.68 | 10.85 | -4.41 | 0.000109435 | 0.000345122 | 0.59 |
| BPGM | -2.68 | 8.42 | -2.41 | 0.024922721 | 0.042376691 | -4.16 |
| SH3BP5L | -2.69 | 8.75 | -4.06 | 0.000425123 | 0.001156107 | -0.51 |
| CCDC184 | -2.69 | 9.32 | -2.86 | 0.009649919 | 0.018376468 | -3.20 |
| ACVR1B | -2.72 | 6.94 | -3.63 | 0.004599552 | 0.009487156 | -1.83 |
| SPAST | -2.73 | 11.05 | -3.68 | 0.000980013 | 0.00240832 | -1.43 |
| RHOA | -2.77 | 11.93 | -6.82 | 2.14E-06 | 1.02E-05 | 5.08 |
| SLC23A2 | -2.78 | 6.90 | -2.65 | 0.020013834 | 0.034980932 | -3.63 |
| NUDT14 | -2.78 | 9.48 | -5.58 | 3.27E-05 | 0.000116512 | 2.48 |
| KDELR1 | -2.83 | 10.18 | -2.46 | 0.021007029 | 0.036510397 | -4.25 |
| SPIRE1 | -2.84 | 9.07 | -4.22 | 0.000649744 | 0.00168743 | -0.20 |
| ADI1 | -2.91 | 8.69 | -4.61 | 0.000480963 | 0.001294559 | 0.08 |
| PIGU | -2.91 | 9.93 | -3.38 | 0.002360209 | 0.005322119 | -2.17 |
| PLSCR3 | -2.92 | 7.38 | -3.05 | 0.007143803 | 0.014036478 | -2.85 |
| PGK2 | -2.94 | 13.51 | -3.97 | 0.000400465 | 0.001095286 | -0.65 |
| RPL15 | -3.09 | 11.18 | -2.81 | 0.009621414 | 0.018335476 | -3.49 |
| CYBC1 | -3.21 | 6.78 | -2.62 | 0.027847583 | 0.046789639 | -3.66 |
| PLEKHO2 | -3.47 | 8.54 | -4.03 | 0.000710379 | 0.001816219 | -0.67 |
| RPS6KA1 | -3.54 | -0.43 | -3.12 | 0.010812677 | 0.020376544 | -2.80 |
| SRGAP2 | -3.68 | 7.13 | -4.02 | 0.000988845 | 0.002428887 | -0.72 |
| CEP43 | -3.69 | -0.22 | -4.29 | 0.001578299 | 0.003701956 | -0.99 |
| HTD2 | -3.75 | 6.13 | -2.88 | 0.012789609 | 0.023677451 | -3.20 |
| FBXL16 | -3.86 | 7.32 | -4.13 | 0.000433377 | 0.001176487 | -0.43 |
| MYL6B | -3.99 | 9.85 | -4.73 | 4.38E-05 | 0.000152102 | 1.49 |
| FMNL1 | -4.20 | 4.24 | -3.30 | 0.009162413 | 0.017562685 | -2.58 |
| RANBP10 | -4.32 | 11.74 | -5.81 | 0.000250431 | 0.000720573 | 0.87 |
| A2ML1 | -4.34 | 15.19 | -2.49 | 0.022235801 | 0.038467863 | -3.96 |
| ADAL | -4.42 | 7.94 | -4.01 | 0.001006042 | 0.002465366 | -0.74 |
| SFRP1 | -4.58 | 10.35 | -7.16 | 2.69E-07 | 1.66E-06 | 6.91 |
| ROBO2 | -4.73 | 9.21 | -5.45 | 7.28E-06 | 3.04E-05 | 3.35 |
| CCN2 | -4.77 | 9.89 | -6.09 | 4.77E-06 | 2.10E-05 | 4.20 |
| AP3M2 | -5.07 | 8.12 | -4.29 | 0.00074688 | 0.001897538 | -0.54 |

**Table S7 Results of differential protein expression analysis according to cellular compartment.** Soma enriched proteins (p.adj. < 0.05, logFC > 1.5) listed on top, neurite enriched proteins (p.adj. < 0.05, logFC < -1.5) listed on bottom of table.

| Neurite-enriched proteins | | | | | | | | | | | |
|---|---|---|---|---|---|---|---|---|---|---|---|
| **Category** | **Term** | **Count** | **%** | **PValue** | **List Total** | **Pop Hits** | **Pop Total** | **Fold Enrichment** | **Bonferroni** | **Benjamini** | **FDR** |
| GOTERM_CC_DIRECT | Plasma Membrane | 83 | 34.58 | 1.15E-05 | 231 | 1202 | 5142 | 1.54 | 0.0040 | 0.0040 | 0.0040 |
| UP_KW_CELLULAR_COMPONENT | Cell Membrane | 48 | 20.00 | 0.0023 | 210 | 716 | 4751 | 1.52 | 0.0691 | 0.0179 | 0.0144 |
| UP_KW_MOLECULAR_FUNCTION | Actin-binding | 16 | 6.67 | 4.16E-04 | 138 | 133 | 3215 | 2.80 | 0.0222 | 0.0225 | 0.0225 |
| GOTERM_CC_DIRECT | Microtubule Cytoskeleton | 13 | 5.42 | 2.47E-04 | 231 | 82 | 5142 | 3.53 | 0.0822 | 0.0429 | 0.0427 |
| UP_KW_CELLULAR_COMPONENT | Microtubule | 13 | 5.42 | 0.0037 | 210 | 113 | 4751 | 2.60 | 0.1096 | 0.0227 | 0.0183 |
| **Soma-enriched proteins** | | | | | | | | | | | |
| **Category** | **Term** | **Count** | **%** | **PValue** | **List Total** | **Pop Hits** | **Pop Total** | **Fold Enrichment** | **Bonferroni** | **Benjamini** | **FDR** |
| UP_KW_PTM | Citrullination | 13 | 9.63 | 2.53E-10 | 123 | 40 | 4525 | 11.96 | 4.55E-09 | 4.80E-09 | 4.04E-09 |
| INTERPRO | Histone H5 | 6 | 4.444 | 2.15E-07 | 133 | 7 | 5109 | 32.93 | 6.24E-05 | 6.24E-05 | 6.24E-05 |
| INTERPRO | Histone H1/H5 | 6 | 4.444 | 5.62E-07 | 133 | 8 | 5109 | 28.81 | 1.63E-04 | 8.15E-05 | 8.15E-05 |
| UP_SEQ_FEATURE | H15 | 6 | 4.444 | 5.89E-07 | 134 | 8 | 5099 | 28.54 | 4.23E-04 | 1.41E-04 | 1.40E-04 |
| UP_KW_CELLULAR_COMPONENT | Chromosome | 21 | 15.56 | 9.38E-07 | 127 | 220 | 4751 | 3.57 | 3.10E-05 | 1.55E-05 | 1.41E-05 |
| GOTERM_MF_DIRECT | Structural Constituent of Chromatin | 8 | 5.926 | 2.20E-06 | 133 | 25 | 5087 | 12.24 | 5.12E-04 | 5.12E-04 | 5.04E-04 |
| GOTERM_BP_DIRECT | Nucleosome Positioning | 6 | 4.444 | 2.29E-06 | 129 | 10 | 5008 | 23.29 | 0.0017 | 0.0015 | 0.0015 |
| GOTERM_BP_DIRECT | Chromosome Condensation | 6 | 4.444 | 4.12E-06 | 129 | 11 | 5008 | 21.18 | 0.0030 | 0.0015 | 0.0015 |
| GOTERM_CC_DIRECT | Nucleosome | 9 | 6.667 | 1.48E-05 | 133 | 45 | 5142 | 7.73 | 0.0036 | 7.24E-04 | 7.03E-04 |
| GOTERM_BP_DIRECT | Negative regulation of DNA recombination | 5 | 3.704 | 4.65E-05 | 129 | 9 | 5008 | 21.57 | 0.0331 | 0.0093 | 0.0092 |
| UP_KW_PTM | Hydroxylation | 7 | 5.185 | 0.0061 | 123 | 62 | 4525 | 4.15 | 0.1040 | 0.0231 | 0.0195 |

**Table S8 Gene ontology (GO) terms and statistics for proteins enriched in neurites (top) and somata (bottom).**

| Analysis | Category | Term | Count | % | PValue | List Total | Pop Hits | Pop Total | Fold Enrichment | Bonferroni | Benjamini | FDR |
|---|---|---|---|---|---|---|---|---|---|---|---|---|
| wt vs. het | UP_KW_DOMAIN | Transit peptide | 11 | 9.09 | 2.29E-04 | 67 | 568 | 14504 | 4.19 | 0.0034 | 0.0034 | 0.0034 |
| wt vs. het | GOTERM_CC_DIRECT | Mitochondrial matrix | 10 | 8.26 | 4.01E-04 | 118 | 392 | 20624 | 4.46 | 0.0965 | 0.0338 | 0.0334 |
| wt vs. het | UP_KW_BIOLOGICAL_PROCESS | Translation regulation | 7 | 5.79 | 1.39E-04 | 70 | 130 | 11262 | 8.66 | 0.0087 | 0.0088 | 0.0088 |
| wt vs. het | UP_KW_DOMAIN | WD repeat | 7 | 5.79 | 0.0018 | 67 | 283 | 14504 | 5.35 | 0.0264 | 0.0134 | 0.0134 |
| wt vs. hom | GOTERM_CC_DIRECT | Mitochondrion | 23 | 19.01 | 2.12E-05 | 118 | 1458 | 20624 | 2.76 | 0.0054 | 0.0054 | 0.0054 |
| wt vs. hom | UP_KW_DOMAIN | Transit peptide | 12 | 9.92 | 1.93E-04 | 78 | 568 | 14504 | 3.93 | 0.0027 | 0.0027 | 0.0027 |
| wt vs. hom | UP_KW_DISEASE | Primary mitochondrial disease | 10 | 8.26 | 6.12E-06 | 34 | 197 | 4623 | 6.90 | 1.53E-04 | 1.53E-04 | 1.53E-04 |
| wt vs. hom | INTERPRO | Ubiquitin-conjugating enzyme, active site | 5 | 4.13 | 1.95E-05 | 116 | 27 | 19144 | 30.56 | 0.004666692 | 0.004677569 | 0.0047 |
| wt vs. hom | GOTERM_MF_DIRECT | Ubiquitin-like protein transferase activity | 5 | 4.13 | 3.13E-05 | 116 | 30 | 18945 | 27.22 | 0.008310655 | 0.00834525 | 0.0083 |
| wt vs. hom | GOTERM_BP_DIRECT | Protein modification by small protein conjugation | 5 | 4.13 | 3.91E-05 | 114 | 33 | 19414 | 25.80 | 0.030975603 | 0.031464875 | 0.0315 |
| wt vs. hom | GOTERM_MF_DIRECT | Ubiquitin conjugating enzyme activity | 5 | 4.13 | 6.53E-05 | 116 | 36 | 18945 | 22.68 | 0.01729033 | 0.008720492 | 0.0087 |
| wt vs. hom | INTERPRO | Ubiquitin-conjugating enzyme, E2 | 5 | 4.13 | 1.39E-04 | 116 | 44 | 19144 | 18.75 | 0.032893141 | 0.016721977 | 0.0167 |
| wt vs. hom | INTERPRO | Ubiquitin-conjugating enzyme/RWD-like | 5 | 4.13 | 4.36E-04 | 116 | 59 | 19144 | 13.99 | 0.099358277 | 0.034874978 | 0.0349 |
| wt vs. hom | UP_KW_MOLECULAR_FUNCTION | Ribosomal protein | 7 | 5.79 | 0.0013 | 75 | 193 | 11749 | 5.68 | 0.045619813 | 0.046662036 | 0.0467 |

**Table S9 Gene ontology (GO) terms and statistics for proteins with nominal differential expression in neurites of ZNF804A$^{+/-}$ and ZNF804A$^{-/-}$ neurites.**

#### Translational Machinery

##### RBMXL1

**Neurites**

| | Df | Sum sq | Mean sq | F value | Pr(>F) | Significant |
|---|---|---|---|---|---|---|
| Genotype | 2 | 1.876 | 0.9381 | 5.204 | 0.02 | * |
| Residuals | 14 | 2.524 | 0.1803 | | | |

TukeyHSD post-hoc tests:

$Genotype

| | diff | lwr | upr | p.adj | Significant |
|---|---|---|---|---|---|
| hom-het | 0.5204 | -0.1525 | 1.193324 | 0.1429 | ns |
| wt-het | -0.257 | -0.9303 | 0.415473 | 0.5881 | ns |
| wt-hom | -0.778 | -1.4194 | -0.13626 | 0.0174 | * |

**Somata**

| | Df | Sum sq | Mean | F value | Pr(>F) | Significant |
|---|---|---|---|---|---|---|
| Genotype | 2 | 1.249 | 0.6244 | 1.602 | 0.242 | ns |
| Residuals | 12 | 4.677 | 0.3897 | | | |

##### RPLP1

**Neurites**

| | Df | Sum sq | Mean sq | F value | Pr(>F) | Significant |
|---|---|---|---|---|---|---|
| Genotype | 2 | 25.5 | 12.752 | 6.036 | 0.013 | * |
| Residuals | 14 | 29.58 | 2.113 | | | |

TukeyHSD post-hoc tests:

$Genotype

| | diff | lwr | upr | p.adj | Significant |
|---|---|---|---|---|---|
| hom-het | -0.299 | -2.6022 | 2.004973 | 0.9388 | ns |
| wt-het | -2.714 | -5.0172 | -0.41007 | 0.0207 | * |
| wt-hom | -2.415 | -4.6114 | -0.21867 | 0.0307 | * |

**Somata**

| | Df | Sum sq | Mean | F value | Pr(>F) | Significant |
|---|---|---|---|---|---|---|
| Genotype | 2 | 4.12 | 2.062 | 0.369 | 0.703 | ns |
| Residuals | 8 | 44.71 | 5.588 | | | |

##### RPL31

**Neurites**

| | Df | Sum sq | Mean sq | F value | Pr(>F) | Significant |
|---|---|---|---|---|---|---|
| Genotype | 2 | 0.1138 | 0.05692 | 0.441 | 0.652 | ns |
| Residuals | 14 | 1.8072 | 0.12909 | | | |

**Somata**

| | Df | Sum sq | Mean sq | F value | Pr(>F) | Significant |
|---|---|---|---|---|---|---|
| Genotype | 2 | 7.826 | 3.913 | 5.837 | 0.0143 | * |
| Residuals | 14 | 9.385 | 0.67 | | | |

TukeyHSD post-hoc tests:

$Genotype

| | diff | lwr | upr | p.adj | Significant |
|---|---|---|---|---|---|
| hom-het | 1.433 | 0.13559 | 2.73076 | 0.0299 | * |
| wt-het | 1.539 | 0.24098 | 2.83615 | 0.0199 | * |
| wt-hom | 0.105 | -1.1318 | 1.34259 | 0.973 | ns |

##### RPS12

**Neurites**

| | Df | Sum sq | Mean sq | F value | Pr(>F) | Significant |
|---|---|---|---|---|---|---|
| Genotype | 2 | 0.256 | 0.1282 | 0.52 | 0.606 | ns |
| Residuals | 14 | 3.453 | 0.2466 | | | |

**Somata**

| | Df | Sum sq | Mean sq | F value | Pr(>F) | Significant |
|---|---|---|---|---|---|---|
| Genotype | 2 | 1.182 | 0.5912 | 5.291 | 0.0225 | * |
| Residuals | 12 | 1.341 | 0.1117 | | | |

TukeyHSD post-hoc tests:

$Genotype

| | diff | lwr | upr | p.adj | Significant |
|---|---|---|---|---|---|
| hom-het | 0.602 | 0.02683 | 1.17812 | 0.0401 | * |
| wt-het | 0.667 | 0.06884 | 1.2653 | 0.029 | * |
| wt-hom | 0.065 | -0.4754 | 0.6046 | 0.9457 | ns |

##### RPS15

**Neurites**

| | Df | Sum sq | Mean sq | F value | Pr(>F) | Significant |
|---|---|---|---|---|---|---|
| Genotype | 2 | 1.063 | 0.5313 | 0.679 | 0.523 | ns |
| Residuals | 14 | 10.958 | 0.7827 | | | |

**Somata**

| | Df | Sum sq | Mean sq | F value | Pr(>F) | Significant |
|---|---|---|---|---|---|---|
| Genotype | 2 | 7.191 | 3.595 | 4.349 | 0.038 | * |
| Residuals | 12 | 9.92 | 0.827 | | | |

TukeyHSD post-hoc tests:

$Genotype

| | diff | lwr | upr | p.adj | Significant |
|---|---|---|---|---|---|
| hom-het | 1.517 | -0.0486 | 3.08289 | 0.0578 | ns |
| wt-het | 1.617 | -0.0098 | 3.24456 | 0.0514 | ns |
| wt-hom | 0.1 | -1.3686 | 1.56906 | 0.9819 | ns |

##### RPS20

**Neurites**

| | Df | Sum sq | Mean sq | F value | Pr(>F) | Significant |
|---|---|---|---|---|---|---|
| Genotype | 2 | 0.032 | 0.0162 | 0.037 | 0.963 | ns |
| Residuals | 14 | 6.095 | 0.4354 | | | |

**Somata**

| | Df | Sum sq | Mean sq | F value | Pr(>F) | Significant |
|---|---|---|---|---|---|---|
| Genotype | 2 | 0.8921 | 0.446 | 5.045 | 0.0257 | * |
| Residuals | 12 | 1.0609 | 0.0884 | | | |

TukeyHSD post-hoc tests:

$Genotype

| | diff | lwr | upr | p.adj | Significant |
|---|---|---|---|---|---|
| hom-het | 0.36 | -0.1519 | 0.87215 | 0.1878 | ns |
| wt-het | 0.634 | 0.10137 | 1.16565 | 0.0202 | * |
| wt-hom | 0.273 | -0.2069 | 0.75375 | 0.3172 | ns |

| RPS21 | | | | | | |
|---|---|---|---|---|---|---|
| | Df | Sum sq | Mean sq | F value | Pr(>F) | Significant |
| Genotype | 2 | 4.47 | 2.2351 | 2.608 | 0.112 | ns |
| Residuals | 13 | 11.14 | 0.8572 | | | |

| RPS21 | | | | | | |
|---|---|---|---|---|---|---|
| | Df | Sum sq | Mean sq | F value | Pr(>F) | Significant |
| Genotype | 2 | 27.635 | 13.817 | 25.52 | 4.76E-05 | *** |
| Residuals | 12 | 6.497 | 0.541 | | | |
| TukeyHSD post-hoc tests: | | | | | | |
| $Genotype | | | | | | |
| | diff | lwr | upr | p.adj | Significant | |
| hom-het | 3.288 | 2.02101 | 4.55519 | 4E-05 | *** | |
| wt-het | 2.684 | 1.3674 | 4.001 | 0.0004 | *** | |
| wt-hom | -0.6 | -1.7925 | 0.58474 | 0.3935 | ns | |

| RPS28 | | | | | | |
|---|---|---|---|---|---|---|
| | Df | Sum sq | Mean sq | F value | Pr(>F) | Significant |
| Genotype | 2 | 1.006 | 0.503 | 0.34 | 0.718 | ns |
| Residuals | 14 | 20.717 | 1.48 | | | |

| RPS28 | | | | | | |
|---|---|---|---|---|---|---|
| | Df | Sum sq | Mean sq | F value | Pr(>F) | Significant |
| Genotype | 2 | 5.139 | 2.57 | 4.341 | 0.0382 | * |
| Residuals | 12 | 7.104 | 0.592 | | | |
| TukeyHSD post-hoc tests: | | | | | | |
| $Genotype | | | | | | |
| | diff | lwr | upr | p.adj | Significant | |
| hom-het | 1.212 | -0.1133 | 2.53667 | 0.0744 | ns | |
| wt-het | 1.423 | 0.04625 | 2.80016 | 0.0427 | * | |
| wt-hom | 0.212 | -1.0314 | 1.45445 | 0.8936 | ns | |

| SNU13 | | | | | | |
|---|---|---|---|---|---|---|
| | Df | Sum sq | Mean sq | F value | Pr(>F) | Significant |
| Genotype | 2 | 0.052 | 0.02579 | 0.104 | 0.902 | ns |
| Residuals | 14 | 3.477 | 0.24839 | | | |

| SNU13 | | | | | | |
|---|---|---|---|---|---|---|
| | Df | Sum sq | Mean sq | F value | Pr(>F) | Significant |
| Genotype | 2 | 1.2904 | 0.6452 | 9.704 | 0.00311 | ** |
| Residuals | 12 | 0.7979 | 0.0665 | | | |
| TukeyHSD post-hoc tests: | | | | | | |
| $Genotype | | | | | | |
| | diff | lwr | upr | p.adj | Significant | |
| hom-het | 0.58 | 0.13557 | 1.02367 | 0.0117 | * | |
| wt-het | 0.729 | 0.26739 | 1.19034 | 0.0032 | ** | |
| wt-hom | 0.149 | -0.2673 | 0.56581 | 0.6171 | ns | |

**Translation Regulation**

Neurites

| CNOT11 | | | | | | |
|---|---|---|---|---|---|---|
| | Df | Sum sq | Mean sq | F value | Pr(>F) | Significant |
| Genotype | 2 | 1.217 | 0.6084 | 1.544 | 0.253 | ns |
| Residuals | 12 | 4.729 | 0.394 | | | |

| NCBP2 | | | | | | |
|---|---|---|---|---|---|---|
| | Df | Sum sq | Mean sq | F value | Pr(>F) | Significant |
| Genotype | 2 | 2.811 | 1.4055 | 5.39 | 0.018 | * |
| Residuals | 14 | 3.651 | 0.2608 | | | |
| TukeyHSD post-hoc tests: | | | | | | |
| $Genotype | | | | | | |
| | diff | lwr | upr | p.adj | Significant | |
| hom-het | -0.265 | -1.0745 | 0.544076 | 0.6746 | ns | |
| wt-het | -0.966 | -1.7753 | -0.15673 | 0.0192 | * | |
| wt-hom | -0.701 | -1.4724 | 0.070816 | 0.0775 | ns | |

| RACK1 | | | | | | |
|---|---|---|---|---|---|---|
| | Df | Sum sq | Mean sq | F value | Pr(>F) | Significant |
| Genotype | 2 | 1.529 | 0.7644 | 7.665 | 0.006 | ** |
| Residuals | 14 | 1.396 | 0.0997 | | | |
| TukeyHSD post-hoc tests: | | | | | | |
| $Genotype | | | | | | |
| | diff | lwr | upr | p.adj | Significant | |
| hom-het | -0.083 | -0.5836 | 0.417366 | 0.9018 | ns | |
| wt-het | -0.669 | -1.1695 | -0.16851 | 0.0093 | ** | |
| wt-hom | -0.586 | -1.0631 | -0.10869 | 0.0161 | * | |

Somata

| CNOT11 | | | | | | |
|---|---|---|---|---|---|---|
| | Df | Sum sq | Mean | F value | Pr(>F) | Significant |
| Genotype | 2 | 2.54 | 1.268 | 0.081 | 0.923 | ns |
| Residuals | 9 | 141.53 | 15.726 | | | |

| NCBP2 | | | | | | |
|---|---|---|---|---|---|---|
| | Df | Sum sq | Mean | F value | Pr(>F) | Significant |
| Genotype | 2 | 0.8003 | 0.4001 | 2.705 | 0.107 | ns |
| Residuals | 12 | 1.7748 | 0.1479 | | | |

| RACK1 | | | | | | |
|---|---|---|---|---|---|---|
| | Df | Sum sq | Mean sq | F value | Pr(>F) | Significant |
| Genotype | 2 | 0.7745 | 0.3873 | 1.964 | 0.183 | ns |
| Residuals | 12 | 2.3662 | 0.1972 | | | |

| TARPB2 | | | | | | | TARPB2 | | | | | | |
|---|---|---|---|---|---|---|---|---|---|---|---|---|---|
| | Df | Sum sq | Mean sq | F value | Pr(>F) | Significant | | Df | Sum sq | Mean sq | F value | Pr(>F) | Significant |
| Genotype | 2 | 1.187 | 0.5936 | 4.532 | 0.03 | * | Genotype | 2 | 0.221 | 0.1107 | 0.308 | 0.74 | ns |
| Residuals | 14 | 1.834 | 0.131 | | | | Residuals | 12 | 4.305 | 0.3587 | | | |
| TukeyHSD post-hoc tests: | | | | | | | | | | | | | |
| $Genotype | | | | | | | | | | | | | |
| | diff | lwr | upr | p.adj | Significant | | | | | | | | |
| hom-het | -0.269 | -0.8423 | 0.304807 | 0.4578 | ns | | | | | | | | |
| wt-het | -0.652 | -1.2252 | -0.07806 | 0.0256 | * | | | | | | | | |
| wt-hom | -0.383 | -0.9297 | 0.164014 | 0.1953 | ns | | | | | | | | |

**Synapse**

| Neurites | | | | | | | Somata | | | | | | |
|---|---|---|---|---|---|---|---|---|---|---|---|---|---|
| VGLUT1 | | | | | | | VGLUT1 | | | | | | |
| | Df | Sum sq | Mean sq | F value | Pr(>F) | Significant | not enough protein detected in this compartment | | | | | | |
| Genotype | 2 | 0.767 | 0.3836 | 0.669 | 0.532 | ns | | | | | | | |
| Residuals | 11 | 6.306 | 0.5733 | | | | | | | | | | |
| BSN | | | | | | | BSN | | | | | | |
| | Df | Sum sq | Mean sq | F value | Pr(>F) | Significant | not enough protein detected in this compartment | | | | | | |
| Genotype | 2 | 1.49 | 0.7448 | 1.16 | 0.344 | ns | | | | | | | |
| Residuals | 13 | 8.347 | 0.6421 | | | | | | | | | | |
| SV2A | | | | | | | SV2A | | | | | | |
| | Df | Sum sq | Mean sq | F value | Pr(>F) | Significant | | Df | Sum sq | Mean sq | F value | Pr(>F) | Significant |
| Genotype | 2 | 0.078 | 0.03899 | 0.141 | 0.87 | ns | Genotype | 2 | 1.937 | 0.9687 | 0.982 | 0.403 | ns |
| Residuals | 14 | 3.869 | 0.27638 | | | | Residuals | 12 | 11.832 | 0.986 | | | |
| PSD95 | | | | | | | PSD95 | | | | | | |
| | Df | Sum sq | Mean sq | F value | Pr(>F) | Significant | | Df | Sum sq | Mean sq | F value | Pr(>F) | Significant |
| Genotype | 2 | 0.511 | 0.2554 | 0.538 | 0.571 | ns | Genotype | 2 | 0.903 | 0.4515 | 1.01 | 0.396 | ns |
| Residuals | 14 | 6.134 | 0.4381 | | | | Residuals | 11 | 4.918 | 0.4471 | | | |

**Table S10 Summary results tables and Tukey HSD post-hoc tests for one-way ANOVAs run for ribosomal, translation regulation, and synaptic proteins mis-localized in ZNF804A$^{+/-}$ and ZNF804A$^{-/-}$ neurons.** * p.adj. < 0.05

| Line | Clone | Diagnosis | Age | Sex | Reprogramming | Cohort |
|---|---|---|---|---|---|---|
| 014_CTM | 01 | Control | 18-30 | Male | CytoTune™ Sendai | LEAP / StemBANCC |
| | 02 | | | | | |
| | 03 | | | | | |
| M3_CTM | 36S | Control | 18-30 | Male | CytoTune™ Sendai | EU-AIMS |
| | 37S | | | | | |
| | 38S | | | | | |
| 127_CTM | 01 | Control | 50 - 60 | Male | CytoTune™ Sendai | EU-AIMS |
| | 02 | | | | | |
| | 04 | | | | | |

**Table S11 Relevant information about donor lines used for ZNF804A expression analysis in neuroprogenitor cells.**

| Medium | Reagent | Final Concentration |
| --- | --- | --- |
| N2 | DMEM (Sigma; D6421) | 1X |
| | N2 Supplement (Life Technologies; 17502-048) | |
| | Glutamax (Life Technologies; 35050-038) | |
| B27 | Neurobasal Medium (Life Technologies; 21103-049) | 1X |
| | B27 Supplement (Life Technologies; 17504-044) | |
| | Glutamax (Life Technologies; 35050-038) | |
| Neuralisation | B27:N2 (as above) | 1:1 mixture |
| | SB431542 (Cambridge Bioscience; ZRD-SB-50) | |
| | LDN193189 (Sigma; SML0559) | |
| NPC maintenance | B27:N2 (components as above) | 1:1 mixture |

**Table S12 Media compositions used in this study for neuroprogenitor cell differentiation.**

| Plate | Cell Density (per well) | Amount of Medium (per well) | Application |
| --- | --- | --- | --- |
| 6-well | 600,000 | 2ml | Transcriptomics & Western blot |
| 18mm coverslips | 50,000 | 1ml | ICC & confocal microscopy |
| 96-well | 10,000 or 15,000 | 100µl | SUnSET & ICC with high-content imaging |

**Table S13 Cell densities for each experimental set up used in this study.**

| Medium | Reagent | Final Concentration |
|---|---|---|
| D1 Neuralisation | StemFlex | 1X |
| | SB431542 (Cambridge Bioscience, ZRD-SB-50) | 10μM |
| | LDN193189 (Sigma, SML0559) | 0.1μM |
| | XAV939 (Sigma-Aldrich, X3004) | 2μM |
| | Doxycycline hyclate (Sigma, D9891) | 2μM |
| D2 Neuralisation | StemFlex:N2 (N2 as in Supplemental Table 2.1) | 1:1 mixture |
| | SB431542 (Cambridge Bioscience, ZRD-SB-50) | 10μM |
| | LDN193189 (Sigma, SML0559) | 0.1μM |
| | XAV939 (Sigma-Aldrich, X3004) | 2μM |
| | Doxycycline hyclate (Sigma, D9891) | 2μM |
| D3 Neuralisation | N2 (as as in Supplemental Table 2.1) | 1X |
| | SB431542 (Cambridge Bioscience, ZRD-SB-50) | 10μM |
| | LDN193189 (Sigma, SML0559) | 0.1μM |
| | XAV939 (Sigma-Aldrich, X3004) | 2μM |
| | Doxycycline hyclate (Sigma, D9891) | 2μM |
| Maturation | B27 (as as in Supplemental Table 2.1) | 1X |
| | BDNF (PeproTech, 450-02) | 10ng/ml |
| | GDNF (PeproTech, 450-10) | 10ng/ml |
| | DAPT (Santa Cruz Biotechnology, SC-201315A) | 10μM |
| | Doxycycline hyclate (Sigma, D9891) | 2μM |

**Table S14 Media used for differentiation protocol combining NGN2 forward programming with SMAD and Wnt inhibition to generate developing patterned forebrain neurons**

| sgRNA1 (Intron2): GTTGTGTTTACCTCTCGACA | | | | | | | |
|---|---|---|---|---|---|---|---|
| | On-target | Off-target | Off-targets for Potential Mismatches | | | | |
| | | | 0 | 1 | 2 | 3 | 4 |
| **IDT** | 65 | 72 | N/A | N/A | N/A | N/A | N/A |
| **CRISPOR** | 65 | 92 | 0 | 0 | 0 | 3 | 54 |

| sgRNA2 (Intron 3): CTACTATAGCCCAATTGGTA | | | | | | | |
|---|---|---|---|---|---|---|---|
| | On-target | Off-target | Off-targets for Potential Mismatches | | | | |
| | | | 0 | 1 | 2 | 3 | 4 |
| **IDT** | 55 | 78 | N/A | N/A | N/A | N/A | N/A |
| **CRISPOR** | 37 | 94 | 0 | 0 | 0 | 2 | 51 |

**Table S15 Single guide RNA (sgRNA) sequences used for PFuncGen including on- and off-target scores provided by IDT or CRISPOR.**

| Sample ID | Barcode Sequence | # Reads | Yield (Mbases) | Mean Quality Score | % Bases >= 30 |
|---|---|---|---|---|---|
| LSI02_61-RP1 | TTAAGCAG+GTCCAGGA | 51608880 | 15483 | 35.86 | 93.54 |
| LSI02_10-23-RP2 | AGCCTGGA+AGCAGACT | 64753607 | 19426 | 35.63 | 92.33 |
| LSI02_61-RP3 | CTTTATTC+CCGTTCCA | 75343202 | 22603 | 35.84 | 93.43 |
| LSI02_44-RP2 | TCACTAAC+AGATAGGG | 57188527 | 17157 | 35.78 | 93.11 |
| LSI02_4-RP2 | TAAGCGCA+TTCTACGG | 69371260 | 20811 | 35.76 | 93.01 |
| LSI02_42-13-RP1 | TTTGAGTC+CTATCCGG | 70197222 | 21059 | 35.82 | 93.31 |
| LSI02_4-RP1 | ACTTCTGC+CCGGGACT | 48283312 | 14485 | 35.81 | 93.3 |
| LSI02_20-RP3 | GCAATGGG+CTTGTACT | 45119939 | 13536 | 35.82 | 93.36 |
| LSI02_61-RP2 | CAACGGAA+AGTAGATG | 66104245 | 19831 | 35.82 | 93.32 |
| LSI02_10-23-RP1 | ATCAAATC+TCATGTAG | 90669053 | 27201 | 35.84 | 93.4 |
| LSI02_20-RP2 | CTATCGAA+ATCATAGT | 126492339 | 37948 | 35.83 | 93.42 |
| LSI02_4-RP3 | TCGATAAG+GGACAACG | 75588458 | 22677 | 35.82 | 93.35 |
| LSI02_44-RP3 | CTGGACAC+TCACAGTA | 72360832 | 21708 | 35.76 | 92.98 |
| LSI02_20-RP1 | AAATCCTC+GGCGGGTT | 47546113 | 14264 | 35.8 | 93.21 |
| LSI02_42-13-RP2 | GAGGACAG+GATTCGGC | 50591801 | 15178 | 35.83 | 93.4 |
| LSI02_42-13-RP3 | ACAACCAA+TTTGGACA | 38022986 | 11407 | 35.91 | 93.85 |
| LSI02_10-23-RP3 | CTGCCTTC+CCGAAGGG | 81840892 | 24552 | 35.81 | 93.24 |
| LSI02_44-RP1 | TACAGATG+CACGTTCG | 73997389 | 22199 | 35.8 | 93.2 |

**Table S16 RNA sequencing quality scores and number of unique reads for each sample.**

| Western Blotting | | | | |
|---|---|---|---|---|
| **Antibody** | **Species** | **Company** | **Cat. No.** | **Dilution** |
| *Primary antibodies* | | | | |
| GAPDH | mouse | ProteinTech | 60004-I | 1:10k |
| H3 | rabbit | abcam | ab1791 | 1:2000 |
| pan-PCDHG | mouse | abcam | ab187186 | 1:1000 |
| RPS10 | rabbit | abcam | ab151550 | 1:1000 |
| RPS6 | mouse | Cell Signalling | 2317S | 1:1000 |
| pRPS6 (Ser235/236) | rabbit | Cell Signalling | 4858S | 1:1000 |
| ZNF804A C2C3 | rabbit | GeneTex | GTX121178 | 1:200 |
| *Secondary antibodies* | | | | |
| IgG Secondary antibody, HRP conjugate | Goat anti-mouse | Invitrogen | 31430 | 1:10k |
| IgG Secondary antibody, HRP conjugate | Goat anti-rabbit | Invitrogen | 31460 | 1:10k |

| Immunocytochemistry | | | | |
|---|---|---|---|---|
| **Antibody** | **Species** | **Company** | **Cat. No.** | **Dilution** |
| *Primary antibodies* | | | | |
| Bassoon | rabbit | Cell Signalling | 6897S | 1:500 |
| CUX2 | rabbit | abcam | ab130395 | 1:500 |
| GluN1 | mouse | BioLegend | 818601 | 1:500 |
| Ki67 | rabbit | abcam | ab15580 | 1:500 |
| MAP2 | chicken | abcam | ab92434 | 1:1000 |
| Neurofilament (pan axonal) | mouse | BioLegend | 837904 | 1:500 |
| OCT3/4 | mouse | Santa Cruz Biotechnology | sc-5279 | 1:100 |
| PAX6 | rabbit | BioLegend | 901301 | 1:500 |
| PSD95 | mouse | BioLegend | 810401 | 1:500 |
| Puromycin | mouse | Millipore | MABE343 | 1:5000 |
| RPS6 | mouse | Cell Signalling | 2317S | 1:500 |
| SV2A | rabbit | abcam | ab32942 | 1:500 |
| TBR1 | rabbit | abcam | ab31940 | 1:200 |
| TUBB3 | chicken | Aves Labs | TUJ | 1:1000 |
| VGLUT1 | rabbit | Synaptic Systems | 135302 | 1:500 |
| *Secondary antibodies* | | | | |
| Alexa Fluor 488 | Goat anti-mouse | Invitrogen | A11001 | 1:750 |
| Alexa Fluor 568 | Goat anti-rabbit | Invitrogen | A11011 | 1:750 |
| Alexa Fluor 633 | Goat anti-chicken | Invitrogen | A21103 | 1:750 |

**Table S17 List of antibodies used in this study.**
